# The influence of microbial mats on travertine precipitation in active hydrothermal systems (Central Italy)

**DOI:** 10.1101/2020.07.29.226266

**Authors:** Giovanna Della Porta, Joachim Reitner

## Abstract

The study of hydrothermal travertines contributes to the understanding of the interaction between physico-chemical processes and the role played by microbial mats and biofilms in influencing carbonate precipitation. Three active travertine sites were investigated in Central Italy to identify the types of carbonate precipitates and the associated microbial mats at varying physico-chemical parameters. Carbonate precipitated fabrics at the decimetre- to millimetre-scale and microbial mats vary with decreasing water temperature: a) at high temperature (55-44°C) calcite or aragonite crystals precipitate on microbial mats of sulphide oxidizing, sulphate reducing and anoxygenic phototrophic bacteria forming filamentous streamer fabrics, b) at intermediate temperature (44-40°C), rafts, coated gas bubbles and dendrites are associated with *Spirulina* cyanobacteria and other filamentous and rod-shaped cyanobacteria, c) low temperature (34-33°C) laminated crusts and oncoids in a terraced slope system are associated with diverse Oscillatoriales and Nostocales filamentous cyanobacteria, sparse *Spirulina* and diatoms. At the microscale, carbonate precipitates are similar in the three sites consisting of prismatic calcite (40-100 μm long, 20-40 μm wide) or acicular aragonite crystals organized in radial spherulites, overlying or embedded within biofilm EPS (Extracellular Polymeric Substances). Microsparite and sparite crystal size decreases with decreasing temperature and clotted peloidal micrite dominates at temperatures < 40°C, also encrusting filamentous microbes. Carbonates are associated with gypsum and Ca-phosphate crystals; EPS elemental composition is enriched in Si, Al, Mg, Ca, P, S and authigenic aluminium-silicates form aggregates on EPS.

This study confirms that microbial communities in hydrothermal travertine settings vary as a function of temperature. Carbonate precipitate types at the microscale do not vary considerably, despite different microbial communities suggesting that travertine precipitation, driven by CO_2_ degassing, is influenced by biofilm EPS acting as template for crystal nucleation (EPS-mediated mineralization) and affecting the fabric types, independently from specific microbial metabolism.

## 1. Introduction

Travertines are terrestrial carbonates precipitated in hydrothermal settings (Pedley, 1990; Capezzuoli et al., 2014), which thrive with microbial mats consisting of a wide spectrum of thermophilic Archaea and Bacteria, including sulphide oxidizing bacteria, sulphate reducing bacteria, anoxygenic phototrophs and oxygenic photosynthetic cyanobacteria, other prokaryotes and eukaryotic algae (Farmer, 2000; Fouke et al., 2003; Pentecost, 2003, 2005; Konhauser, 2007; Della Porta, 2015). Sedimentological and microbiological investigations on present-day active hydrothermal systems are fundamental because these unusual aquatic terrestrial environments store important palaeobiological information due to their high rates of mineralization (e.g., Farmer, 2000; Campbell et al., 2015).

The study of microbial biofilms and microbial mats in warm to hot hydrothermal vent environments and their interaction with siliceous and carbonate mineral precipitation is linked to the understanding of early life forms on Earth and could also be important for the search of biosignatures on putative habitable planets and moons (cf. Cady and Farmer, 1996; Farmer, 1998; Reysenbach et al., 1999; Westall et al., 2000; Reysenbach and Cady, 2001; Reysenbach and Shock, 2002; Rothshield and Mancinelli, 2001; Cady and Noffke, 2009; Ruff and Farmer, 2016; Cady et al., 2018; Reinhardt et al., 2019; Sanchez-Garcia et al. 2019; Franchi and Frisia, 2020). Geomicrobiological research on the origin of life has recently shifted from very high temperature submarine Black Smokers to terrestrial thermal springs, because the relatively lower temperatures of terrestrial geothermal sites facilitate the preservation of organic molecules (Des Marais and Walter, 2019). Despite the abundance of Cenozoic terrestrial thermal spring settings, mainly depositing siliceous sinter and/or carbonate travertines (Pentecost, 2005; Jones and Renaut, 2010, 2011; Capezzuoli et al., 2014), these deposits are scarce in the fossil record (Des Marais and Walter, 2019). Few examples are known from the Jurassic in Argentina with travertine and siliceous sinter (Guido and Campbell, 2011), the Devonian hydrothermal siliceous deposits from the Drummond Basin (Queensland, Australia; White et al., 1989) and the lower Devonian Rhynie Chert in Scotland, which exhibits excellently preserved plant, animal and prokaryote fossils (Rice et al., 2002; Edwards et al., 2017). In the Proterozoic and Archaean Eon, identified terrestrial thermal spring deposits are rare (Des Marais and Walter, 2019). The oldest reported terrestrial hydrothermal travertine deposits, attributed to deep-sourced CO_2_ fluids at high temperature, are hosted in the Kuetsjärvi Sedimentary Formation of the Pechenga Greenstone Belt in the Fennoscandian Shield, and are dated to the Palaeoproterozoic, at approximately 2.2 Ga (Melezhik and Fallick, 2001; Brasier, 2011; Salminen et al., 2014). These travertines consist of dolomite and are associated with dolomitic stromatolites formed at the shorelines of a rift-related playa lake system (Melezhik and Fallick, 2001). It is suggested that these carbonates might have formed through similar processes to modern hydrothermal springs but they do not contain traces of organic carbon (Salminen et al., 2014). The oldest siliceous deposits attributed to terrestrial thermal springs are known from the ca. 3.5 Ga Dresser Formation (Pilbara, Western Australia), which were interpreted as putative geyser environments (Djokic et al., 2017). Marine hydrothermal settings in the early Archaean were, however, common. Black Smoker environments were first reported from the Sulphur Spring locality, also in Pilbara, with an approximate age of 3.2 Ga (Buick et al., 2002). Hydrothermal pumping was a common feature and played a central role in carbon sequestration (cf. Duda et al., 2018), forming several kilometres long Black Chert veins within lower Archaean oceanic crust formations (Green Stone Belts; van Kranendonk, 2006).

Widespread active terrestrial thermal springs and fossil travertines in Central Italy represent key localities to investigate carbonate precipitation under thermal water conditions. In subaerial hydrothermal systems, supersaturation for carbonate minerals is primarily achieved by CO_2_ evasion from thermal water issuing out of a vent and flowing along the topographic profile (Pentecost, 2005). The precipitation of carbonate minerals might be primarily driven by physico-chemical processes (Pentecost and Coletta, 2007) or the diverse travertine facies might result from a combination of inorganic, biologically induced and influenced processes in association with microbial mats (Chafetz and Folk, 1984; Folk, 1994; Guo and Riding, 1994; Pentecost, 1995a; Guo et al., 1996; Chafetz and Guidry, 1999; Fouke et al., 2000; Riding, 2008; Rainey and Jones, 2009; Fouke, 2011; Gandin and Capezzuoli, 2014; Jones and Peng, 2014a; Della Porta, 2015; Erthal et al., 2017). There is, however, not a full understanding of the geomicrobiological processes active in these carbonate-dominated hydrothermal systems including the role played by diverse microorganisms, their metabolic pathways, biofilm organic substrates and the physico-chemical and biochemical factors enhancing or inhibiting the precipitation of carbonate and other minerals.

Numerous studies on present-day marine and terrestrial settings, from the geological record and laboratory experiments have demonstrated that carbonate mineral precipitation associated with microbial mats can take place in aquatic environments through different mechanisms as a function of environmental conditions, microbial communities and nature of organic substrates. Carbonate precipitation associated with microbial mats and organic substrates in aqueous fluids, under favourable physico-chemical conditions, can be both the result of metabolic activity of alive microbes (Chafetz and Buczynski, 1992; Castanier et al., 1999; Knorre and Krumbein, 2000; biologically induced mineralization after Lowenstam and Weiner, 1989; Dupraz et al., 2009) and/or the result of mineralization of non living organic substrates with acidic macromolecules able to bind Ca^2+^ and Mg^2+^ ions (organomineralization after Reitner, 1993; Défarge and Trichet, 1995; Trichet and Défarge, 1995; Reitner et al., 1995ab; Défarge et al., 1996; Neuweiler et al., 1999; Reitner et al., 2000, 2001; Défarge et al., 2009; organomineralization *sensu strictu* and biologically influenced mineralization after Dupraz et al., 2009; organic-compound catalyzed mineralization after Franchi and Frisia, 2020). Organomineralization has been identified in association with various organic substrates such as microbial biofilm EPS (Extracellular Polymeric Substances), post-mortem encrustation of bacterial cells, sponge tissues or even abiotic organic compounds (Reitner, 1993; Reitner, 2004; Défarge et al., 2009).

Among the microbial metabolic pathways that appear to play a key role in carbonate precipitation by modifying the microenvironment, driving increased alkalinity and carbonate supersaturation, the most investigated are: a) autotrophic oxygenic cyanobacteria photosynthesis (Pentecost and Riding, 1986; Robbins and Blackwelder, 1992; Merz, 1992; Castanier et al., 1999; Merz-Preiß and Riding, 1999; Merz-Preiß, 2000; Golubic et al., 2000; Bissett et al., 2008; Obst et al., 2009; Plée et al., 2010; Eymard et al., 2020), with eventually carbonate encrustation of microbial cells in settings with low dissolved inorganic carbon and high in Ca^2+^ (Merz, 1992; Défarge et al., 1996; Arp et al., 2001, 2002, 2010; Kamennaya et al., 2012); and b) heterotrophic bacteria ammonification of amino-acids and sulphate reduction (Castanier et al., 1999; Knorre and Krumbein, 2000; Visscher et al., 2000; Dupraz et al., 2004; Dupraz and Visscher, 2005; Andres et al., 2006; Baumgartner et al., 2006; Dupraz et al., 2009).

Microbial mat and biofilm exopolymers (EPS) play a fundamental controlling role in the precipitation of carbonate and other minerals (cf. Défarge and Trichet, 1995; Reitner et al., 1995a; Wingender et al., 1999; Westall et al., 2000; Decho, 2010; Decho and Gutierrez, 2017). All these studies demonstrate that microbial cells and eukaryote algae in marine and terrestrial environments can secrete diverse arrays of polymers (EPS). Dominant macromolecules are often acidic polysaccharides and (glycol-) proteins, including lectins, which are rich in negatively charged carboxylic- and sulphate groups, which may bond divalent cations. EPS facilitate also attachment to surfaces that lead to the formation of microbial mat and biofilm communities, stabilizing cells, protecting from physical stresses, such as changes in salinity, temperature, UV irradiation and desiccation. One main function of EPS is the inhibition of fast precipitation of various minerals, mainly carbonates, to avoid the blockade of ionic exchange between cells and ambient water (Défarge and Trichet, 1995; Reitner et al., 1995a; Arp et al., 1999, 2001, 2003). The EPS acidic macromolecules have a matrix and template function of mineralisation and inhibit precipitation by providing bonding of divalent cations, such as Ca^2+^, Sr^2+^, Mg^2+^ (Arp et al., 1999, 2012; Ionescu et al., 2014). Therefore, EPS can either promote calcium carbonate precipitation acting as a template for crystal nucleation (EPS-mediated mineralization; Reitner, 1993; Reitner et al., 1995a) or inhibit precipitation by binding free calcium ions (Arp et al., 1999, 2001, 2003; Ionescu et al., 2014). Crystal nucleation is promoted by highly ordered acidic groups at defined distances that correspond to the crystal lattice, whereas disordered organic matrices as EPS groups inhibit precipitation (Arp et al., 2001). Nucleation of CaCO_3_ only occurs at acidic groups, which are suitably arranged mainly by accident, after a sufficient diffusive Ca^2+^ supply surpasses the complexation capacity of EPS. The binding capabilities of acidic exopolymers and their inhibiting effect increase with pH, therefore this effect is stronger in alkaline settings and in the phototrophic zone (Arp et al., 1999, 2003; Ionescu et al., 2014). The EPS inhibiting capability is surpassed by degradation that releases Ca^2+^ increasing calcium carbonate supersaturation and enhancing mineral precipitation (Reitner et al., 1996, 1997; Arp et al., 1999; Braissant et al., 2007, 2009; Ionescu et al., 2014). Various studies suggest that biofilm EPS degradation through bacterial sulphate reduction is pivotal to carbonate precipitation in present-day microbialites (Dupraz et al., 2004; Dupraz and Visscher, 2005; Baumgartner et al., 2006; Dupraz et al., 2009; Glunk et al., 2011). Hence, carbonate precipitation can be induced by living microorganisms and their metabolic pathways, but it can also occur without the contribution of microbial metabolism, mediated by organic compounds independently from the living organisms from which these compounds may derive (Défarge et al., 2009). These two mechanisms must have been both active in the geologic record together with physico-chemically driven abiotic mineralization (Défarge et al., 1996; Riding, 2000, 2008). The first probable EPS-mediated mineralisation deposits are the Strelley Pool stromatolites (ca. 3.35 Ga) from Pilbara Craton, Western Australia (e.g. Allwood et al., 2006; Viehmann et al., 2020).

More investigation is required to unravel the microbial community composition, the spatial distribution of carbonate precipitates within the microbial mats and the controls exerted by microbial biofilms on carbonate mineral precipitation and fabric types in travertine hydrothermal systems, where water is supersaturated with respect to carbonate due to physico-chemical processes. By comparing three active sites in Central Italy with different water chemistry and temperature (from 33 to 55°C), this study aims to improve the understanding on the interaction among water physico-chemical parameters, microbial community composition, biofilm EPS and the specific carbonate precipitated fabrics and associated minerals.

## 2. Geological background of studied travertines

From the Neogene, Central Italy is the site of widespread deposition of hydrothermal travertine deposits, in particular during the Pleistocene and Holocene times (e.g., Chafetz and Folk, 1984; Guo and Riding, 1992, 1994, 1998, 1999; Pentecost, 1995ab; Minissale et al., 2002ab; Minissale, 2004; Faccenna et al., 2008; Brogi and Capezzuoli, 2009; Capezzuoli et al., 2014; Della Porta, 2015; Croci et al., 2016; Brogi et al., 2016; Della Porta et al., 2017ab; Erthal et al., 2017). Travertines in Central Italy are associated with Neogene and Quaternary magmatic activity and extensional tectonics superimposed on the Apennine orogen, consisting of Mesozoic and Cenozoic thrust sheets (Figure 1). The western side of the Apennine fold-and-thrust belt was affected by extensional and strike-slip tectonics from Miocene time due to back-arc related extension of the Tyrrhenian Sea in the West, while in the East the Adriatic plate was in subduction westwards and the Apennine thrusts propagated eastwards (Malinverno and Ryan, 1986; Doglioni, 1991; Carminati and Doglioni, 2012). Due to Neogene tectonics, several sedimentary basins filled by Miocene to Quaternary marine and terrestrial sedimentary successions developed with preferential orientation NW-SE (Faccenna et al., 2008; Carminati and Doglioni, 2012). In these extensional and strike-slip basins, hydrothermal activity with Ca-SO_4_-HCO_3_ water composition (Minissale, 2004) and related travertine deposition are common features due to: a) Pliocene-Holocene intrusive and effusive rocks with up to present-day active volcanic activity, b) active faults acting as fluid conduits, c) humid climate and mountain relief driving atmospheric precipitation, d) substrate rocks consisting of hundreds of metres thick Mesozoic carbonate succession providing calcium and carbonate ions and H_2_S derived from Triassic evaporites (Minissale, 2004).

**Figure 1.**
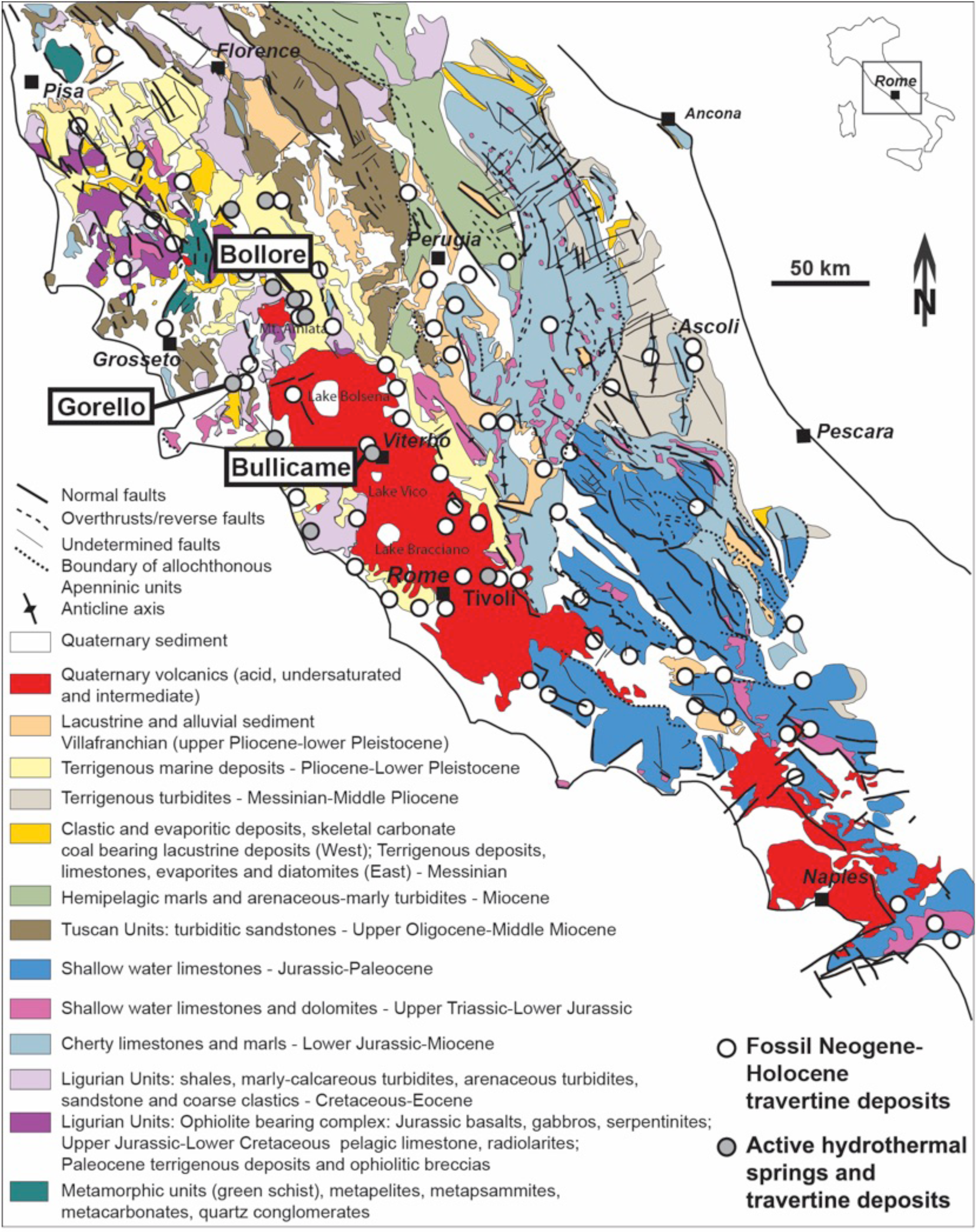
Generalized geological map of Central Italy (modified after Bigi et al., 1990; redrafted after Della Porta, 2015) with sites of active and fossil hydrothermal springs as reported by Minissale (2004). The three active hydrothermal travertine sites sampled in this study are indicated: Bullicame (near the town of Viterbo), Bollore (near the village of Bagni San Filippo and NE of the volcanic complex of Monte Amiata) and Gorello Waterfall (near the village of Saturnia).

At present, travertine deposition takes place at numerous active hydrothermal vents (Figure 1) where water, with temperatures varying from 20°C to 68°C (Minissale, 2004), emerges at the surface and degasses CO_2_ while outflowing away from the vent, following the local topographic gradient. The origin of CO_2_ in the central Italian hydrothermal systems is debated (Minissale, 2004) and has been attributed to: a) hydrolysis of regionally extensive Mesozoic limestones; b) metamorphism of limestones within the Palaeozoic basement; and c) mantle derived CO_2_. Based on stable isotope data of thermal water and travertine deposits, the main source of CO_2_ is related to decarbonation of limestones triggered by the presence of shallow mantle intrusions in the crust (Minissale, 2004).

The three selected sites of active hydrothermal travertine deposition (Figure 1) are Bullicame (Latium region), the Bollore vent and Gorello Waterfall in Southern Tuscany. The Bullicame hydrothermal vent is located near the town of Viterbo; it has water temperature of 55-57°C and pH 6.5 at the vent (Pentecost, 1995a; Piscopo et al., 2006). The Bollore (Bagni San Filippo village) vent issues water with temperatures of nearly 47-52°C and pH 6.7-6.5 (Pentecost, 1995a; Minissale, 2004; Brogi and Fabbrini, 2009). The Gorello Waterfall travertine system, close to Saturnia village, consists of a cascade and travertine apron deposits formed by thermal water with temperature of 37°C and pH 6.3 at the vent (Minissale, 2004; Ronchi and Cruciani, 2015).

## 3. Methods

Travertine and water sampling was performed in Central Italy in three sites (Figure 1) with active hydrothermal vents and travertine precipitation during the summer month of July with high air temperature and limited atmospheric precipitation. The three selected travertine settings are: a) Bullicame, near the town of Viterbo (N 42° 25’ 13.53”, E 12° 04’ 22.58”, elevation 293 m a.s.l.); b) Bollore, close to the village of Bagni San Filippo north-east of the Mount Amiata volcanic complex (N 42° 55’ 38”, E 11° 41’ 41.5”, elevation 588 m a.s.l.); c) Gorello Waterfall, near the village of Saturnia (N 42° 38’ 53.30”, E 11° 30’ 45.45”, elevation 138 m a.s.l.). Water temperature, pH and total alkalinity were measured in the field with a hand-held pH-metre and titrator, Mortimer solution and sulphuric acid cartridges. Data relative to water analyses are displayed in Table 1.

**Table 1.** Analyses of water parameters (temperature, pH, alkalinity) in the field from proximal to distal.

| Sample locality | Temperature<br>(°C) | pH | Alkalinity<br>(meq/l) |
| --- | --- | --- | --- |
| **Bullicame (Viterbo)** |  |  |  |
| **The vent is fenced; first sampling site in the channel 12.5 m from the centre of the vent pool** |  |  |  |
| Channel 12.5 m from centre of the vent | 55 | 6.7 | 15.6 |
| 15.5 m | 54.6 | 6.7 |  |
| 17.5 m | 54.3 | 6.7 |  |
| 21.5 m | 53.9 | 6.8 | 15.76 |
| 26.5 m | 53.4 | 6.8 |  |
| 30.5 m | 53.1 | 6.9 |  |
| 35.5 m | 52.5 | 6.9 |  |
| 41.5 m | 52 | 7.1 |  |
| 48.5 m | 50 | 7.3 |  |
| 55.5 m | 50.5 | 7.2 |  |
| 60.5 m | 50 | 7.3 | 14.52 |
| 73.5 m In water at centre of channel 61 m | 49 | 7.4 |  |
| 73.5 m In green microbial mat at rim of channel | 48.4 | 7.3 |  |
| 83.5 m | 48.2 | 7.4 |  |
| **Bollore (Bagni San Filippo) main spring** |  |  |  |
| Vent A of the main spring | 46.5 | 6.5 | 31.65 |
| Vent B of main spring | 46.1 | 6.6 |  |
| Vent C of main spring | 49.5 | 6.5 |  |
| Channel 0.4 m from vent C | 49 | 6.6 |  |
| 3.10 m | 47.3 | 6.8 |  |
| 8.10 m | 44 | 7.1 |  |
| 14 m | 41.4 | 7.3 | 21.55 |
| 20.3 m | 38.2 | 7.4 |  |
| 24.5 m | 36.4 | 7.5 |  |
| **Bollore second spring** |  |  |  |
| Vent of second spring | 49.3 | 6.5 |  |
| Channel 2 m from vent | 48.4 | 6.5 |  |
| 5 m | 40.6 | 7 |  |
| **Gorello Waterfalls (Saturnia)** |  |  |  |
| At waterfall | 33.8 | 7.8 |  |
| 1 m from waterfall in pools of terraced slope | 33.4 | 7.8 |  |
| 2 m | 33.6 | 7.8 |  |
| 3 m | 33.6 | 7.9 |  |
| 4 m | 33.7 | 7.8 |  |
| 5 m | 33 | 7.9 | 9.07; 9.04 |
| 6 m | 33.5 | 7.9 |  |
| 7 m | 33.6 | 7.9 |  |
| 8 m | 33.5 | 7.9 |  |

Sixteen samples of precipitated carbonate and the adherent microbial mats were collected in falcon tubes of 50 ml and fixed with formaldehyde 4% in 1x PBS (Phosphate Buffer Solution) and in glutaraldehyde with concentration of 4% in 1x PBS. Samples were stored in a fridge at 4°C during transportation from the sampling site to the Geobiology Department of the Georg-August-University Göttingen (Germany) and processed within 4 days from collection. The fixative was removed and the samples were dehydrated in graded ethanol series and then embedded in a resin (LR White, London Resin Company, UK). Once hardened, 70 thin sections were prepared for petrographic analysis with a Zeiss AX10 microscope equipped with a digital camera. Prior to resin embedding, subsamples were stained with the fluorochrome calcein, tetracycline, DAPI and toluidine blue. Calcein and/or tetracycline allow detecting Ca-binding areas. Calcein, the most important Ca^2+^ detecting fluorochrome, can be transported through the cell membrane into living cells. The acetoxymethyl-group of calcein is able to chelate calcium. This is then able to complex calcium ions within the cell, which results in a strong green fluorescence. DAPI is a good DNA fluorochrome and toluidine blue is a basic thiazine metachromatic dye with high affinity for acidic tissue components. Eight subsamples of prevalent organic soft biofilm were prepared for embedding in paraffin wax after decalcification. Paraffin embedded samples were stained also with alcian blue and cell centre red staining and with Masson-Goldener solution (Romeis, 1989). Alcian blue is a polysaccharide stain and characterises at a first step EPS. Alcian blue stains carboxyl- and sulphate residues/groups in acidic solution, but not nucleic acids. The dissociation of the carboxyl groups can be suppressed by lowering the pH or by adding high salt concentrations so that only sulphate groups bind the dye (Scott and Dorling, 1965; Hoffmann et al., 2003; Reitner et al., 2004). Thirteen thin sections dyed with calcein and DAPI were analysed with a confocal laser scanning microscope (Nikon A1) at the laboratories Unitech of the University of Milan.

Thirteen subsamples, previously fixed with glutaraldehyde 4%, were prepared for Scanning Electron Microscope (SEM) analyses. Samples were dehydrated in graded ethanol series; ethanol was removed and samples were air-dried. Samples were mounted on stubs, coated with platinum 13 nm thick and analysed with a field emission scanning electron microscope LEO 1530 with Gemini column operating at 3.8 to 15 kV equipped with INCA Energy EDX system from Oxford Instruments at the Geobiology Department of the Georg-August-University Göttingen. Six additional SEM analyses were performed on gold-coated samples, with a Cambridge Stereoscan 360, operating at 20 kV with working distance of 15 mm at the Earth Sciences Department, University of Milan.

X-ray powder diffraction (XRD) analyses of 6 travertine samples form mineralogical determination were performed with an X-RAY Powder Diffractometer Philips X’Pert MPD with high temperature chamber at the University of Milan.

Stable isotope (oxygen and carbon) analyses of 48 samples (9 Bullicame, 15 Bollore, 24 Gorello; Table S1) were performed using an automated carbonate preparation device (GasBench II) connected to a Delta V Advantage (Thermo Fisher Scientific Inc.) isotopic ratio mass spectrometer at the Earth Sciences Department, University of Milan. Carbonate powders were extracted with a dental microdrill and were reacted with > 99 % orthophosphoric acid at 70°C. The carbon and oxygen isotope compositions are expressed in the conventional delta notation calibrated to the Vienna Pee-Dee Belemnite (V-PDB) scale by the international standards IAEA 603 and NBS-18. Analytical reproducibility for these analyses was better than ± 0.1‰ for both δ^18^O and δ^13^C values.

## 4. Results

### 4.1 Travertine depositional setting and thermal water

#### 4.1.1 Bullicame

Bullicame is a Holocene travertine mound with a thickness up to 5-10 m, a diameter of at least 220 m and gentle flank dips, with a central orifice from which thermal water is radially flowing and precipitating carbonates (Figure 2 and Figure S1). The vent circular pool (8 m wide) is surrounded by a fence (Figure 2A-B) limiting sampling only in a channel (20-40 cm wide) departing from the vent. The channel is accessible at a distance of 12.5 m from the centre of the vent orifice (Figure 2A-C), where measured water temperature (Table 1) is 55°C, cooling down to 50°C at a distance of 48.5-60.5 m from the vent and to 48.2°C at a distance of 83.5 m; pH values gradually increase along the channel from 6.7 to 7.4. Alkalinity decreases from 15.6 meq/l to 14.5 meq/l along the channel. The 1-4 cm deep channel has a laminar flow of water. For the first 25 m (12.5-37.5 m from the vent), the channel centre is draped by a dark orange colour microbial mat changing into a green mat overlain by centimetre-size bundles of white carbonate-encrusted filaments oriented with the flow direction; the channel sides are draped by orange/yellow to green microbial mats with embedded sparse millimetre-size gas bubbles not coated by carbonate (Figure 2C-E). The bundles of filaments are poorly to non-calcified in the proximal areas close to the vent and become distally progressively calcified (Figure 2C-D); the channel floor colour gradually changes acquiring a dark green colour below the carbonate-encrusted white fans of filamentous microbes (Figure 2D-E and Figure S1). The Holocene fossil travertines surrounding the active vent show calcified filamentous bundles comparable to the bacterial streamer fabric as labelled by Farmer (2000) in the Mammoth Hot Spring (Yellowstone National Park, Wyoming, USA). Nearly 50 m from the vent the channel becomes progressively dominated by light green to yellow colour microbial mats with abundant gas bubbles, some of which coated by carbonates (Figure 2F and Figure S1H). The centimetre-size levees at the sides of the channel are light orange to green and draped by microbial mats with calcified gas bubbles, carbonate crusts and paper-thin rafts (Figure 2C-F).

**Figure 2.**
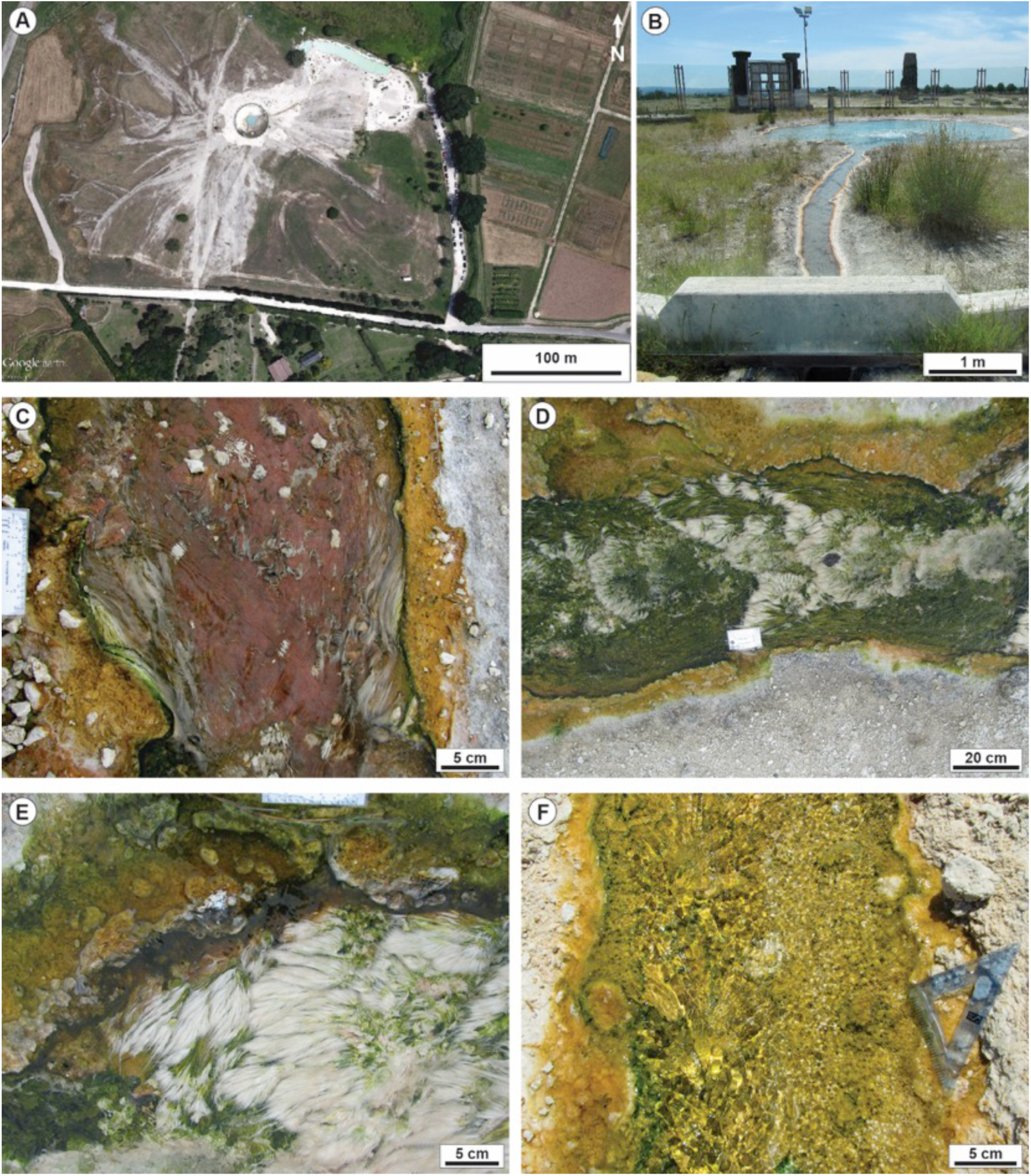
Bullicame (Viterbo) active hydrothermal travertine system. A) Google Earth Pro satellite image showing the Bullicame travertine mound with the fenced active vent in the centre. B) Image of the vent circular pool across the inaccessible glass fence. C) Proximal channel, 14 m from the vent centre, draped by purple microbial mats in the centre, with sparse bundles of white filaments oriented according to the current flow direction and with yellow/orange to green microbial mats on the channel margins. D) Proximal channel further downstream from image in Figure 2C at 22 m from vent centre: the bundles of carbonate-coated filaments are abundant and the channel floor is draped by green microbial mats, while the channel margins are yellow/orange to green in colour with gas bubbles. E) Close-up view of proximal channel at 30 m from the vent centre with bundles of carbonate-coated filaments (bacterial streamers *sensu* Farmer, 2000). F) Distal channel at 60 m from the vent with centre and margins draped by light green to yellow/orange microbial mats with abundant carbonate-coated gas bubbles and rafts.

#### 4.1.2 Bollore

The travertine deposits of the Bollore vent form an asymmetric mound, approximately 10 m thick and 100 m wide (Figure 3 and Figure S2). The main vent at the mound top consists of a circular pool with three orifices (20-50 cm in diameter) from which thermal water outflows into a 15-40 cm wide channel, following the main topographic gradient towards the North (Figure 3A-B). The maximum water temperature is 49.5°C from one orifice, while for the other two orifices temperature is 46.1-46.5°C (Table 1). The values of pH are 6.5-6.6 at the vent and 7.5 nearly 24.5 m from the orifices where temperature drops to 36.4°C. Alkalinity decreases from 31.7 meq/l to 21.6 meq/l 14 m along the channel.

**Figure 3.**
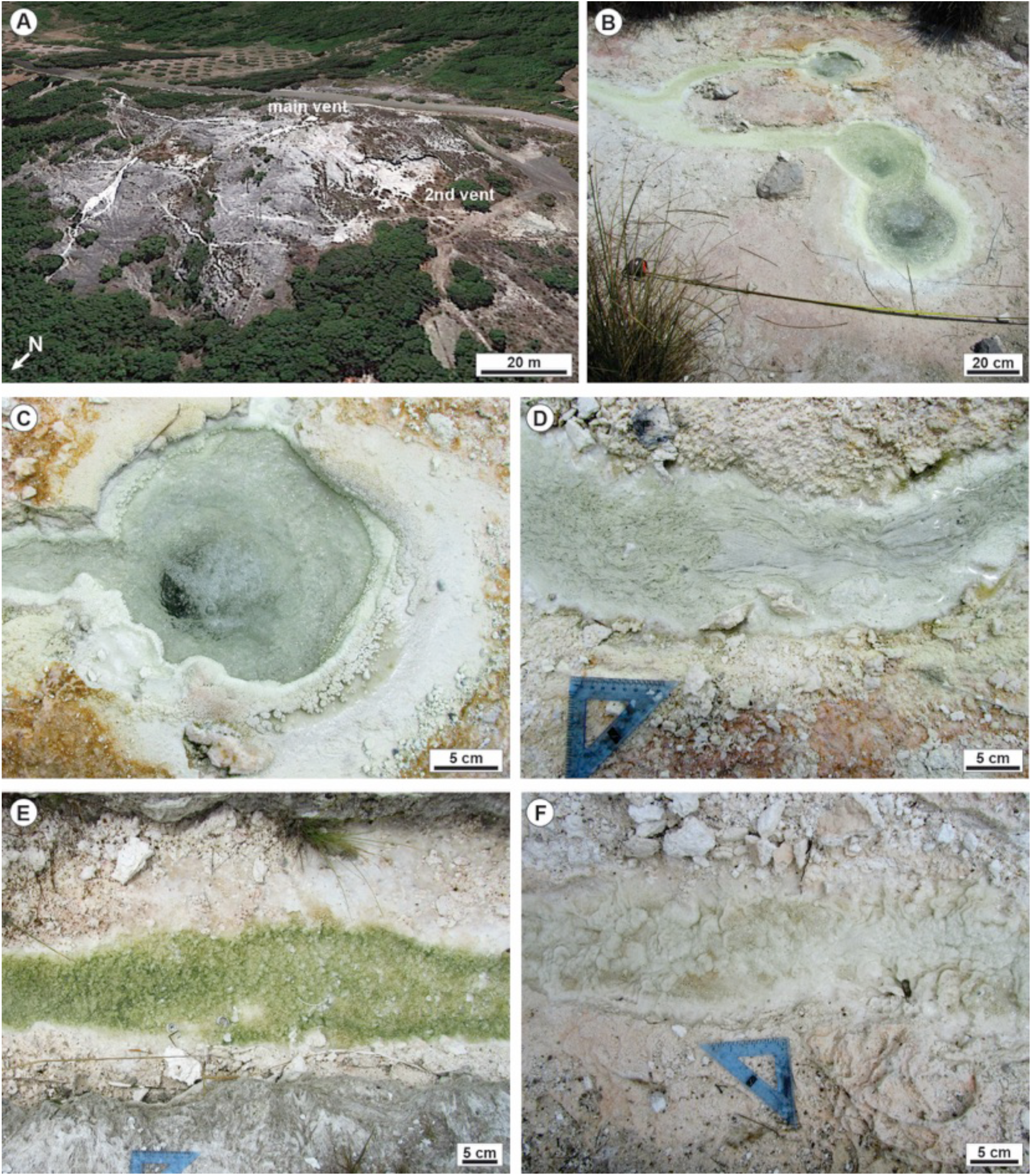
Bollore (Bagni San Filippo) active hydrothermal travertine system. A) Google Earth Pro image showing the Bollore mound and the location of the main vent at the top of the mound and the second, topographically lower, vent. B) Photograph of the three orifices of the main vent and the channel departing on the left side. C) Close-up view of one of the vent orifices with white carbonate precipitates along the rims and pink to orange microbial mat. D) Proximal channel 1 m from the vent orifices with white bundles of carbonate-encrusted filaments oriented following the water flow direction and margins with pink microbial mat. E) Distal channel, 10 m from the vent, with the channel floor draped by green microbial mat, light pink margins and with abundant carbonate-coated gas bubbles. The fossil channel margins (lower part of image) consist of lithified fans of streamers. F) Channel 20 m from the vent where the increase in topographic gradient induces the formation of centimetre-size terraces.

At the rims of the vent orifices and in the first 9 m of the channel, white filamentous bundles encrusted by carbonate form fans 1-2 cm in size (Figure 3C-D) oriented with the water flow direction. These carbonate-encrusted filaments can be observed also as fossil calcified streamer deposits on the channel sides (Figure 3E). Around the vent pool, travertines are coated by orange-pink microbial mat, while decimetre-size stagnant pools with a few millimetres water depth and the sides of the channel are covered by white, submillimetre-thick paper-thin rafts with underlying adherent green microbial mats. From nearly 9 m from the vent, where temperature drops to 44-41°C (Table 1), light green microbial mats drape the channel floor and are associated with millimetre-size carbonate coated gas bubbles and rafts (Figure 3E). At 20 to 25 m from the vent, the topographic gradient increases and the light pink to green channel floor is characterized by centimetre-size terraces and pools with coated gas bubbles and dendrites (Figure 3F). The main vent and channel at Bollore visited during the humid winter season (January) had a different appearance thriving with green to brown/pink coloured microbial mats and abundant carbonate-encrusted filamentous bundles (Figure S2).

A second vent is present at the Bollore mound (Figure 3A and Figure S2) in a lower topographic position, at the base of the mound. Here the temperature measured is 49.3 °C and pH 6.5. The decimetre-wide channel departing from this second vent is draped by carbonate-encrusted filamentous bundles. The channel floor and margins are stained by bright yellow sulphur precipitates and the carbonate and detrital sediments appear black stained by sulphide coatings.

#### 4.1.3 Gorello Waterfall

The Gorello Waterfall travertines form a slope apron fed by a channel running for 1170 m from the thermal spa vent and discharging thermal water at a morphologic break in slope (Figure 4 and Figure S3). The travertine slope apron consists of a waterfall (5 m high) and a terraced slope system that distally merges with a river (Figure 4A-B). Travertines precipitate along the channel, at the waterfall and form the terraced slope (20 m long) with metre-scale sub-horizontal pools separated by rounded rims and sub-vertical walls 0.1-1.5 m high (Figure 4B). Water temperature is nearly 33-33.8°C in the pools; pH values are 7.8-7.9 and alkalinity 9 meq/l (Table 1). The channel levee and the pool rims are coated by olive green millimetre-thick microbial mats (Figure 4C). The pool walls are draped by dark to light green filamentous mats encrusted by carbonate (Figure 4D). Areas of the terraced slope temporarily not flooded by thermal water are sites of vegetation growth, mostly reeds, which are encrusted by carbonate at renewed flows. The pool floor includes terrigenous mud to sand-size detrital sediment, carbonate coated plant fragments and millimetre- to centimetre-size carbonate coated grains (oncoids; Figure 4E-F). The outer surface of these oncoids appears irregular and pitted by green microbial mats (Figure 4F).

**Figure 4.**
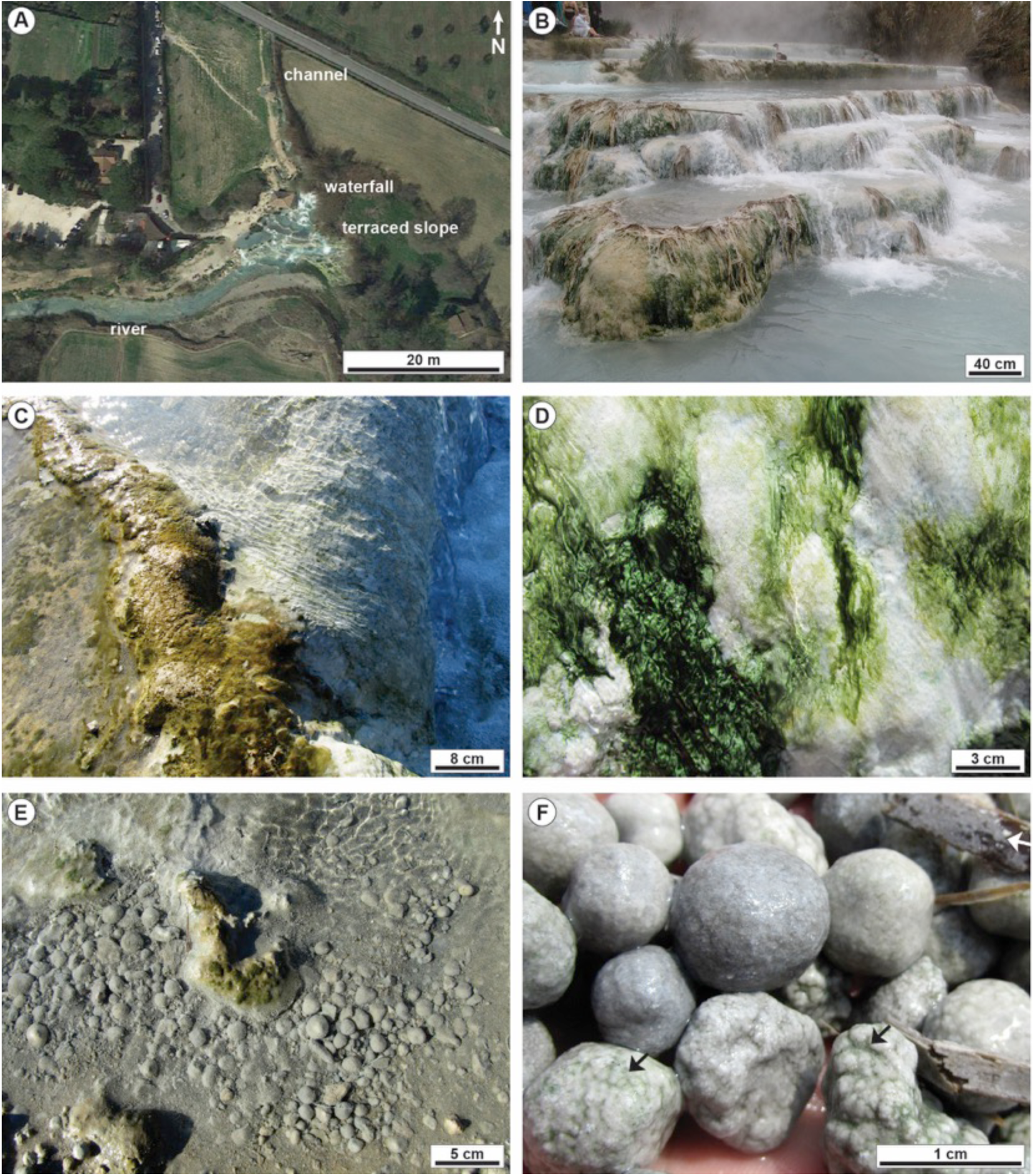
Gorello Waterfall (Saturnia) active hydrothermal travertine system. A) Google Earth Pro image of the Gorello Waterfall showing the channel running from the thermal spa southward, the waterfall developing at the morphologic break in slope where in the past there was a windmill, the terraced slope and the river south of the terraced slope. B) Terraced slope with metre-size pools rimmed by rounded margins with vertical decimetres high walls. C) The pool rims are coated by olive green microbial mat. D) The pool vertical walls are coated by white to dark green filamentous microbial mat. E) The pool floors include detrital mud and siliciclastic grains, microbial mats and centimetre-size carbonate coated grains (oncoids). F) Close-up view of the oncoids with the pitted outer surface with green microbial mats (black arrows). On the upper right corner plant fragments are coated by carbonate (white arrow).

### 4.2 Travertine facies and microbial mats

#### 4.2.1 Bullicame

In the proximal channel centre (Figure 2C-D), carbonate precipitates overlie or are embedded within the microbial mats, made of extracellular polymeric substances (EPS) and filamentous microbes, and follow the shape and structure of the organic substrate (Figure 5A-D and Figure S4). The bundles of coated microbial filaments and EPS are 100 μm to several millimetres thick and variable centimetres long. In proximal samples, calcite crystals consist of microsparite to fine sparite made of euhedral prismatic crystals with a spindle shape (10-80 μm long, 5-40 μm wide), which form radial spherulites with four to ten crystals departing radially outward from a centre (Figure 5B-D). Spherulites (40-120 μm in diameter) are embedded in EPS and filamentous microbes (Figure 5B-E) and contain a micrite clot and/or organic matter nucleus, 10-40 μm in diameter (Figure 5C, F). Calcite crystal size seems to increase downstream reaching 40-120 μm in length and 20-40 μm in width; spherulites diameter rises from average 50 to 200 μm and the micrite nuclei become 50-150 μm in diameter around 30.5 m from the vent (Figure 5F). There are also areas of clotted peloidal micrite with 10-20 μm wide peloids embedded in microsparite mosaics (Figure 5F). The dominant filamentous microbes are associated with rare rod-shaped microbes. Ca-phosphate crystals, likely apatite, are up to 200-300 μm in length. Microbial mat organic matter and some filaments appear birefringent in crossed polarizers and are enriched in Si, Al, Ca, Mg, Na and K or Ca and S measured via EDX. Rare gypsum crystals were also identified as well as detrital pyroxenes from volcanic rocks.

**Figure 5.**
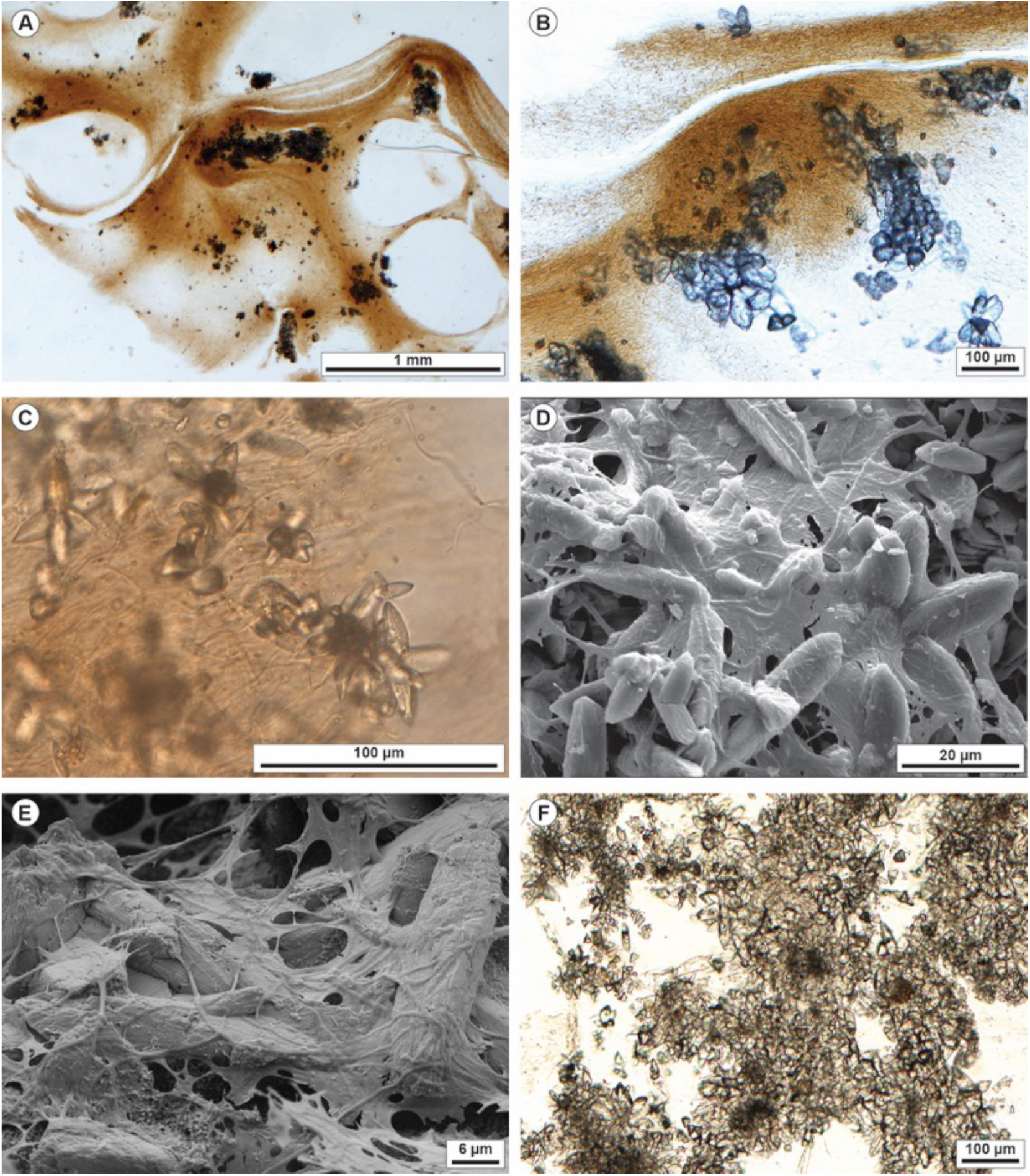
Petrographic and SEM images of the Bullicame travertines in the proximal channel. A) Photomicrograph of sample dyed with tetracycline. The orange colour stained material represents the microbial mat consisting of EPS and filamentous microbes. Carbonates as microsparite/sparite aggregates precipitated within the organic substrate. B) Close-up view of the tetracycline-stained filamentous microbes embedding spherulites of prismatic calcite crystals. C) Calcite crystal spherulites consist of euhedral spindle-shaped calcite crystals around a micrite nucleus embedded within the filamentous microbes and EPS. D) SEM image of calcite crystal spherulites embedded within EPS including filamentous microbes. E) Spindle-shaped prismatic calcite crystals within EPS and filamentous microbes at SEM. F) Photomicrograph of calcite microsparite/sparite mosaic with sparse clots of micrite.

The carbonate crusts (20-100 μm thick) at the channel margin consist of microsparite/sparite spherulites made of prismatic spindle-shaped crystals (10-50 μm long, 5-20 μm wide) that precipitate within EPS, embedding prevalent filamentous, spiral-shaped microbes and rare rod-shaped microbes (Figure 6A-C and Figure S5). SEM analysis shows the presence of hollow spheres (10-20 μm in diameter) made of acicular aragonite crystals, preceding calcite precipitation and Ca-phosphate crystals (Figure S5). Calcite crystals are coated by mucilaginous substances with a grumous appearance, likely EPS, enriched in Si and Al (Figure 6C) or show tubular perforations (Figure S5) from which filamentous microbes are emerging as they were entombed in the crystals.

**Figure 6.**
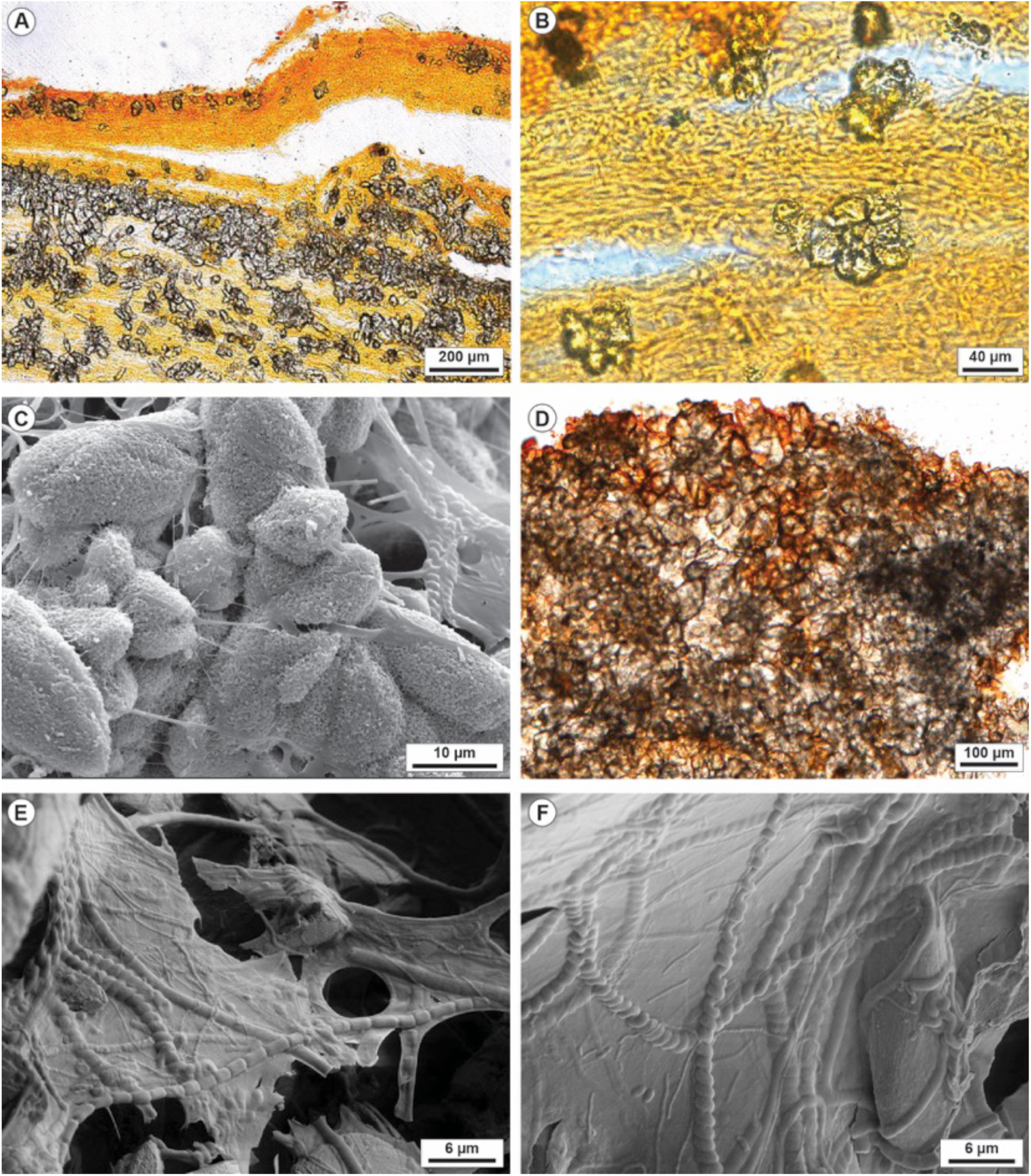
Petrographic and SEM images of the Bullicame travertines in the proximal channel margins (A-C) and distal channel (D-F). A) Photomicrograph of calcein-dyed sample showing the microsparite/sparite spherulites precipitated within the orange-stained microbial mat, spatially distributed following the framework of the organic substrate. B) Close-up view of microsparite spherulites surrounded by EPS and filamentous microbes. C) SEM image showing the prismatic calcite crystals surrounded by grumous EPS with filamentous microbes, also spiral-shaped, that emerge from the crystal as they had been entombed. The grumous mucilaginous organic substance results enriched in Si and Al in EDX analyses. D) Photomicrograph of calcein-stained sample showing the microsparite/sparite mosaic formed by aggregated calcite spherulites and micrite clots. E) SEM image of EPS embedding spiral-shaped and segmented filamentous microbes. F) SEM image of EPS embedding spiral-shaped and various filamentous and coccoid microbes and prismatic calcite crystals.

In the distal channel (Figure 2F), the carbonate precipitates are similar to the proximal part in terms of crystal shape, size and organization in radial spherulites with often micrite nuclei, 50-300 μm in diameter (Figure 6D and Figure S5). Calcite spherulites precipitate within EPS embedding filamentous microbes (Figure 6E-F and Figure S5), which are spiral-shaped or segmented associated with rare coccoid or rod-shaped forms (Figure 6E-F). Calcite crystals show tubular perforations with filamentous microbes exiting the hollow tube as they had been entombed in the crystal. Calcite crystals have locally a coating of grumous EPS enriched in Si and Al or P. Calcium phosphate crystals (Figure S5) are draped by filamentous microbes and EPS.

#### 4.2.2 Bollore

At Bollore main vent, bundles of filamentous microbes are 50-500 μm wide and act as organic substrates for carbonate precipitation (Figure 7A and Figure S6). Carbonate precipitates consist of prismatic euhedral calcite crystals (20-100 μm long, mostly 50-60 μm, 5-20 μm wide) organized in spherulitic structures (diameters 50-300 μm) with micrite nuclei, 20-150 μm in diameter, or coating micritic filament with diameter 10-20 μm (Figure 7B). Clotted peloidal micrite occurs in patches up to 200-300 μm in diameter embedded in sparite mosaics. Calcein and tetracycline dyes stain the micritic nucleus indicating the presence of organic matter associated with free Ca^2+^ in the micrite. Calcite crystals are embedded in EPS with abundant filamentous microbes (diameters 0.2-1 μm), some spiral-shaped, and sparse, 1-2 μm size, rod-shaped microbes (Figure 7C-D). Some calcite crystals are coated by a grumous organic film, which appears to be transitional to and/or overlain by EPS and shows square-shaped moulds (Figure 7C). This organic material coating the calcite crystals, EPS and microbes are enriched in Al, Si, S, Ca and K. Samples proximal to the vent contain acicular aragonite spherulites that reach diameters of 7-20 μm (Figure 7D and Figure S6) and seem to have precipitates both before and after calcite. Ca phosphate and euhedral (swallow tail) crystals of gypsum, forming also rosettes up to 200 μm in diameter, postdate calcite and aragonite precipitation (Figure S6).

**Figure 7.**
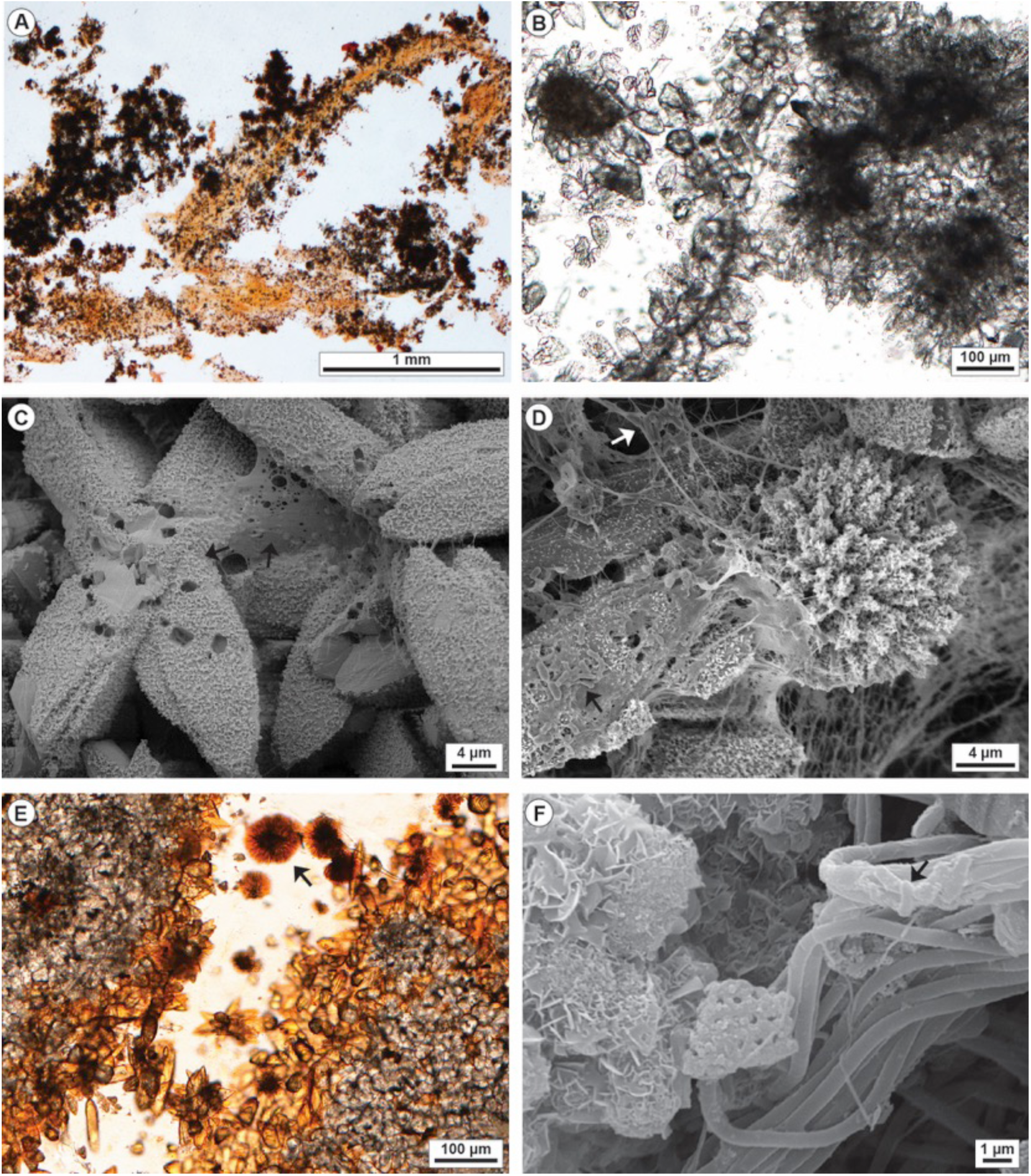
Petrographic and SEM images of the Bollore (Bagni San Filippo) travertines at the main (A-D) and second vent (E-F). A) Photomicrograph of sample dyed with calcein showing carbonate precipitates draping and embedded within orange colour organic substrate of the filamentous microbe bundles. B) Photomicrograph showing the euhedral prismatic microsparite/fine sparite calcite crystals forming spherulites and aggregates of spherulites with a central micrite nucleus or around elongated micrite filaments. C) SEM image showing the radial arrangement of the prismatic calcite crystals with squared moulds, coated by grumous EPS (black arrows). D) SEM image of aragonitic spherulites and calcite crystals embedded in EPS with rod-shaped microbes (black and white arrows). E) A mosaic of microsparite/fine sparite crystals forming spherulites or irregular aggregates around micrite clots and aragonite spherulites (black arrow) stained orange by calcein dye reacting with organic matter. F) SEM image of the bundles of filamentous microbes associated with rod-shaped microbes (black arrow). On the left side a reticulate aggregate of aluminium-silicate, probably an authigenic clay mineral, forming on the filamentous microbes.

At the second Bollore vent, carbonate precipitates show similar features to the main vent: prismatic calcite crystals 5-40 μm wide, 30-120 μm long forming spherulites dispersed in the EPS or clustered forming mosaics, and abundant aragonite spherulites, 20-200 μm in diameter (Figure 7E). Filamentous microbes (0.1 to 1 μm cross section size), rarely spiral-shaped, are dominant but associated with sparse rod-shaped (Figure 7F) and coccoid microbes around 1 μm in size. In the channel 2 m from the vent, spiral-shaped filaments become common. Aggregates of minerals with Si, Al, Mg, Ca composition form a reticulate fabric and might represent authigenic clay mineral (Figure 7F); these authigenic aluminium-silicate form aggregates on the filamentous microbe bundles but also coat some individual microbial filaments and calcite crystals (Figure 7F and Figure S6).

In the proximal channel departing from the main vent, there are centimetre-size fan-shaped bundles of filamentous microbes (bacterial streamers *sensu* Farmer, 2000) encrusted by euhedral prismatic calcite crystals (15-80 μm long, 5-30 μm wide but also 5-2 μm in size), sometimes with gothic-arch shape, organized in spherulites with a micrite clot at the nucleus or aligned along micritic filamentous structures (Figure 8A-B and Figure S7A-B). Carbonate precipitates are similar to those at the vent pools with more abundant acicular aragonite spherulites, in general 30-100 μm in diameter but also millimetre size, lining the precipitated calcite (Figure 8A-B). Gypsum crystals (50-80 μm long, with swallow tail habit) form rosettes 100-200 μm in diameter (Figure 8C), and phosphates postdate calcite precipitation. All the mineral precipitates are surrounded by EPS and filamentous and rod-shaped microbes (Figure 8C and Figure S7C-D).

**Figure 8.**
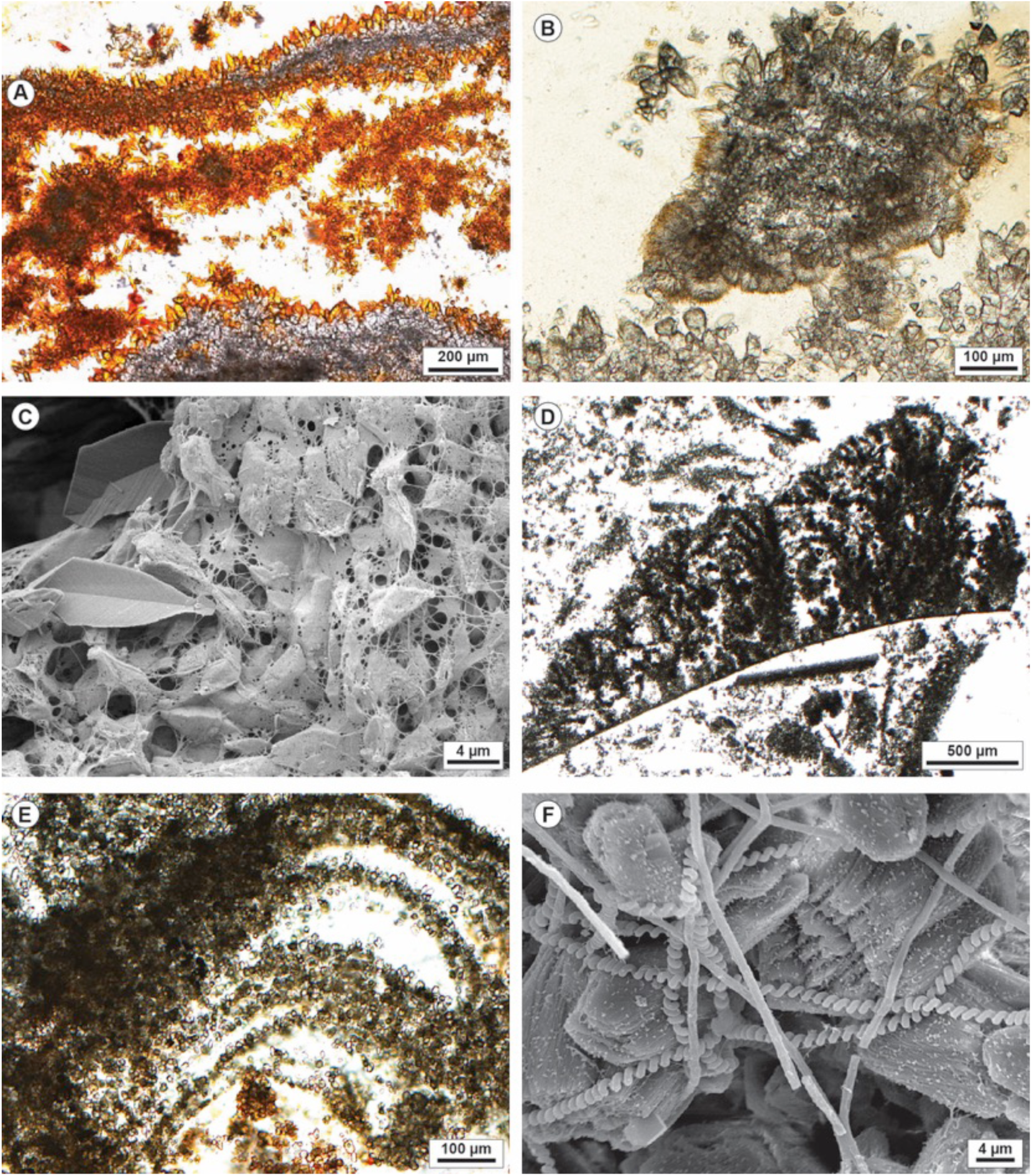
Petrographic and SEM images of the Bollore (Bagni San Filippo) travertines at the proximal (A-D) and distal channel (E-F). A) Carbonate encrusted bundles of filamentous microbes are characterized by microsparite to fine sparite euhedral crystals coating micritic filaments and peloids. The orange colour is related to the calcein dye. B) Calcite microsparite and micrite aggregates coated by acicular aragonite crystal fans. C) SEM image showing EPS embedding micrite calcite crystals and a swallow-tail gypsum crystals on the left side. D) Distal channel carbonate precipitates are characterized by abundant micrite and clotted peloidal micrite forming dendritic structures, rafts and coated gas bubbles. E) Laminae made of microsparite and micrite supported by microbial mat and filamentous microbes. F) SEM image of calcite prismatic crystals overlain by filamentous microbes including spiral-shaped *Spirulina* cyanobacteria.

At a distance of 14 m from the vent (Figure 3; Figure 8D-E and Figure S7E-F), there are no aragonite, gypsum or phosphate crystals and precipitated carbonates are dominated by micrite and microsparite crystals (4-20 μm in size, mostly 4-10 μm) with spherulites diameters up to 10-50 μm. Filamentous microbes, largely spiral-shaped, are abundant (Figure 8F and Figure S7G-H).

#### 4.2.3 Gorello Waterfall

Samples from the pool rims and walls of the Gorello Waterfall (Figure 4) include a 0.5-1.5 mm thick superficial microbial mat with only sparse carbonate precipitates, followed by a centimetre-thick laminated carbonate deposit with alternation of carbonate precipitated laminae and organic matter films (Figure 9 and Figure S8, S9, S10). Microbial mats consist of a network of undulated filamentous microorganisms oriented mostly up-right, perpendicular to the lamination, or prostrated horizontally. Carbonate precipitates form subparallel undulated laminae or a framework of horizontal, oblique and vertical crusts mimicking the spatial distribution of biofilm EPS and filamentous microbes (Figure 9A-D and Figure S8, S9, S10). Carbonates occur only embedded in organic matter and follows the alveolar structure of the EPS (Figure 9D). Pore spaces of the microbial mat framework lack carbonate precipitates, which appear to occur within the EPS following the orientation of microbial filaments (Figure 9C-E). Carbonate precipitates consist of clotted peloidal micrite and euhedral microsparite to fine sparite crystals (5-100 μm in size). Calcite crystals are prismatic to sub-equant with rhombohedral or trigonal cross-section or dodecahedral shape, often organised in radial spherulitic structures with a diameter of 10-200 μm (Figure 9E-F and Figure 10A-B) and nuclei of peloids and organic matter, 5-20 μm in diameter. Spherulites can occur embedded in clotted peloidal micrite or surrounded by erect filamentous microbes or suspended in EPS. Filamentous microbes can depart radially from the calcite rosettes (Figure 10A) and emerge from hollows within the crystals as they were entombed during crystal growth (Figure 10B and Figure S8, S9, S10). Precipitated carbonate occurs also in the form of micrite with nanometre scale particles forming aggregates. Filamentous microbes with a diameter of 1 μm are dominant, associated with thicker segmented filamentous forms (4-5 μm in diameter) with an outer sheath (Figure S8, S9, S10), spiral-shaped, segmented and chain-like filamentous microbes (Figure 10C), rod-shaped microbes and pennate diatoms. Micritic carbonate precipitates encrust the microbial filaments forming a carbonate tube coating them (Figure 10D). In addition to calcite, other authigenic minerals observed are Ca-phosphate crystals (platy crystals and spongy texture Figure S8, S9, S10), gypsum, framboidal pyrite and Si-Al minerals. Authigenic silicates (Figure 10E) occur as aggregates among the calcite crystals and as coatings or infillings of filamentous microbes that appear birefringent in crossed polarizers and indicate, together with EPS, a chemical composition enriched in Si, Al, K, Ca, Mg, Fe or S and Ca.

**Figure 9.**
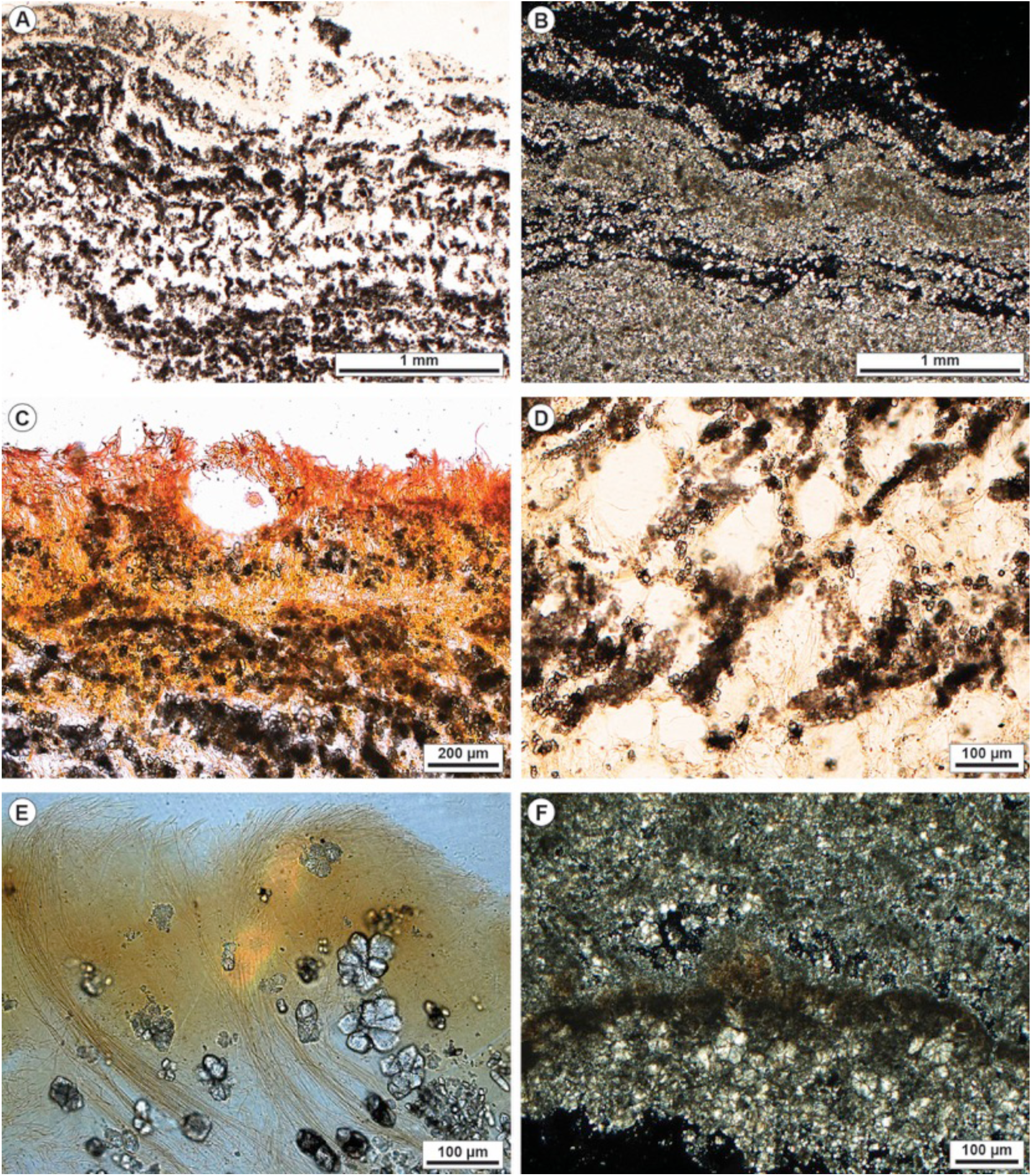
Petrographic analysis of the Gorello Waterfall (Saturnia) travertines from the rims and walls of the terraced slope system. A) Travertine laminated boundstone made of laminae of clotted peloidal micrite and microsparite spherulites alternating with organic matter consisting of filamentous microbes and EPS. B) Crossed-polarizers image of undulated laminated boundstone made of clotted micrite and microsparite spherulites. Carbonate precipitates follow the geometry of the organic substrate of the microbial mat. C) Calcein-dyed sample showing the outer surface of the laminated boundstone with up-right filamentous microbes and clots of micrite/microsparite distributed within the microbial EPS and in between erect filamentous microbes. D) Alveolar network of microbial filaments and EPS within which carbonate spherulites and micrite clots precipitate mimicking the EPS alveolar framework. E) Outer portion of the laminated boundstone with microsparite spherulites embedded in between the undulated up-right filamentous microbes. F) Crossed-polarizers image of a framework of microsparite spherulites and patches and laminae of clotted peloidal micrite.

**Figure 10.**
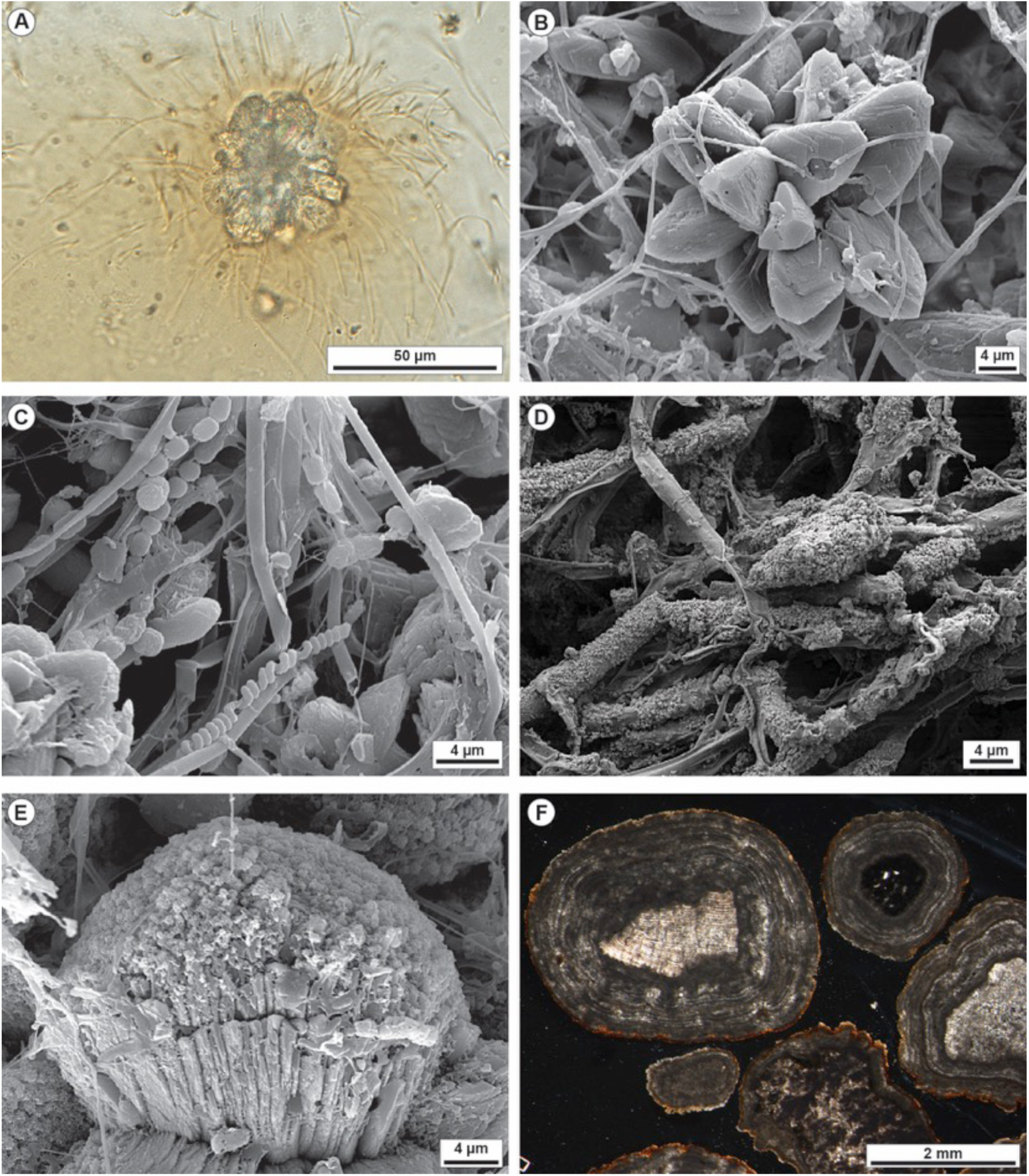
Petrographic and SEM images of the Gorello Waterfall (Saturnia) travertines from the rims, walls and pools of the terraced slope system. A) Photomicrograph of a microsparite spherulite with filamentous microbes departing from the radial crystalline structure. B) SEM image of prismatic calcite crystals forming a spherulite. Filamentous microbes emerge from tubular moulds within the crystals as they were entombed during crystal growth. C) SEM image showing a diverse community of filamentous microbes including spiral-shaped, segmented and chain like probably cyanobacteria. D) SEM image of filamentous microbes encrusted by precipitated micrite. E) Bladed calcite crystals covered by Si-Al minerals, probably authigenic clay mineral. F) Crossed-polarizers image of pool floor oncoids with nuclei made of detrital intraclasts or substrate rock extraclasts and cortex consisting of an alternation of micrite and microsparite laminae. The tetracycline dye shows that the outer surface of the oncoids consists of Ca^2+^ rich organic matter.

Coated grains (oncoids) in pools (Figure 4E-F, Figure 10F and Figure S11) have nuclei varying from travertine intraclasts, clasts of substrate rocks forming the detrital sediment accumulating in the pools (fragments of fluvial tufa with peloidal packstone, coated plants and calcified cyanobacteria, peloidal packstone, sandstone, quartz grains, planktonic and benthic foraminifers from the Pliocene marine shales), and plant stem fragments. The precipitated cortex varies from predominant micritic laminae to an alternation of micrite (4-50 μm thick) and microsparite/sparite (10-200 μm thick) laminae with crystals from equant to bladed, oriented perpendicular to the underlying micritic lamina forming palisades or crystalline fans adjacent to each other. The micritic laminae can be discontinuous and develop millimetre-size columnar structures. The outer rim of the oncoids is rich in organic matter as well as some internal laminae (Figure S11); filamentous microbes might extend outward from the mat coating the outer surface of the oncoids. In between the calcite crystals there are EPS and filamentous microbes oriented parallel to the crystals growing perpendicular to the laminae.

### 4.3 Travertine stable carbon and oxygen isotopes

The carbon and oxygen stable isotope data of the three investigated travertine sites are plotted in Figure 11 and summarised in Table S1. The three travertine localities plot in distinct fields of the δ^13^C-δ^18^O diagram with some overlap between Bullicame and Bollore. All the three data sets show a positive correlation between δ^13^C and δ^18^O values. Bullicame travertines are characterized by the highest δ^13^C values for the rafts and coated bubbles and coated reeds (average δ^13^C 7.3 and 6.7 ‰, respectively; average δ^18^O −9.4 ‰ for both); the streamer fabrics show ^13^C depleted δ^13^C (average 5.4 ‰) and ^18^O depleted δ^18^O (average −11.8 ‰). The Bollore travertines show lower carbon and oxygen isotope values than Bullicame but the stable isotope signatures partly overlap. Rafts, coated bubbles and dendrites are characterized by higher values of δ^13^C and δ^18^O (average δ^13^C 5.3 and 5.4 ‰, average δ^18^O −12.0 ‰ and −12.4 ‰, respectively) with respect to the streamer fabrics (average δ^13^C 4.5; δ^18^O −12.6 ‰) as observed in Bullicame. The Gorello Waterfall travertines have an isotopic signature characterized by fairly uniform δ^18^O, around −8.5 ‰, and δ^13^C ranging between 2.2 and 3.5 ‰.

**Figure 11.**
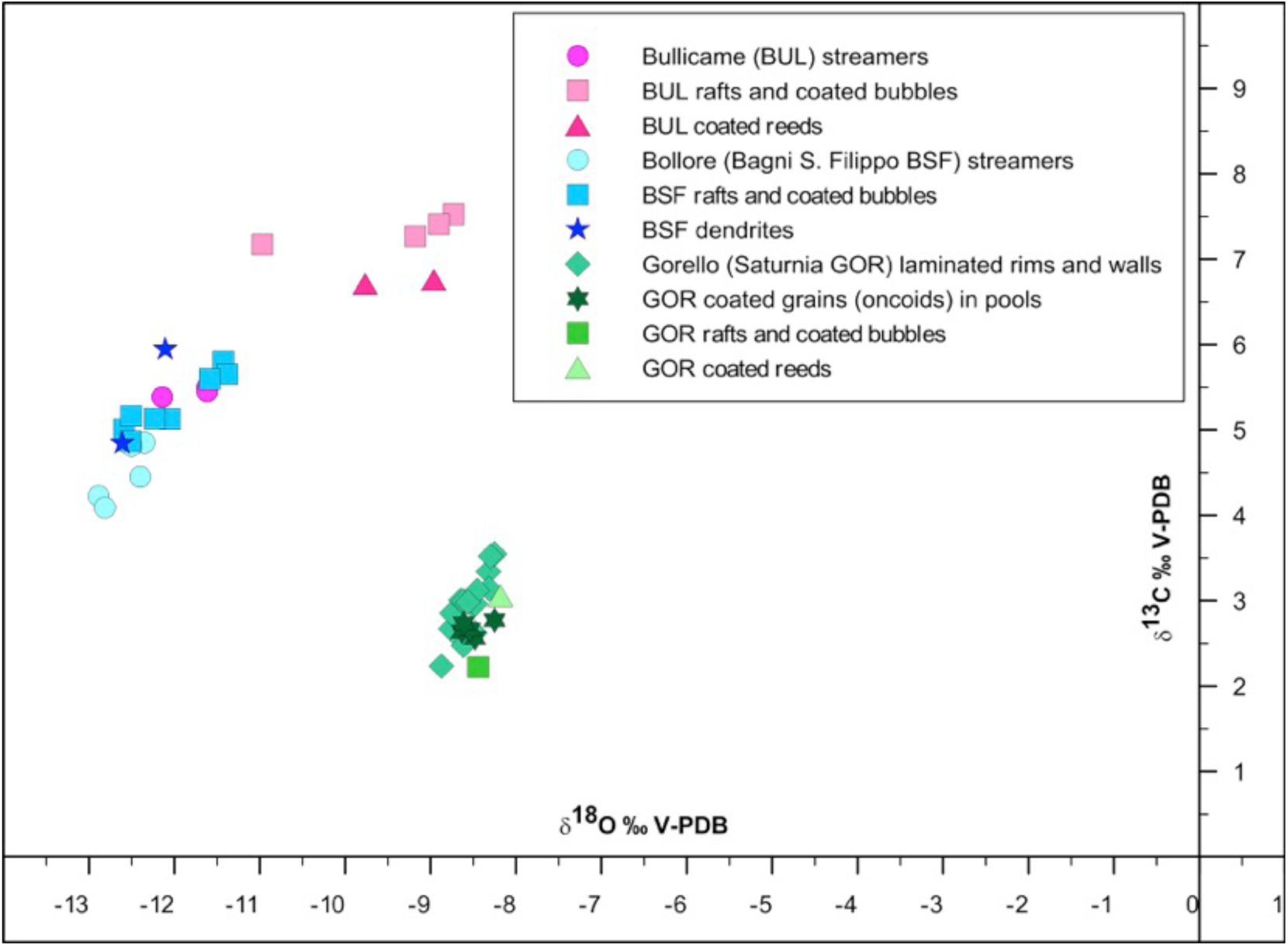
Plot of the carbon and oxygen stable isotope measurements of the three analysed case studies.

## 5 Interpretation and Discussion

The three investigated travertine sites are characterized by different thermal water temperature, pH and alkalinity and different microbial mat composition. The mesoscale to macroscale travertine facies vary according to local water temperature, flow velocity, turbulence, topographic gradients (cf. Capezzuoli et al., 2014; Gandin and Capezzuoli, 2014; Della Porta, 2015 Della Porta et al., 2017ab) and organic substrate for carbonate precipitation, whereas, at the microscale, carbonate precipitates appear similar, with only some differences with respect to carbonate crystal mineralogy, size and shape.

### 5.1 Characteristics of precipitated minerals

At the microscale, despite the differences in size and morphology, calcite and aragonite crystals are similarly organized in spherulites precipitating within and above the microbial mat.

Previous studies suggested that the prismatic shapes of travertine calcite crystals are related to crystalline lattice and growth morphologies being affected by impurities, concentrations of Mg, phosphate or sulphate, high fluid supersaturation with respect to carbonates, and presence of microbial biofilms (Folk et al., 1985; Folk, 1993, 1994; Tracy et al., 1998; Bosak and Newman, 2005; Di Benedetto et al., 2011; Jones and Peng, 2014a; Jones, 2017a). Numerous experimental studies on the precipitation of carbonate crystals confirm that crystal morphology is influenced by the presence of impurities, hydrogels, polymers and various organic molecules (Gower and Tirrell, 1998; Falini et al., 2000; Meldrum and Hyde, 2001; Kato et al., 2002; Cölfen, 2003; Oaki and Imai, 2003; Chekroun et al., 2004; Tong et al., 2004; Bosak and Newman, 2005; Sand et al., 2011; Keller and Plank, 2013; Tobler et al., 2014; Kosanović et al., 2017). Konopacka-Lyskawa et al. (2017) showed that calcite crystal morphology changes from scalenohedral to rhombo-scalenohedral elongated crystals and crystal size decreases at increasing organic compound concentrations. Also the amount of dissolved silica seems to affect calcium carbonate crystal morphology and size (Lakshtanov and Stipp, 2010). At hydrothermal springs, carbonate minerals precipitate in conditions far from equilibrium and develop a wide spectrum of crystal morphologies ranging from monocrystals, mesocrystals, skeletal crystals, dendrites, and spherulites at increasing disequilibrium conditions and non classical crystal growth pattern (Jones and Renaut, 1995; Jones, 2017a). Non classical crystal growth produces mesocrystals, which are mesoscopically structured crystals made of nanocrystals or ACC (amorphous calcium carbonate) arranged into an isooriented crystal via oriented attachment that shows birefringent properties of a single crystal (Cölfen and Antonietti, 2005; Meldrum and Cölfen, 2008; Rodriguez-Navarro et al., 2015; Jones, 2017a). The development of ACC in travertines may be widespread and may play a critical, but transitory role, in the development of crystalline CaCO_3_ in high temperature (Jones and Peng, 2012; Peng and Jones, 2013; Jones and Peng, 2014ab, 2016), but also in ambient temperature environments (Pedley et al., 2009; Pedley, 2014). Nevertheless, in the studied travertines, SEM observations did not provide clear evidence of mesocrystal formation through the aggregation of nanometre-scale particles in the higher temperature Bullicame and Bollore calcites. At Bollore, in some cases acicular aragonite crystals departing from the spherulite micritic nucleus appear to result from aggregation of nanometre micrite. In the lower temperature Gorello Waterfall travertines, nanometre-scale micrite precipitates are common and mesocrystals, derived from the aggregation of nanoparticles, comparable to published examples were rarely identified (cf. Pedley et al., 2009; Pedley, 2014; Jones and Peng, 2014a).

In travertines in Central Italy, aragonite generally precipitates at sites with water temperature >40°C or Mg/Ca ratio > 1 (Folk, 1993, 1994). Jones (2017b) proposed alternative mechanisms driving aragonite vs. calcite precipitation in travertines suggesting that aragonite may precipitate either due to rapid CO_2_ degassing, to high Mg/Ca ratio or due to the presence of different micro-domains within microbial biofilms. Precipitation of different carbonate minerals could also be controlled by the acidic EPS organic matrix. The distances of the COO- groups initiate the nucleation via 001 plane of the carbonate crystal and determine the type of mineral (e.g., 4.99Å = calcite, 4.96Å = aragonite, 4.13Å = vaterite; Addadi and Weiner, 1989). However, often the COO- groups are not at the right distance and the initial carbonate phase is amorphous (ACC) and, in later stages, the better ordered crystal nanostructure can form faced crystals (cf. Gong et al., 2012; Ma and Feng, 2014; Rodriguez-Navarro et al., 2015).

Calcite and aragonite spherulites have been observed in various travertine and lacustrine case studies and attributed to microbial mediation or EPS influence and presence of organic acids (Guo and Riding, 1992; Folk, 1993; Dupraz et al., 2009; Arp et al., 2012; Mercedes Martin et al., 2016; Kirkham and Tucker, 2018), or abiotic precipitation within silica gels (Wright and Barnett, 2015; Tosca and Wright, 2018). Spherulites are polycrystals commonly linked to high supersaturation levels and far from equilibrium precipitation (Jones and Renaut, 1995; Jones, 2017a). Spherulites seem to form due to increased supersaturation and slow diffusion in a viscous gel media (Tracy et al., 1998; Oaki and Imai, 2003; Sanchez-Navas et al., 2009). The addition of impurities, such as additives, polymers, organic molecules, Mg and sulphate ions, may be responsible for spherulitic growth (Fernández-Díaz et al., 1996; Tracy et al., 1998; Davis et al., 2000; Sanchez-Navas et al. 2009; Shtukenberg et al., 2011). However, Andreassen et al. (2012) suggested that spherulites may form also without additives. Chekroun et al. (2004), instead, proposed that spherulitic and dipyramid crystals in natural mucilaginous biofilms are linked to biologically induced mineralization. Braissant et al. (2003) demonstrated, through laboratory experiments, that calcium carbonate polymorphs (calcite and vaterite) form spherulites at increasing concentrations of EPS and amino acid acidity, which enhance sphere formation and the morphology of calcite crystals evolves from rhombohedral to needle-shape due to stretching along the c axis as the amino acid changes from glutamine to aspartic acid and as the medium is progressively enriched in EPS.

There is no definite explanation for the formation of Ca-phosphate in the three investigated sites. Ca-phosphate might be a fixation artefact because PBS (Phosphate Buffer Solution) was used. However, Ca-phosphate crystals are observed draped by EPS and microbes and might represent authigenic precipitates (Figure S5H and Figure S10C). Gypsum crystals might be related to the abundant presence of H_2_S in thermal water that is oxidized to elemental sulphur and sulphate either due to mixing with atmospheric oxygen or due to microbial activity (e.g., sulphide oxidizing bacteria, green sulphur anoxygenic phototroph bacteria). Authigenic Ca-Mg aluminium-silicates occur as aggregates on the calcite crystals, EPS and as coatings/fillings of the filamentous microbes. The impregnation of EPS with Si, Al, Ca, Mg, K and Fe, and the formation of amorphous silica or authigenic aluminium-silicate minerals have been observed in various hydrothermal and lacustrine case studies and attributed to biologically induced and influenced mineralization (Allen et al., 2000; Konhauser et al., 2001; Arp et al., 2003; Lalonde et al., 2005; Pentecost, 2005; Jones and Peng, 2012; Peng and Jones, 2013; Jones and Peng, 2014ab, 2015, 2016; Kremer et al., 2019).

### 5.2 Microbial mats associated with travertines

The identification of microbial communities based on morphologies observed at SEM is tentative because microbe morphologic characteristics do not reflect phenotypic characteristics and it is difficult to ascribe metabolism based on morphology (cf. Giovannoni et al., 1987; Reysenbach and Cady, 2001). Nevertheless, the three sites show that there is a temperature control on the composition of the microbial mats and within the same system from proximal to distal cooled-down settings (Figure 12). Various studies have suggested a temperature control on the composition of microbial communities in hydrothermal terrestrial settings (Cady and Farmer, 1996; Farmer, 2000; Dunckel et al., 2009; Di Benedetto et al., 2011; Djokic et al., 2017; Des Marais and Walter, 2019; Sanchez-Garcia et al., 2019; Gong et al., 2020).

**Figure 12.**
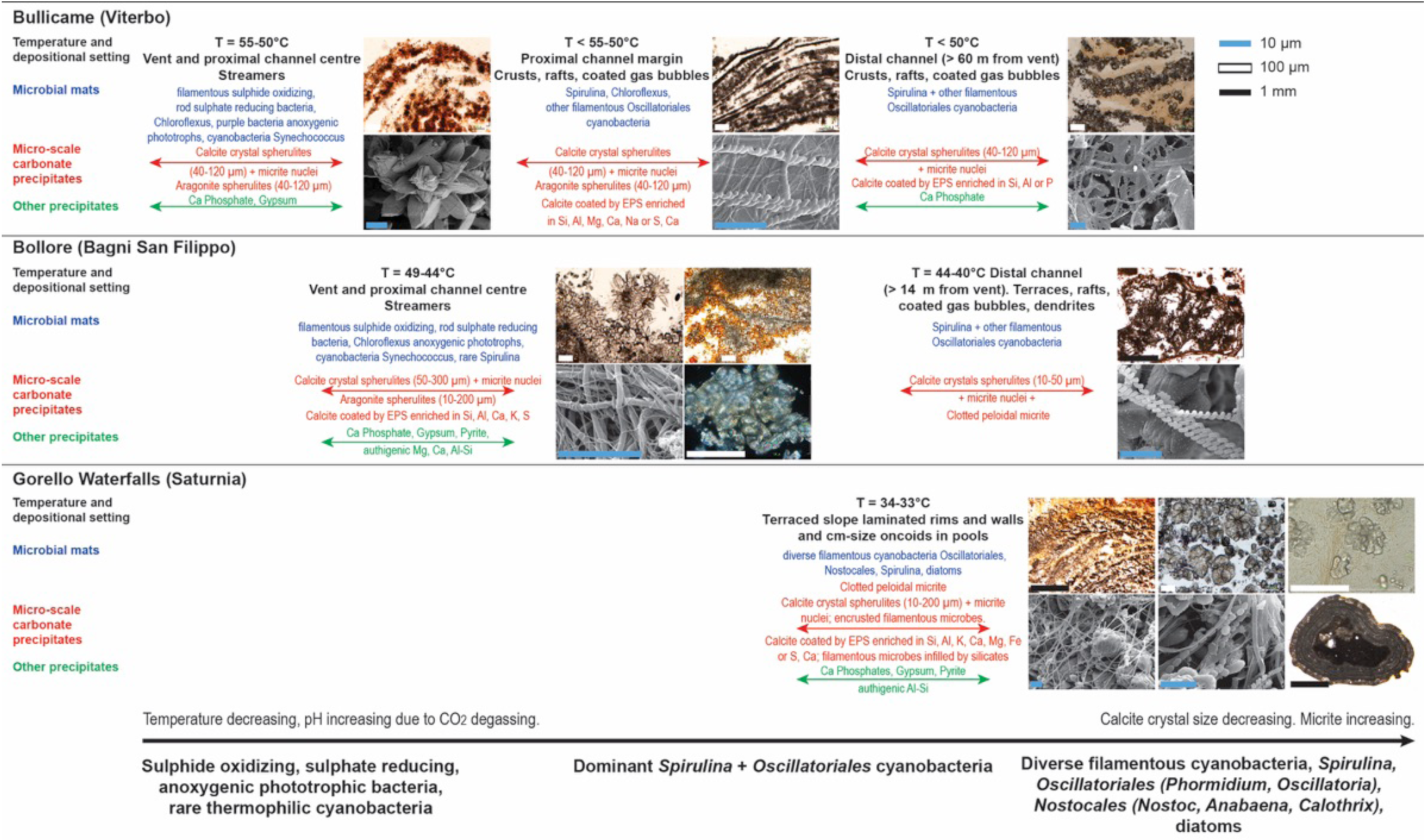
Diagram summarizing the main features of the carbonate and other mineral precipitates and associated microbial communities in the three study sites at decreasing temperature.

#### 5.2.1 Bullicame travertine facies and microbial mats

In the three Bullicame sampling sites (i.e., proximal channel centre and margins, distal channel), at the microscale the predominant carbonate precipitates are similarly consisting of microsparite to fine sparite prismatic calcite crystals with a spindle shape forming spherulites, despite the different mesoscale carbonate fabrics varying from calcified streamers in the proximal channel to rafts and coated gas bubbles in the distal channel. Carbonate precipitation occurs exclusively above or within the organic substrates provided by the microbial mats that control the spatial distribution of the precipitated travertines. Calcite crystals show numerous perforation as tubular pores from which filamentous microbes emerge. These pores seem moulds formed by entombment of microbes during fast crystal growth rather than borings of endolithic microorganisms. A difference in the three sampling sites is that aragonite spherulites occur only in the proximal location of the channel centre and margins where temperature is above 50°C. The proximal channel centre (50-55°C) is dominated by filamentous bacterial streamers and purple microbial mats associated with rod-shaped bacteria. These streamers might be identified as sulphide oxidizing bacteria and/or green non sulphur anoxygenic photoautotroph bacteria belonging to the phylum Chloroflexi as identified in various hydrothermal travertine settings (Folk, 1993; Allen et al., 2000; Farmer, 2000; Fouke et al., 2000; Pentecost, 2003; Pentecost and Coletta, 2007; Fouke, 2011; Valeriani et al., 2018). At Mammoth Hot Springs, the streamer forming filamentous microbes were identified as thermophilic sulphide oxidizing bacteria of the Aquificales group (Farmer, 2000; Fouke et al., 2000; Reysenbach et al., 2000; Reysenbach and Cady, 2001; Fouke et al., 2003; Veysey et al., 2008; Fouke, 2011). They occur at the vent, apron and channel facies in water temperature of 73-60°C (Fouke et al., 2000; Fouke, 2011). The Aquificales bacterium *Sulfurihydrogenibium yellowstonensis* is a chemolithoautotrophic microbe fixing CO_2_ via sulphur oxidation (Fouke et al., 2003; Fouke, 2011). The intermediate temperature settings (62-39°C) of the pond facies are dominated by the filamentous green non sulphur anoxygenic phototroph *Chloroflexus*, associated with cyanobacteria (*Oscillatoria*, *Spirulina*, *Synechococcus*), green sulphur bacteria (*Chlorobium*) and β-proteobacteria (Farmer, 2000; Fouke et al., 2000, 2003). *Chloroflexus aurantiacus* occurs at temperatures of 65-55°C with the purple sulphur anoxygenic phototrophic bacterium *Thermochromatium tepidum* (Giovannoni et al., 1987; Madigan, 2003; Fouke, 2011). In the proximal slope facies (63-35°C), the presence of sulphate reducing bacteria is confirmed by clones of *Desulfovibrio*, which is associated with green non sulphur bacteria (*Heliothrix*), cyanobacteria (*Pseudoanabaena, Spirulina, Synechococcus*), *Thermus-Deinococcus* and *Chlorobium* green sulphur bacteria (Fouke et al., 2003; Fouke, 2011). Giovannoni et al. (1987) determined that there is 20% decrease in night time sulphate concentrations in the pond spring-water indicative of enhanced rates of sulphate reduction when oxygenic photosynthesis is reduced. The distal slope environments (44-28°C) are characterized by high diversity of microorganisms with dominant cyanobacteria (*Spirulina*, *Calothrix*, *Synechococcus*) but also diatoms, algae and plants (Farmer, 2000; Fouke et al., 2000, 2003; Fouke, 2011). The Yellowstone silica depositing thermal springs evaluated by Cady and Farmer (1996) are characterized by green non sulphur *Chloroflexus* and *Synechococcu*s cyanobacteria at temperatures of 60-73°C, whereas cyanobacteria *Phormidium* and *Spirulina* occur at 59-35°C and the association of *Phormidium* and *Calothrix* cyanobacteria thrives at temperatures less than 35°C.

Folk (1993) indicated the presence of anoxygenic photosynthetic microbes (green sulphur *Chlorobium*, purple sulphur bacterium *Chromatium*, green non sulphur *Chloroflexus*), *Thiobacillus* (sulphur oxidising proteobacteria) and *Oscillatoria* cyanobacteria in the Bullicame and Bagnaccio (nearly 6 km North of Bullicame) hydrothermal vents at temperature of 56-64°C. At Le Zitelle vent (nearly 1 km NW of Bullicame), at temperatures of 61-50°C, the carbonate encrusted filamentous mats were attributed to *Chloroflexus* associated with the sulphur reducing bacterium *Desulfovibrio thermophilus*, whereas at temperatures < 50°C the microbial mat was dominated by the cyanobacteria *Spirulina* and *Synechococcus elongatus* (Folk, 1994; Allen et al., 2000; Pentecost and Coletta, 2007).

Valeriani et al. (2018) performed metagenomics analysis of the Bullicame microbial mats sampled at 54°C. The channel water was dominated by the sulphur oxidizing bacteria *Thiofaba*, which was very rare in the channel mat; sulphate reducing bacteria were common in both water and microbial mat. The anoxygenic phototrophs *Chloroflexus* and *Roseiflexus* were present both in water and mat but they did not dominate; the Archaea community was very scarce in water and mat samples (Valeriani et al., 2018). In the channel lithified microbial mat, the phyla identified were Proteobacteria (40%), Cyanobacteria (13%), Chloroflexi (11%), Firmicutes (7%), Thermotogae (6%), Bacteroidetes (3%), Acidobacteria and Chlorobi (2%) and the most represented genera were: cyanobacteria *Leptolyngbya* (8%), aerobic heterotroph *Chondromyces* (7%), sulphur reducing/anoxygenic heterotroph *Marinitoga* (5%), anoxygenic phototrophs *Gloeotrichia* and *Roseiflexus*, *Oscillochloris*, aerobic chemoheterotrophic *Sphingomonas*, sulphate reducing *Desulfobacca*, hydrogen oxidizing *Hydrogenophilus*, and sulphur oxidizing *Thiofaba* (1%).

It is uncertain whether the Bullicame streamers are formed by the sulphur oxidizing *Thiofaba* identified by Valeriani et al. (2018) or by the Chloroflexi anoxygenic phototrophs as suggested by Folk (1994), Allen et al. (2000) and Pentecost and Coletta (2007). The Aquificales microbes identified at Yellowstone hot springs at temperature higher than 60°C seem to have a broad niche and a wide range of environmental tolerance (Fouke, 2011) but there is no evidence they could dominate also in the lower temperature (50-55°C) conditions of the Bullicame channel. The genus *Sulfurihydrogenibium* tends to be the predominant member of the Aquificales group in terrestrial springs with pH near neutral and elevated sulphide concentrations (Flores et al., 2008). Nevertheless, there are various species of *Sulfurihydrogenibium* and some grow in temperature intervals of 40-70°C (Nakagawa et al., 2005; Flores et al., 2008). The sulphide oxidizing bacteria *Thiothrix* has been identified also in travertines at 24°C associated with *Beggiatoa*, purple sulphur and sulphate reducing bacteria (Bonny and Jones, 2008). *Thiofaba tepidiphila* identified by Valeriani et al. (2018) at Bullicame is a chemolithoautotrophic sulphur oxidizing bacterium of the Gammaproteobacteria, family Halothiobacillaceae, isolated from a hot spring in Japan at temperature of 45°C and pH 7.0 (Mori and Suzuki, 2008). Both *Thiofaba* and Aquificales exhibit a rod-shaped morphology and this contrasts with the observed filamentous streamers. Nevertheless, it seems that, according to the environmental conditions and permanent water flow, Aquificales can develop macroscopic filaments or aggregates (Eder and Huber, 2002; Alain et al., 2003). Kubo et al. (2011) observed that at the Nakabusa hot spring in Japan at 65°C, three thermophilic bacteria groups (sulphide oxidizers, anoxygenic phototrophs, sulphate reducers) occur spatially distributed controlling the sulphur cycle in the microbial mat. The aerobic chemolithotrophic sulphide-oxidizing *Sulfurihydrogenibium* dominated near the mat surface, while the filamentous anoxygenic photosynthetic *Chloroflexus* was in deeper layers. Sulphide was produced by the anaerobic sulphate reducing *Thermodesulfobacterium*/*Thermodesulfatator* under anoxic-dark conditions, while sulphide was consumed by the *Chloroflexus* anoxygenic photosynthetic bacteria under anoxic-light conditions and strong sulphide oxidation took place by the chemolithotrophic members of the Aquificales under oxic-dark conditions. Kubo et al. (2011) proposed the intimate association between rod-shaped sulphide oxidizers and filamentous anoxygenic phototrophs where Aquificales act as highly efficient scavengers of oxygen from the spring water, creating a favourable anoxic environment for *Chloroflexus* and sulphate reducers in deeper layers.

Hence, the proximal channel streamers at Bullicame might be formed by carbonate precipitation encrusting microbial mats from either sulphide oxidizing bacteria or anoxygenic phototrophs or an association of both. The purple microbial mats in the proximal channel could represent purple bacteria, which also are anoxygenic phototrophs (Reysenbach and Cady, 2001). Purple sulphur bacteria (*Chromatium*, *Thiospirillum*, *Thiocapsa*) utilize bacteriophylls with red and purple carotenoid pigments. They often overlie sulphate reducing bacteria that produce H_2_S; some can grow also in the presence of O_2_ (Konhauser, 2007). *Chloroflexus* is often associated with purple sulphur bacteria and under fully aerobic conditions bacteriophyll synthesis is repressed, the organism grows chemoheterotrophically and the colour of the culture changes from dull green to orange (Norris et al., 2002; Konhauser, 2007; Dunckel et al., 2009). This may suggest that the orange microbial mats of the Bullicame proximal channel floor might represent *Chloroflexus* mats. The sparse rod-shaped microbes observed could be sulphate reducing bacteria as suggested by the identification of *Desulfobacca* by Valeriani et al. (2018), or the thermophilic sulphur reducing bacterium *Desulphovibrio thermophilus* (Allen et al., 2000), or the rod-shaped thermophilic cyanobacteria *Synechococcus* as reported in numerous travertine microbial mats (Pentecost, 2003).

The channel margins and distal channel, at temperatures < 50°C, are characterized by filamentous microbes, dominated by spiral-shaped forms attributed to *Spirulina labyrinthiformis* (cf. Pentecost, 2003), associated with segmented filamentous and coccoid microbes that could belong to *Lyngbya limnetica* or *Phormidium* (cf. Di Benedetto et al., 2011) or to *Leptolyngbya* (cf. Valeriani et al., 2018; Gong et al., 2020) and to *Synechococcus* (Pentecost, 2003), respectively. Pentecost and Coletta (2007) identified *Spirulina labyrinthiformis* as the most abundant cyanobacteria from temperatures < 54°C, associated with *Fischerella laminosus*, *Oscillatoria* cf. *geminata* and *Phormidium laminosum*. Diatoms were present only occasionally and the desmid green alga *Cosmarium leave* was found at temperatures below 50°C.

The cyanobacterium *Fischerella laminosus* seems to live at temperatures of 47-35°C, whereas *Oscillatoria formosa* and *Phormidium laminosum* occur at 56-40°C (Pentecost, 2003). Di Benedetto et al. (2011) in the microbial mat at Bullicame 3 vent, located half way between Bullicame mound and Le Zitelle spring, determined cyanobacteria associations varying as a function of water temperature: 1) the high temperature (57°C) cyanobacteria were *Synechococcus eximius*, *Chroococcus* cf. *yellowstonensis*; 2) the intermediate temperature (47°C) forms were *Synechococcus eximius, Lyngbia limnetica*, *Phormidium*, 3) at 37°C *Synechococcus eximius*, *Oscillatoria*, diatoms and Chlorophyta *Dyctiospherium*, *Scenedesmus acuminata*; 4) the coolest temperature (27°C) cyanobacteria included *Nostoc comune*, *Anabaena circinalis*, *Oscillatoria*, associated with diatoms and green algae.

#### 5.2.2 Bollore travertine facies and microbial mats

At Bollore, travertine precipitates vary downstream at the macro- and mesoscale with streamers at the vent and proximal channel (49-44°C), and centimetre-size terraced coated gas bubbles, dendrites, laminae and rafts in the distal channel (44-40°C). These distal channel fabrics are characteristics of numerous fossil travertines (Chafetz and Folk, 1984; Guo and Riding, 1998; Gandin and Capezzuoli, 2014; Della Porta et al., 2017ab). As for Bullicame, at the microscale, the carbonate precipitates are similar from proximal to distal settings with microsparite/fine sparite spherulites embedded within and overlying microbial mats. The EPS act as substrate for crystal nucleation controlling the spatial distribution of calcite spherulites and carbonate fabrics at the mesoscale. Aragonite spherulites occur only in the highest temperature proximal setting, as in Bullicame, and crystal size decreases distally. At Bollore, the high temperature (49-44°C) microbial mats consist of filamentous, rod-shaped and coccoid microbes and rare *Spirulina* cyanobacteria, whereas the low temperature (44-40°C) assemblage shows dominant *Spirulina* and other filamentous forms. Similarly to Bullicame, the Bollore streamers might be formed either by sulphide oxidizing microbes or Chloroflexi anoxygenic phototrophs or an association of both (cf. Kubo et al., 2011). The rod-shaped microbes can represent sulphate reducing bacteria or the cyanobacterium *Synechococcus*, which tolerates the highest temperature conditions (73°C) for oxygenic photosynthesis (Cady and Farmer, 1996; Pentecost, 2003). The lower temperature, distal channel microbial mats are dominated by *Spirulina* and other filamentous cyanobacteria, possibly *Phormidium* and *Oscillatoria*. Pentecost (2003) identified at temperature of 40-38°C the filamentous cyanobacteria *Schizotrix perforans* and the coccoid *Aphanocapsa thermalis*. At the nearby Bagno Vignoni hydrothermal system (nearly 17 km far from Bollore), the cyanobacteria identified at temperatures of 43-34°C were *Spirulina labyrinthiformins*, *Phormidium laminosum*, *Lyngbya* sp., *Pseudoanabaena* and *Synechococcus elongatus* (Pentecost, 1994, 2003).

#### 5.2.3 Gorello Waterfall travertine facies and microbial mats

The Gorello Waterfall (33-34°C) carbonate precipitates consist of microsparite/sparite spherulites as at Bullicame and Bollore but micron- and nano-metre scale micritic precipitates are common and aragonite is absent. Carbonate crystal spherulites precipitate exclusively on the organic substrates provided by the microbial mats, spatially distributed in an ordered reticulate or laminated framework, as in the other two case studies but here also filamentous microbes are encrusted by carbonate crystals. The microbial mats include a diverse community of filamentous cyanobacteria with various trichome sizes. Bazzichelli et al. (1978) identified in the Saturnia thermal spring cyanobacteria (*Pseudanabaena ulula*, *Oscillatoria boryana*, *Synechococcus eximius*) and sulphur bacteria. In addition to *Spirulina labyrinthiformis*, the filamentous microbes with a diameter of 1 μm might belong to *Oscillatoria*, *Phormidium*, or *Schizotrix* following the diagnostic criteria proposed by Pentecost (2003). The wide filamentous microbes (4-5 μm in diameter; Figure S12) might belong to *Oscillatoria*, *Phormidium*, *Leptolyngbya, Calothrix thermalis* or *Scytonema*. The chain-like forms resemble the cyanobacteria *Nostoc*, *Anabaena* or *Fischerella* (cf. Pentecost, 2003). Most of these inferred cyanobacteria forms have been identified in moderate temperature (20-50°C) hydrothermal systems associated with diatoms (Pentecost, 2003; Shiraishi et al., 2019; Gong et al., 2020). In particular, *Calothrix thermalis* has been described in various hydrothermal vents with temperature of 20-40°C (Cady and Farmer, 1996; Farmer, 2000; Pentecost, 2003, 2005; Jones and Peng, 2015). Alternatively, some segmented and chain-like forms resemble the hyphae of Actinomycetes described in cave deposits (Jones, 2009).

### 5.3 Travertine carbon and oxygen stable isotope signature

The measured δ^13^C and δ^18^O fall (Figure 11) within the field of stable isotope signature for hydrothermal travertine in Central Italy (Minissale et al., 2002ab; Minissale, 2004; Gandin and Capezzuoli, 2008; Della Porta, 2015; Kele et al., 2015). Travertine δ^13^C and δ^18^O values reflect the isotopic composition of thermal water, water temperature, the origin of the dissolved inorganic carbon (DIC) from decarbonation of Mesozoic limestone in contact with magmatic fluids and kinetic fractionation due to CO_2_ degassing and fast carbonate precipitation rates (Fouke et al., 2000; Minissale, 2004; Pentecost, 2005; Kele et al., 2015; Della Porta, 2015; Della Porta et al., 2017ab and references therein). The Bullicame and Bollore streamers record the lowest δ^18^O and δ^13^C values probably because streamers precipitate in the proximal setting, where water temperature is higher and degassing of CO_2_, removing light ^12^CO_2_ has not been prolonged. An alternative explanation is that the streamer δ^13^C is affected by microbial metabolism with predominant effect of sulphide oxidation, anoxygenic photosynthesis or microbial respiration in the proximal setting and effect of photosynthesis in the distal setting as suggested for some travertine case studies (Guo et al., 1996; Fouke et al., 2000; Reysenbach and Cady, 2001; Zhang et al., 2004). This effect of microbial metabolism on the carbon stable isotopes seems, however, unlikely because the physico-chemical processes (CO_2_ degassing and water cooling) would produce isotopic shifts of larger magnitude that might override any possible microbial influence on the fractionation of carbon stable isotopes (cf. discussion in Fouke et al., 2000; Fouke, 2011; Della Porta, 2015). In addition, carbonate precipitation taking place through EPS-mediated mineralization does not produce enzymatic fractionation (Reitner, 1993; Reitner et al., 1995b, 2000).

### 5.4 Microbial mats and EPS-mediated carbonate mineralization

This study identifies three temperature-controlled microbial communities and related carbonate facies: a) high temperature (50-55°C in Bullicame, 44-49°C in Bollore) streamers with sulphide oxidizing bacteria and/or green non sulphur *Chloroflexus* anoxygenic phototrophs associated with sulphate reducing bacteria and rod-shaped cyanobacteria *Synechococcus*; b) intermediate temperature (<50°C in Bullicame, 40-44°C in Bollore) crystalline and clotted peloidal micrite dendrites, rafts and coated bubbles with cyanobacteria dominated by *Spirulina*, possible *Synechococcus* and other filamentous Oscillatoriales (possible *Oscillatoria*, *Phormidium*); c) low temperature (34-33°C in Gorello Waterfall) laminated boundstone and coated grains with high diversity of Oscillatoriales filamentous cyanobacteria (possible *Oscillatoria*, *Phormidium*, *Lyngbya*, *Leptolyngbya*, *Schizotrix*) associated with *Spirulina*, possible *Nostocales* (*Nostoc*, *Anabaena*, *Pseudoanabaena*, *Calothrix*, *Fischerella*), and diatoms as observed in the numerous reported studies about hydrothermal terrestrial spring deposits (cf. Pentecost and Tortora, 1989; Ward et al., 1998; Pentecost, 2003; Norris et al., 2002; Sompong et al., 2005; Norris and Castenholz, 2006; Pentecost et al., 2007; Roeselers et al., 2007; Okumura et al., 2013; Roy et al., 2014; Smythe et al., 2016; Sugihara et al., 2016; Shiraishi et al., 2019). These temperature controlled microbial communities are similar to those proposed by various authors for the Yellowstone National Park springs (Cady and Farmer, 1996; Farmer; 2000; Fouke, 2011; Des Marais and Walter, 2019). Farmer (2000) proposed that the three temperature-controlled geomicrobiological facies from Yellowstone reflect the inferred sequence of evolutionary events implied by the RNA universal phylogenetic tree with chemolithoautotrophic sulphide oxidizers, anoxygenic phototroph *Chloroflexus* and then the oxygenic cyanobacteria (*Synechococcus*, *Spirulina*) followed by cyanobacteria as *Calothrix* and eukaryotes with diatoms and grazing insects at lowest temperature.

In the studied travertines, the temperature control on the composition of the microbial mats is paralleled by a change in carbonate facies at the mesoscale with proximal higher temperature streamers to distal lower temperature laminated boundstone, rafts, coated gas bubbles and clotted peloidal micrite dendrites. However, at the microscale the carbonate precipitates are similarly consisting of calcite and/or aragonite crystal spherulites nucleated on/within the microbial mat independently from the microbial metabolic pathways. It appears that microbial mats exert an influence on carbonate precipitates by acting as substrates for mineral nucleation rather than inducing carbonate precipitation through microbial metabolism. Carbonate supersaturation must be achieved through physico-chemical processes, primarily CO_2_ degassing of neutral pH and high alkalinity waters, whereas microbial mats provide passive substrates for crystal nucleation and influence the characteristics of carbonate fabrics controlling the spatial distribution of carbonate crystals. Crystal spherulites are distributed in ordered reticulate and laminated framework structures mimicking the organic template that act as substrates for crystal nucleation. These evidences suggest that carbonate precipitation in the investigated hydrothermal travertines is largely the result of biofilm EPS-mediated mineralization (cf. Reitner, 1993; Défarge and Trichet, 1995; Reitner et al., 1995a, 2001; Arp et al., 1999, 2003, 2012; Ionescu et al., 2014).

## 6 Conclusions

The three investigated present-day hydrothermal travertine settings differ for morphology, water chemistry and temperature. At the microscale, however, carbonate precipitates are dominated by microsparite/fine sparite crystals organised in radial spherulitic structures within or above microbial organic substrates. Despite a similarity of the microscale carbonate precipitates, the microbial communities vary with water temperature in the three systems. Three microbial mat associations can be distinguished: a) a proximal higher temperature association (55-44°C) of filamentous and rod-shaped sulphide oxidizing bacteria, sulphate reducing and anoxygenic phototrophic bacteria forming streamer fabrics; b) an intermediate temperature (44-40°C) association dominated by *Spirulina* cyanobacteria with rafts, dendrites and coated gas bubble fabrics, c) a lower temperature (40-33°C) microbial community of filamentous Oscillatoriales, Nostocales and *Spirulina* cyanobacteria and diatoms occurring within laminated boundstone and coated grains. In terms of mineralogy, calcite is the predominant mineral, whereas aragonite occurs only in the hottest water at temperature > 44°C. Gypsum, Ca-phosphate, authigenic aluminium-silicate minerals and impregnation of EPS by Si, Al, Ca and Mg or P and S can occur. Stable carbon and oxygen isotope values are similar to other travertines in Central Italy and reflect the isotopic composition of thermal water, temperature, source of CO_2_ and distance from the vent due to progressive CO_2_ degassing, cooling, and carbonate precipitation.

This study confirms that microbial communities in terrestrial hydrothermal travertine systems vary as a function of water temperature but that the microscale precipitates are similarly consisting of carbonate crystals forming spherulites distributed in an ordered framework reflecting the organic substrate template. This evidence suggests that microbial metabolism does not play a relevant role in controlling the type of microscale carbonate precipitates. Carbonate precipitation in hydrothermal settings, driven by thermal water physico-chemical processes increasing carbonate supersaturation, is influenced by the microbial biofilm EPS (EPS-mediated mineralization) acting as low-energy template for crystal nucleation, controlling the spatial distribution of carbonate crystals and influencing the mesoscale fabrics.

## Supplementary Materials

The following are available online at www.mdpi.com/xxx/s1, Figures S1-S12, Table S1.

## Author Contributions

Conceptualization, Data curation: G.D.P. and J.R. Formal analysis, G.D.P.; Funding acquisition, G.D.P. and J.R.; Methodology, G.D.P. and J.R.; Supervision, J.R.; Writing-original draft, G.D.P.; Writing-review & editing, G.D.P. and J.R. All authors have read and agreed to the published version of the manuscript.

## Funding

This work was supported by DAAD (Deutscher Akademischer Austauschdient) for funding the research stay of Giovanna Della Porta at the Geobiology, Göttingen Centre of Geosciences, Georg-August-University Göttingen (Germany). Joachim Reitner received financial support from the German Research Council (DFG) DFG-For 571 “Geobiology of Organo- and Biofilms”.

## Acknowledgments

Wolfgang Dröse is warmly thanked for all the assistance in sample preparation at the Geobiology Laboratory of the University Göttingen. Dorothea Hause-Reitner and Agostino Rizzi are warmly thanked for the assistance in the SEM analyses at the University of Göttingen and University of Milan, respectively. Elena Ferrari is thanked for support in the stable isotope analyses at the Earth Sciences Department of the University of Milan, Miriam Ascagni for the confocal laser scanning microscope analyses at the laboratories Unitech of the University of Milan. Enrico Capezzuoli is thanked for fundamental help during fieldwork and stimulating discussions.

## Conflict of Interest

The authors declare no conflict of interest. The funders had no role in the design of the study, in the collection, analyses, or interpretation of data, in the writing of the manuscript, or in the decision to publish the results.

## Supplementary materials

**Figure S1.**
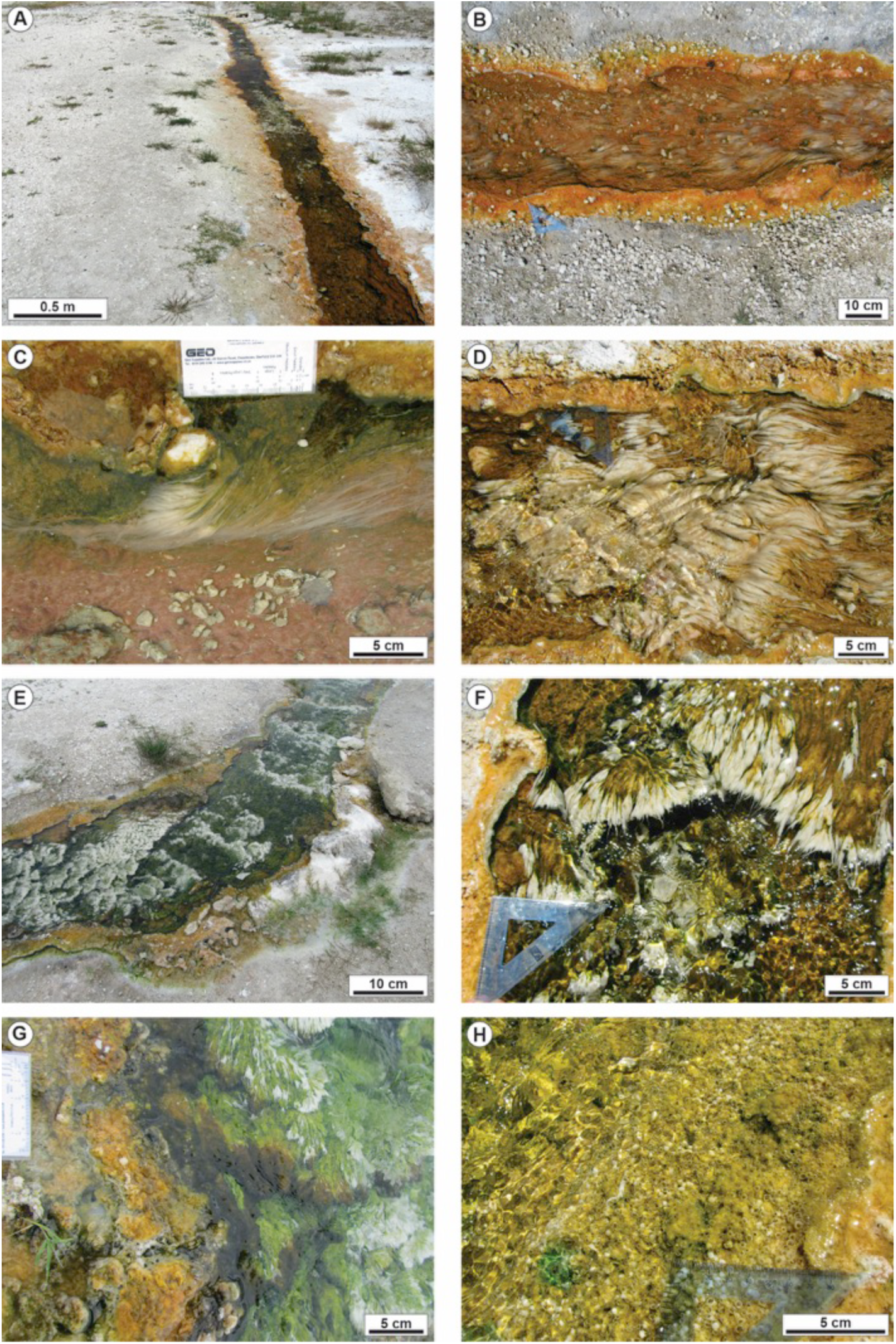
Bullicame (Viterbo) active hydrothermal travertine system. A-B-C) Proximal channel, from 12.5 m to 20 m from the vent centre, draped by purple microbial mats in the centre, with sparse bundles of white filaments oriented according to the current flow direction and with orange/yellow to green microbial mats on the channel margin. D-E-F) Proximal channel in the stretch from 20 m to 40 m from the vent centre with abundant bundles of carbonate-coated filaments, the channel floor draped by microbial mats that change in colour from purple to green while the channel margins are orange/yellow to green in colour with gas bubbles. G) Proximal channel at 30 m from the vent with channel margin characterized by yellow/orange microbial mats with coated gas bubbles and the channel centre rich of white filamentous bundles coated by green to brown microbial mat. H) Distal channel at 60 m from the vent centre, with centre and margin draped by orange/yellow to green microbial mat with abundant carbonate-coated gas bubbles and rafts.

**Figure S2.**
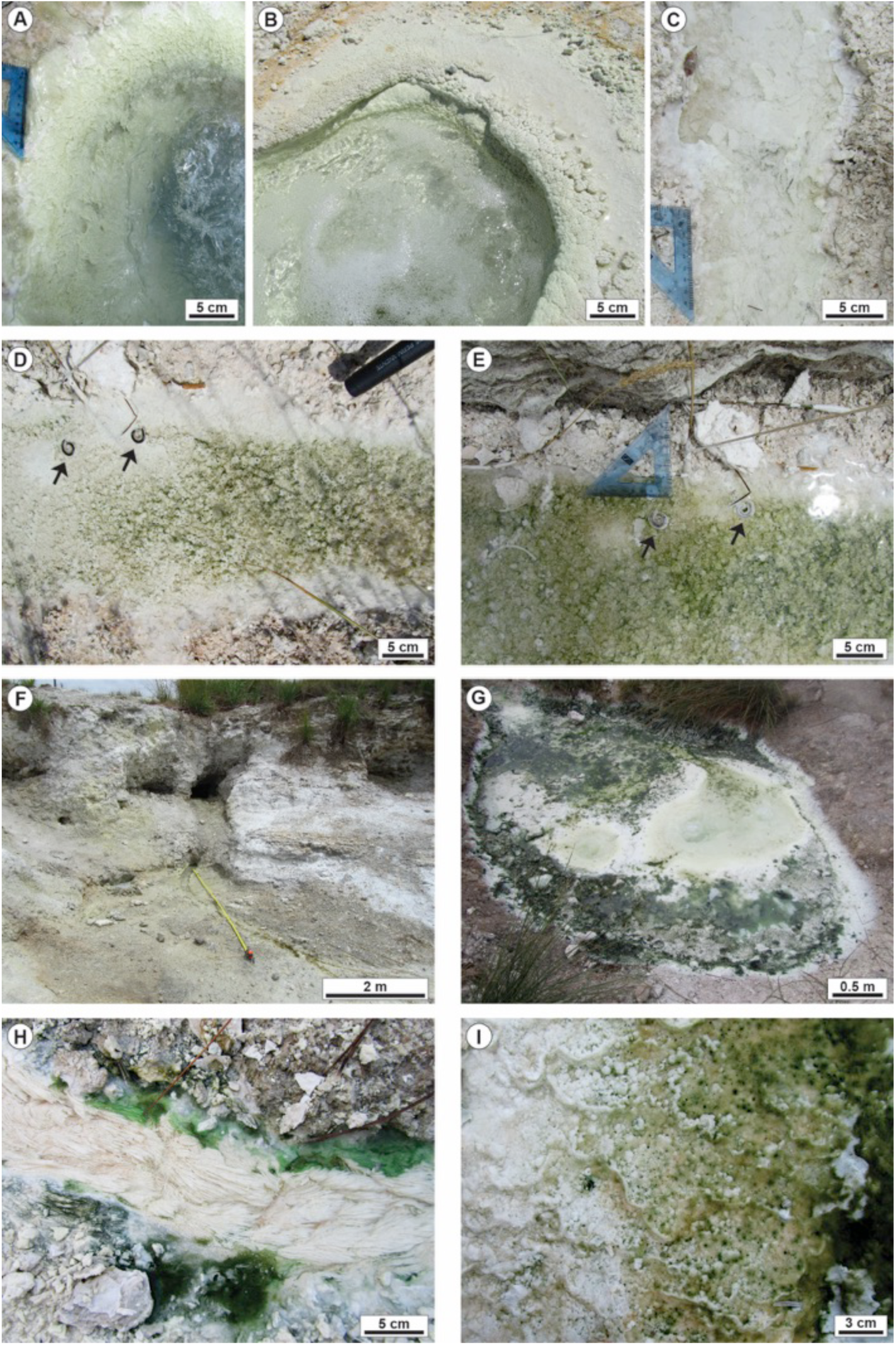
Bollore (Bagni San Filippo) active hydrothermal travertine system. A-B) Close-up views of the vent orifices with the pool rims encrusted by carbonate as coated filaments and dendrites. C) Proximal channel with white carbonate-encrusted bundles of filaments and paper thin rafts. D-E) Distal channel 12 m from the vent with the channel floor draped by green microbial mat with abundant carbonate-coated gas bubbles. The two images where taken during two following days: to notice that the two dead worms (black arrows) are not coated by carbonate in Figure S2D whereas are carbonate-coated in the image in Figure S2E taken the following day. This confirms the fast rates of carbonate precipitation producing millimetres-thick carbonate coatings in 24 hours. F) Second vent and channel in a lower topographic position. The yellow colour of the sediment is provided by abundant sulphur draping the precipitated travertines. G-H-I) During the winter humid season (January), the main Bollore vent and channel thrive with dark green microbial mats, whereas during the summer sampling (July) the green microbial mats were lacking and the channel floor colour was white to light pink.

**Figure S3.**
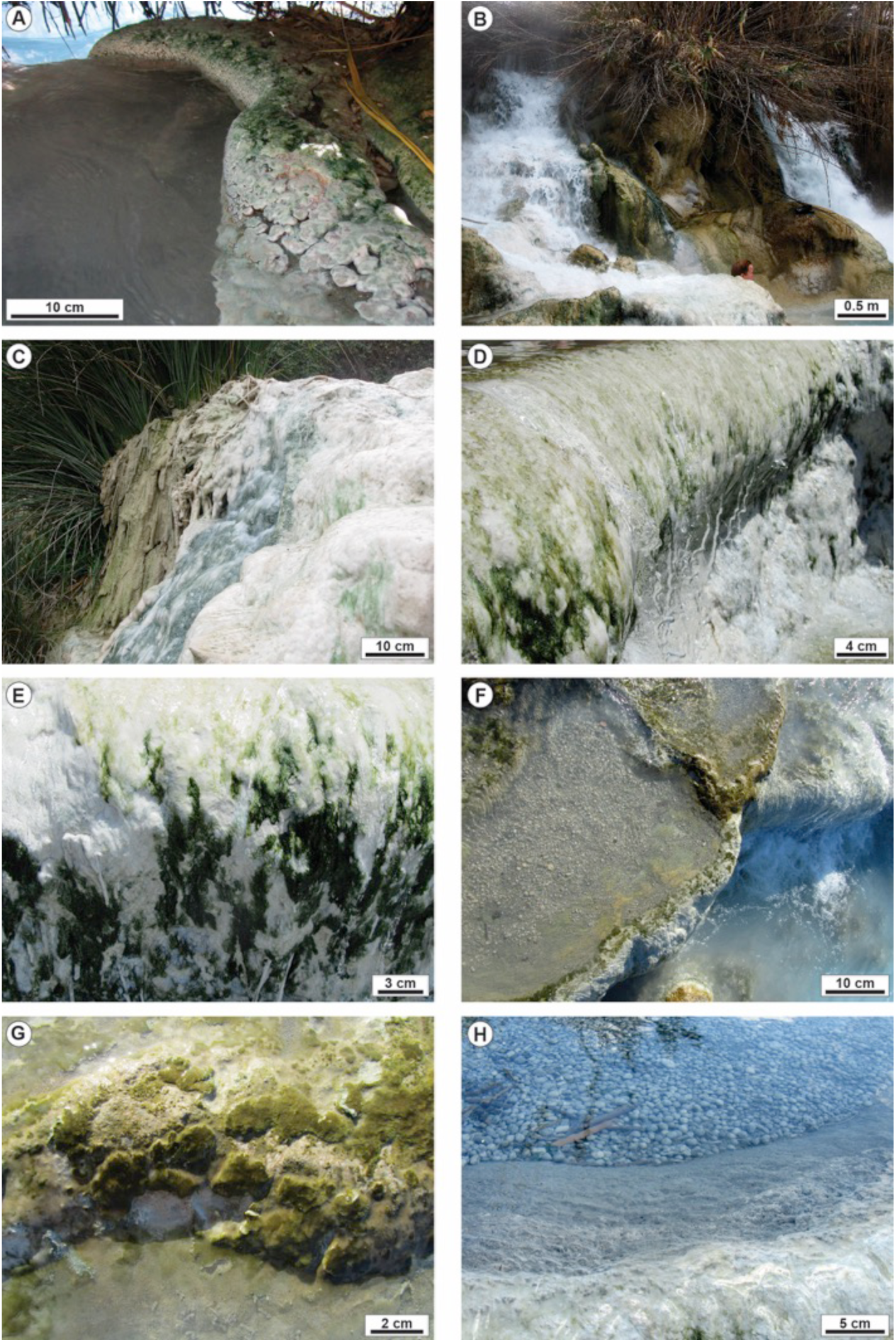
Gorello Waterfall (Saturnia) active hydrothermal travertine system. A) Photograph showing the channel deriving from the thermal spa before the break in slope that generates the waterfall. The channel levees are draped by green to pink microbial mats. B) The nearly 5 m high waterfall developed at the channel topographic break in slope. C) Areas of the pool rims and walls not flooded by thermal water are temporarily colonized by reed vegetation coated by carbonate precipitates when thermal water flow is resumed. D-E) The vertical walls of the pools of the terraced slope are coated by white to dark green microbial mat. F-G) The rims of the pools of the terraced slope are coated by olive green microbial mat. H) Centimetre-size carbonate coated grains (oncoids) forming on the pool floor.

**Figure S4.**
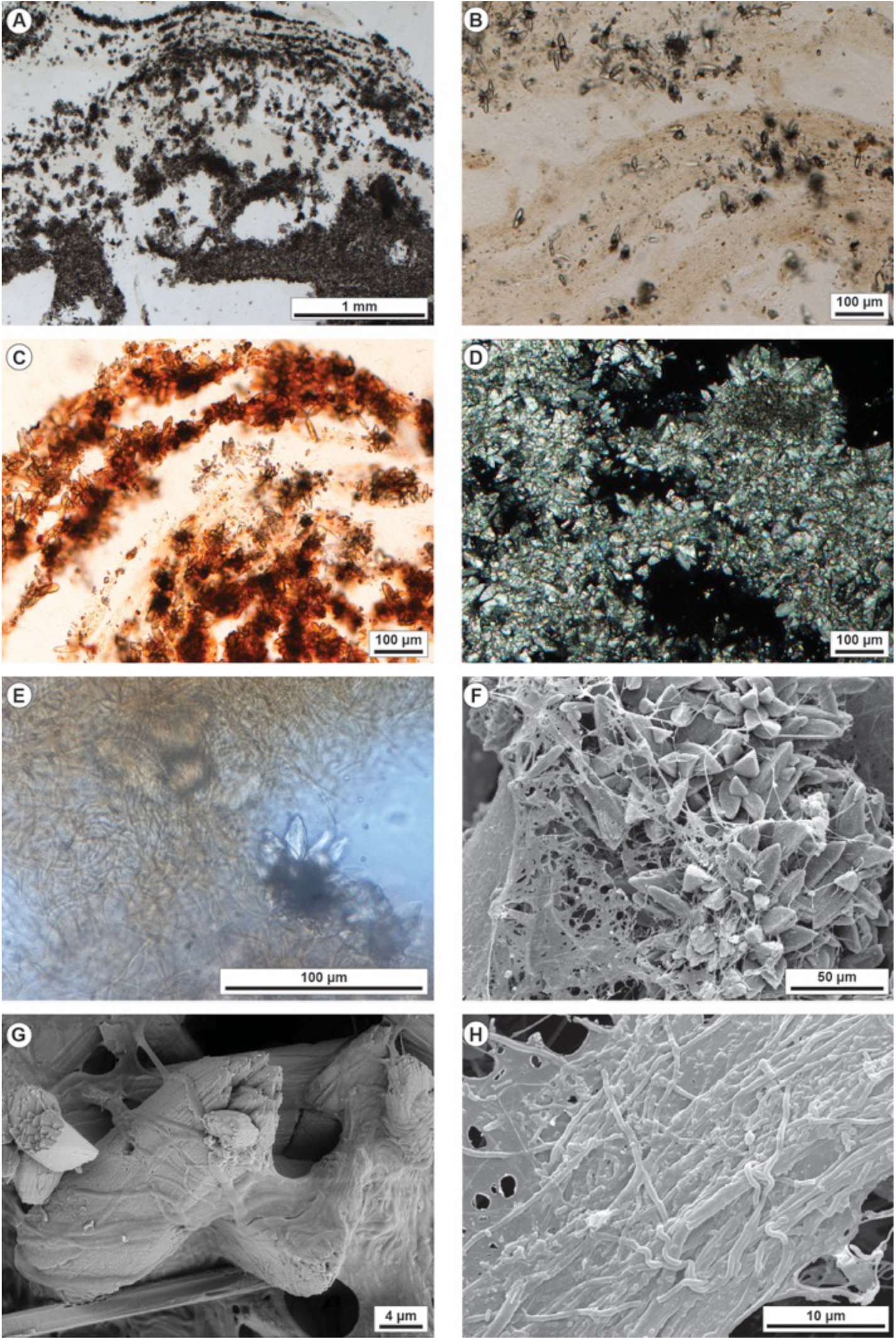
Petrographic and SEM images of the Bullicame travertines in the proximal channel. A) Carbonate precipitates as microsparite/sparite forming spherulites within and above organic substrates. The spatial distribution of the carbonate precipitates forming laminae and irregular mosaics is controlled by the framework of the EPS. B) Calcite crystal spherulites embedded within microbial mat following the shape and geometry of the organic substrate. C) Calcein-dyed sample showing that the calcite crystal spherulites mimic the shape and geometry of the organic substrate forming undulated laminae. D) Photomicrograph in crossed-polarisers showing the aggregates of microsparite crystals and the sparse clots of micrite mostly at spherulite nuclei. E) Close-up view of calcite spherulites surrounded by filamentous microbes and EPS. F) SEM image showing the rosettes of euhedral prismatic calcite crystals embedded in EPS with filamentous microbes. G) Close-up view at SEM of prismatic calcite crystals embedded in EPS and filamentous microbes. At the bottom a Ca phosphate, probably apatite, crystal. H) SEM image of the microbial mat in the proximal channel consisting of bundles of filamentous microbes. EPS include also rod-shaped microbes (upper left corner).

**Figure S5.**
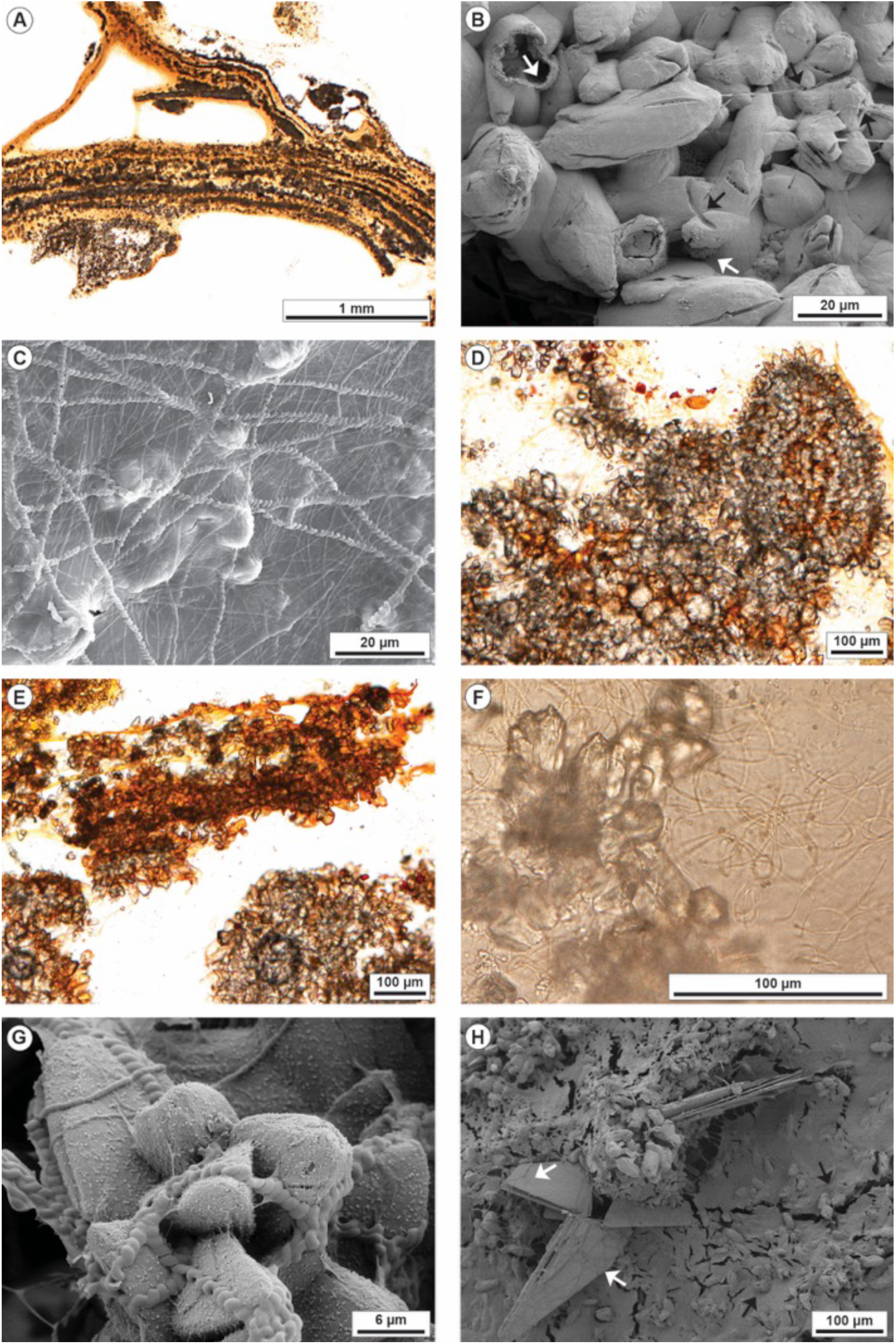
Petrographic and SEM images of the Bullicame travertines in the proximal channel margins (A-C) and distal channel (D-H). A) Photomicrograph of tetracycline-dyed sample showing how the calcite crystal spherulites precipitate within the microbial mat following the framework and lamination of the organic substrate. B) SEM image of spindle-shaped microsparite crystals showing tubular perforations likely related to moulds of filamentous microbes entombed during crystal growth because some organic filaments are emerging from these tubular hollows (black arrows). The image shows also acicular aragonite hollow spheres (white arrows). C) SEM image of EPS surrounding the calcite crystals embedding *Spirulina* cyanobacteria and other filamentous microbes. D) Photomicrograph of calcein-dyed sample showing the framework of aggregates of microsparite spherulites associated with micrite clots and laminae. E) Photomicrograph of calcein-dyed sample showing the framework of aggregates of microsparite spherulites associated with micrite clots and laminae impregnated by organic matter of the microbial mat. F) Close-up view of the microsparite spherulites with a micrite nucleus surrounded by EPS embedding filamentous microbes including *Spirulina* cyanobacteria. G) SEM image of calcite crystals draped by *Spirulina* cyanobacteria and other filamentous microbes and by a grumous mucilaginous organic material. H) SEM image showing EPS embedding calcite crystals (black arrows) and Ca phosphate crystals coated by filamentous microbes (white arrows).

**Figure S6.**
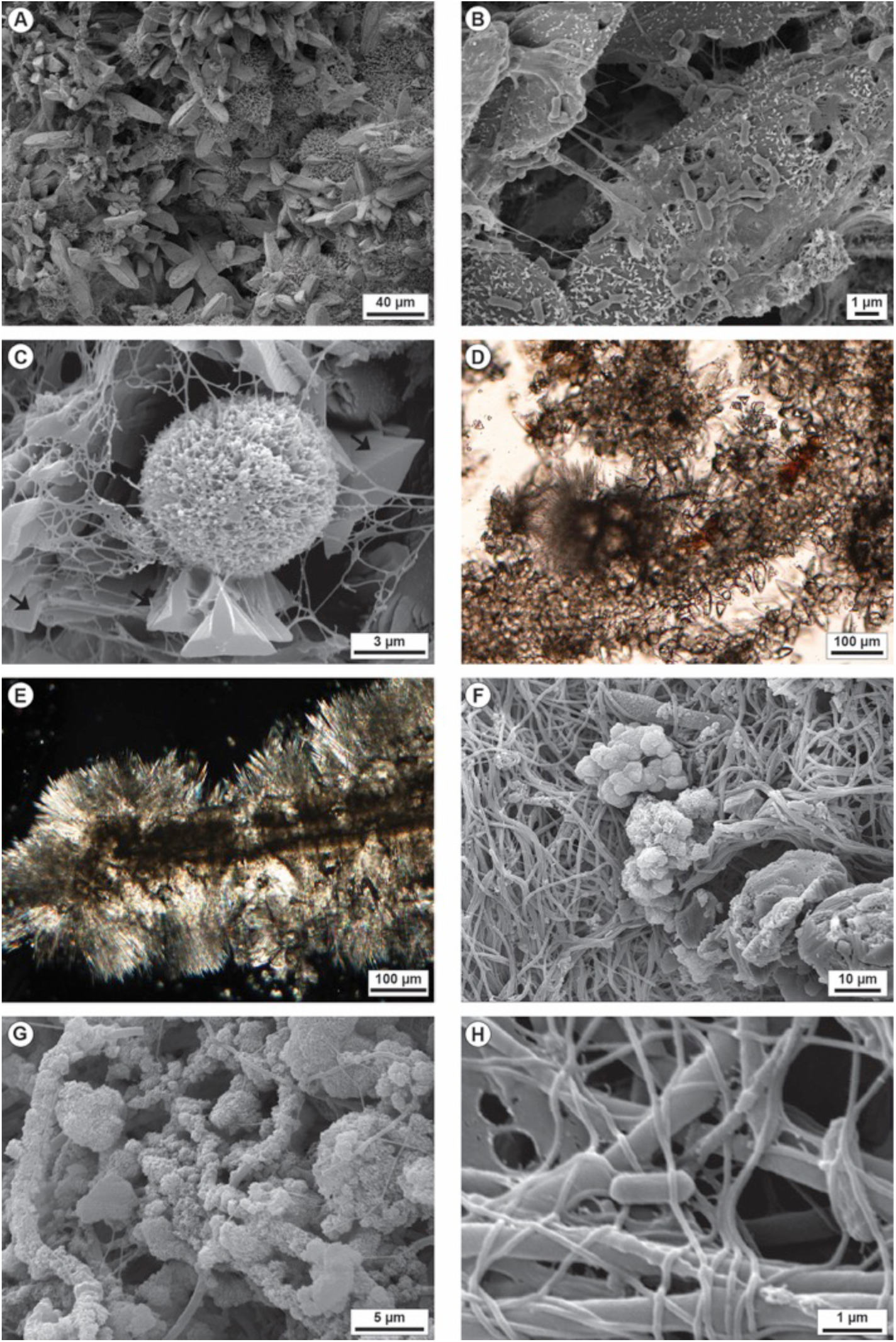
Petrographic and SEM images of the Bollore (Bagni San Filippo) travertines at the main vent (A-C) and second vent (D-H). A) SEM image showing the prismatic calcite crystal forming radial rosettes and the acicular aragonite spherulites that appear to postdate calcite precipitation. B) Calcite crystals embedded in EPS and overlain by rod-shaped microbes. C) SEM image of acicular aragonite spherulites surrounded by gypsum crystals (black arrows) all draped by EPS. D) Carbonate fabric of microsparite spherulites forming aggregates and laminae followed by acicular aragonite spherulites. E) Crossed-polarizers image of micritic and microsparitic laminae surrounded by acicular aragonite crystal fans. F) Bundles of filamentous microbes overlain by aggregates of possible authigenic clay mineral. G) SEM image showing the filamentous microbes coated by authigenic silicate. H) SEM image showing entangled filamentous microbes with different size in cross section and rod-shaped microbes.

**Figure S7.**
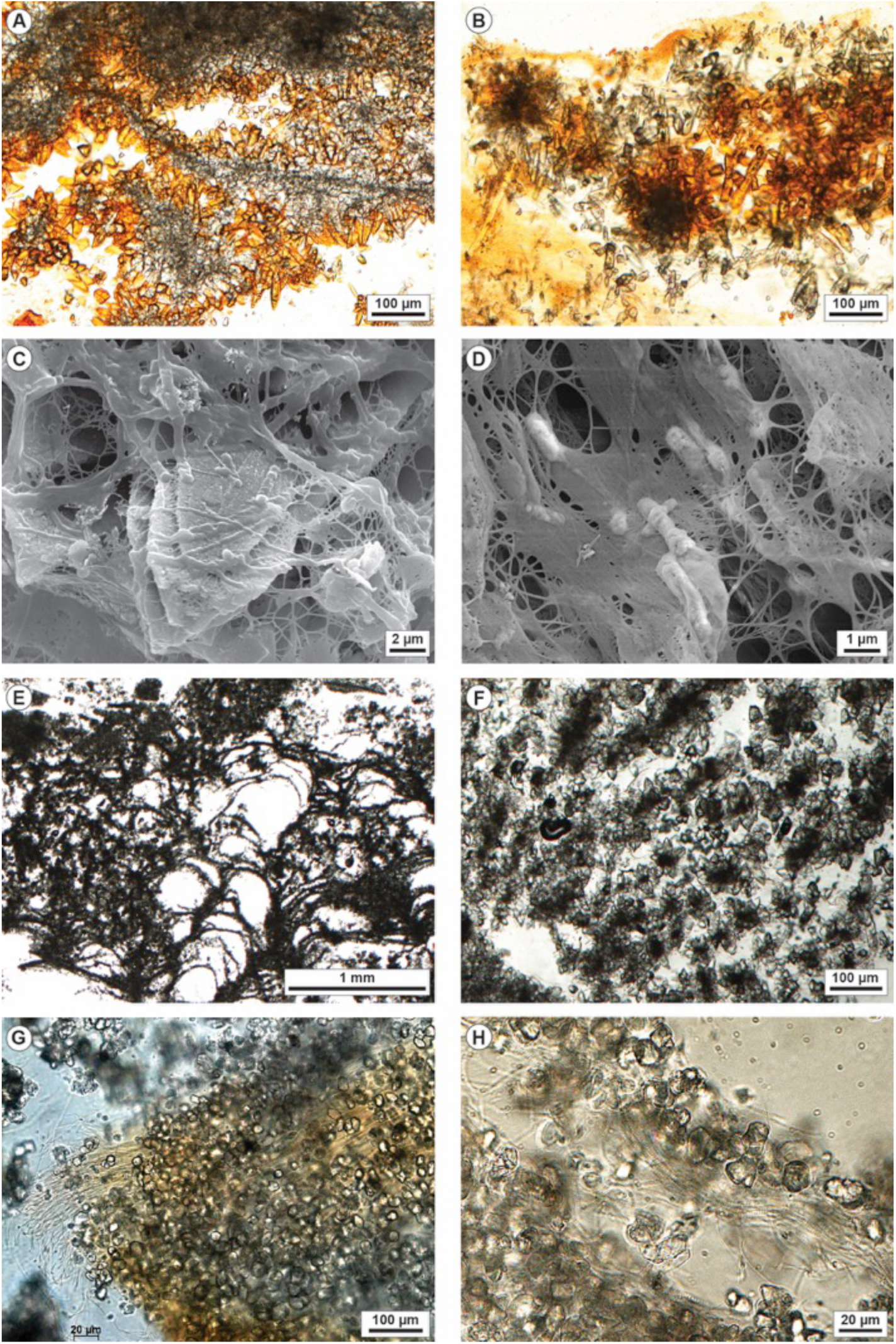
Petrographic and SEM images of the Bollore (Bagni San Filippo) travertines at the proximal (A-D) and distal channel (E-F). A) Micrite filaments and clots surrounded by microsparite to fine sparite crystals departing radially from the micritic substrate. Crystals are stained by orange calcein dye as they were coated by organic matter. B) Calcite crystal spherulites embedded in calcein dyed microbial mat. Some rosettes have the nuclei made of micrite clots. C) SEM image of prismatic calcite crystals with gothic-arch shape surrounded by EPS embedding filamentous and rod-shaped microbes. D) SEM image of EPS embedding rod-shaped microbes. E) Photomicrograph of distal channel coated bubble boundstone made of clotted peloidal micrite. F) Photomicrograph of a framework of calcite crystal spherulites with micrite nuclei aligned and embedded in organic matter that must sustain the crystal framework. G) Microsparite crystals embedded in EPS and filamentous microbes. H) Microsparite crystals surrounded by filamentous microbes with abundant *Spirulina* cyanobacteria.

**Figure S8.**
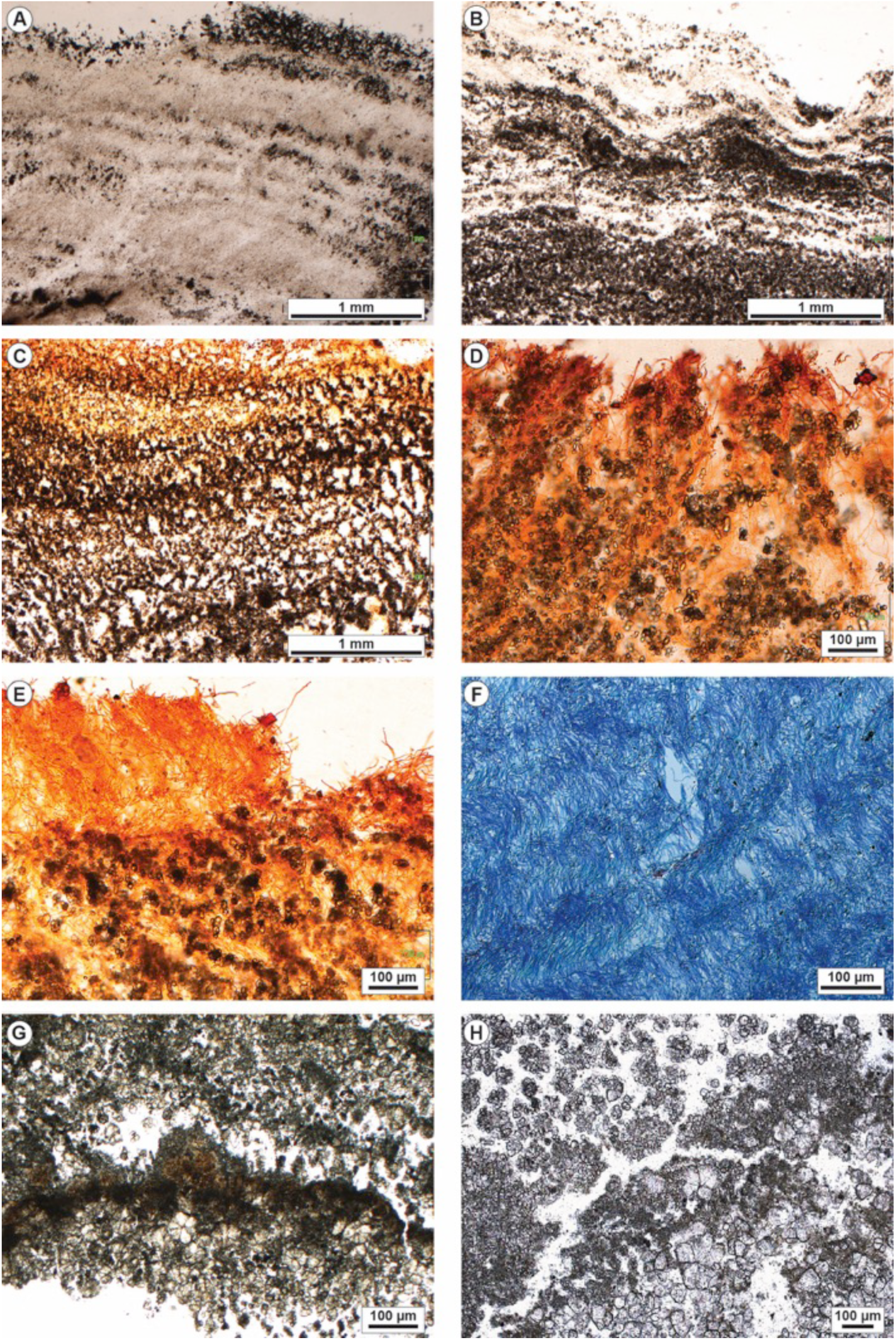
Petrographic analysis images of the Gorello Waterfall (Saturnia) travertines from the rims and walls of the terraced slope system. A) Photomicrograph showing the micritic laminated boundstone with alternation of carbonate precipitates and microbial mat that acted as organic matter substrate for carbonate precipitation. B) Laminated boundstone showing the wavy lamination made of precipitated carbonate alternating with organic matter from the microbial mat. C) Calcein dyed laminated boundstone showing that carbonate crystals precipitated following the alveolar fabric of the biofilm EPS. D) Calcein dyed sample showing the microsparite crystals and spherulites precipitating in between the erect filamentous microbes making the outer layer of the microbial mat. E) Calcein dyed sample showing the erect filamentous microbes making the outer layer of the microbial mat with below micrite and microsparite precipitated following the spatial distribution of the filamentous microbes. F) Paraffin sample dyed with alcian blue showing the geometry of filamentous microbes in the microbial mats that is mimicked by the laminated carbonate precipitates. G) Laminated boundstone with microsparite/sparite spherulites and clotted peloidal micrite laminae. H) Framework made of microsparite/sparite spherulites and clotted peloidal micrite.

**Figure S9.**
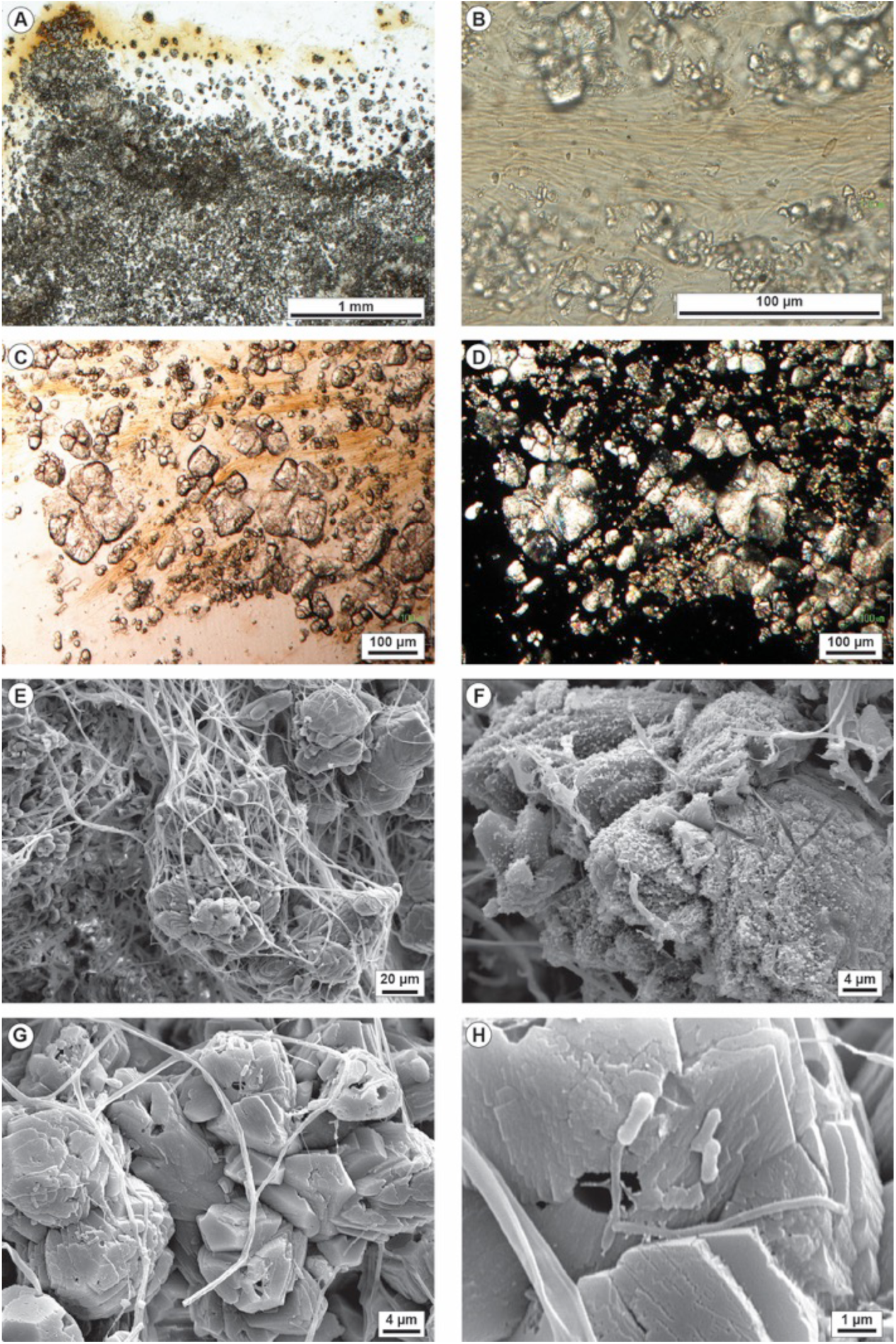
Petrographic and SEM images of the Gorello Waterfall (Saturnia) travertines from the rims and walls of the terraced slope system. A) Tetracycline dyed sample showing the framework of clotted peloidal micrite and microsparite spherulites overlain by the outer microbial mat. B) Close-up view in a tetracycline dyed sample showing microsparite spherulites alternating with microbial biofilm with dense concentration of filamentous microbes including *Spirulina* cyanobacteria. C) Microsparite to sparite spherulites floating within filamentous microbes. D) Crossed-polarizers image of microsparite/sparite spherulites showing the undulose extinction of the fan-shaped calcite crystals. E) SEM image showing dodecahedral calcite crystals forming aggregates and rosettes surrounded by filamentous microbes. F) SEM image of calcite crystals coated by mucilaginous organic matter, likely EPS, with tubular moulds related to the entombed filamentous microbes. G) Dodecahedral calcite crystals with tubular moulds related to the entrapped filamentous microbes and rod-shaped microbes. H) Close-up view of rod-shaped microbes with a dumbbell shape.

**Figure S10.**
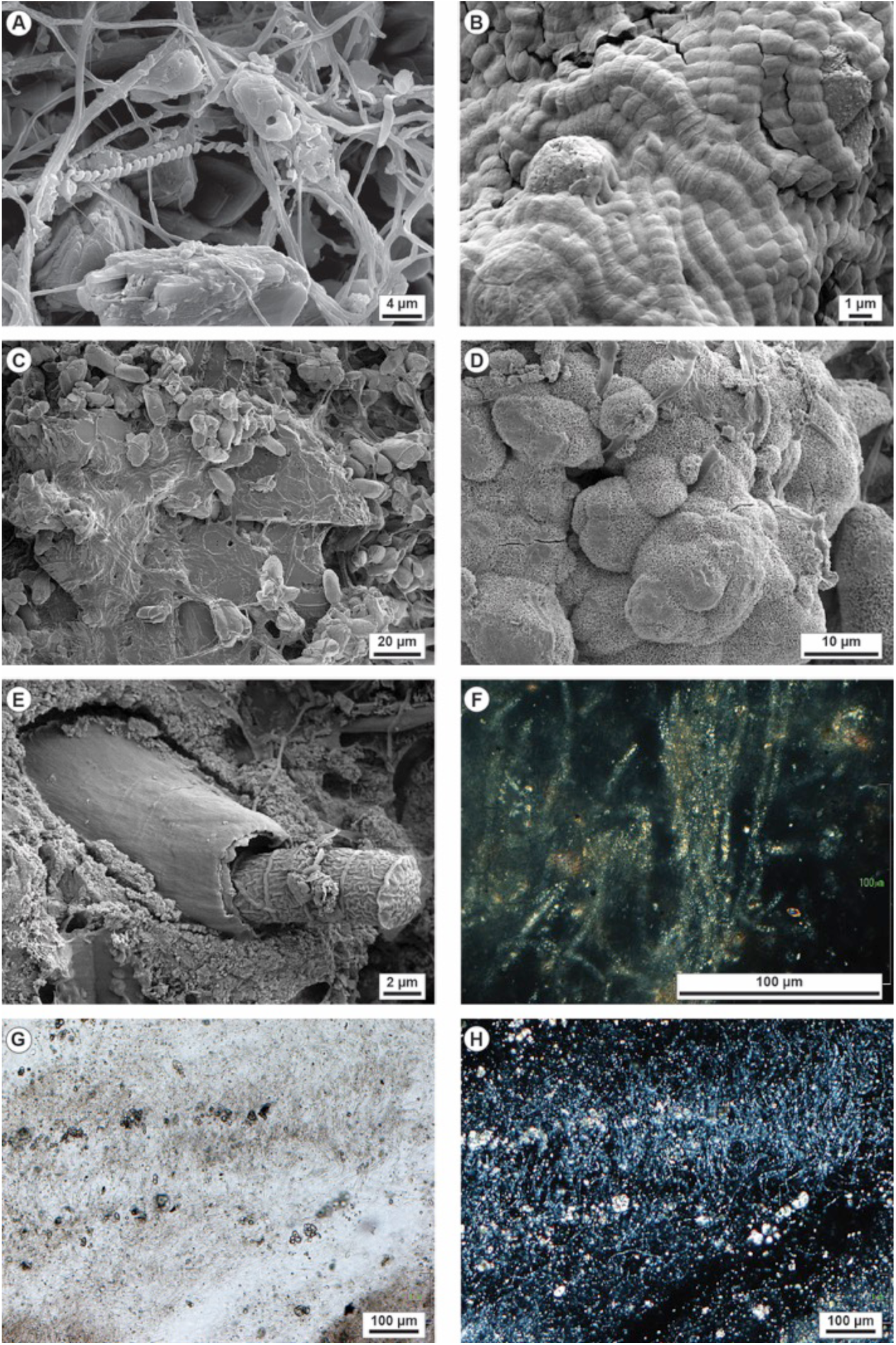
Petrographic and SEM images of the Gorello Waterfall (Saturnia) travertines from the rims and walls of the terraced slope system. A) SEM image showing calcite crystals surrounded by filamentous cyanobacteria including *Spirulina*. B) Segmented filamentous microbes probably belonging to *Phormidium*, *Oscillatoria* or *Fischerella* cyanobacteria. C) SEM image showing Ca phosphate crystals, probably apatite, coated by microsparite crystals, EPS and filamentous microbes with *Spirulina*. D) Spongy texture of phosphate coating the calcite crystals with a chemical composition of Ca and P through EDX analysis. E) Large segmented cyanobacteria with a thick sheath likely belonging to *Calothrix thermalis*. F) Crossed-polarizers image of large filamentous microbes probably belonging to the cyanobacteria *Calothrix* that show a birefringent internal filling material that could be an authigenic aluminium-silicate probably a clay mineral. G and H) Parallel and crossed polarizers photomicrographs of the microbial mat with erect filamentous microbes and sparse microsparite spherulites where the filamentous cyanobacteria are filled by aluminium-silicate material showing birefringence.

**Figure S11.**
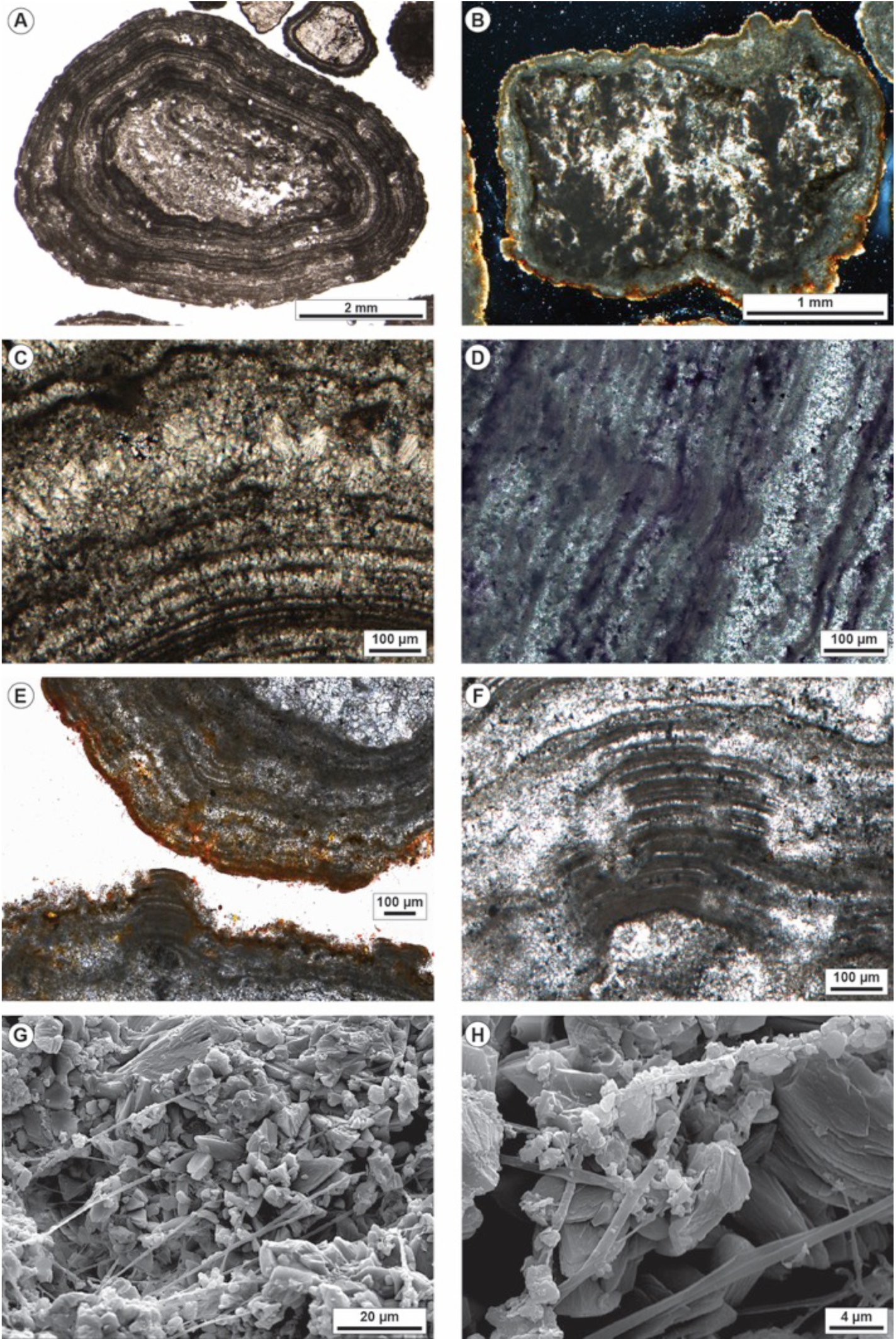
Petrographic and SEM images of the Gorello Waterfall (Saturnia) oncoids from the pools of the terraced slope system. A) Photomicrograph of oncoids with regular parallel undulated laminae made of micrite and microsparite and nuclei made of travertine intraclasts. B) Crossed polarizers image of an oncoid with the outer surface draped by tetracycline dyed microbial organic matter and nucleus made of travertine intraclast with clotted peloidal micrite dendrites surrounded by calcite spar. C) Crossed-polarizers image of oncoid laminae made of micrite alternating with palisades of bladed calcite crystals. D) Micritic and microsparitic laminae of oncoids containing organic matter as evidenced by toluidine dye. E) Outer portion of the cortex of two oncoids showing the microbial mat draping the outer surface as evidenced by calcein dye. The micritic laminae are locally eroded and truncated with organic matter infilling the microborings. F) Close-up view of truncated micritic laminae and microsparite replacing and filling the void left by removal of the micritic laminae. G) SEM image of calcite crystals forming the oncoid laminae with filamentous microbes extending perpendicular to the laminae surface. H) Filamentous microbes within the oncoid cortex encrusted by precipitated micrite.

**Figure S12.**
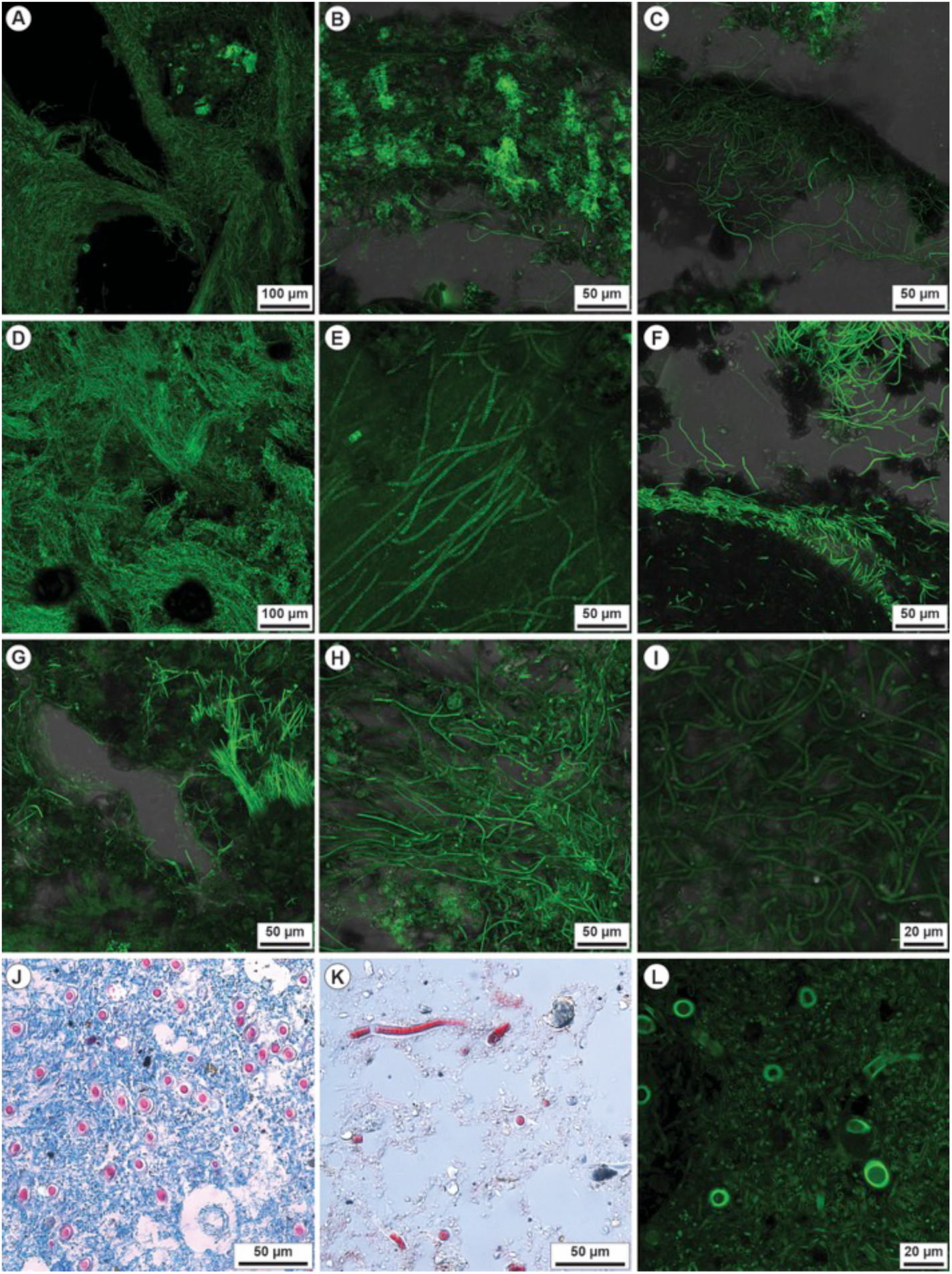
Confocal laser scanning microscope images of samples dyed with calcein from Bullicame (A-C), Bollore (D-F) and Gorello Waterfall (G-L). A) Sample from the Bullicame proximal channel showing the bundles of filamentous microbes forming the streamers, probably sulphide oxidizers. B-C) Sample from the Bullicame distal channel with abundant EPS and *Spirulina* filamentous cyanobacteria. D) Filamentous microbes, likely thermophilic sulphide oxidizing bacteria, from the calcified filamentous streamers sampled in the Bollore proximal channel. E) Bollore second vent channel with EPS embedding filamentous microbes including possible sulphide oxidizers and *Spirulina* cyanobacteria. F) Bollore distal channel with microbial mat dominated by *Spirulina* cyanobacteria. At the image top, calcite crystals are lined by green fluorescent organic matter. G) Gorello laminated boundstone with carbonate precipitates embedded in EPS and filamentous cyanobacteria including *Spirulina* and segmented forms. The porosity within the microbial mat lacks carbonate precipitates. H) Erect filamentous cyanobacteria alternating with prostrated filamentous microbes controlling the formation of lamination in the pool rim boundstone. I) Entangled filamentous cyanobacteria of the Gorello Waterfall microbial mat including also segmented *Phormidium* or *Oscillatoria* specimens. J) Paraffin prepared sample stained with alcian blue and cell centre red staining (Kernechtrot) showing the blue network of filamentous cyanobacteria associated with larger size cyanobacteria with a thick sheath, probably *Calothrix thermalis*. Alcian blue is a polysaccharide stain characterising EPS with abundant COO- groups. K) Paraffin prepared sample stained with Masson-Goldener solution showing the longitudinal section and the segmented appearance of the larger size cyanobacteria attributed to *Calothrix*. L) Confocal laser scanning microscope image showing the cross-section of the thinner filamentous cyanobacteria and the sparse larger size putative *Calothrix* with thick sheath.

**Table S1:**
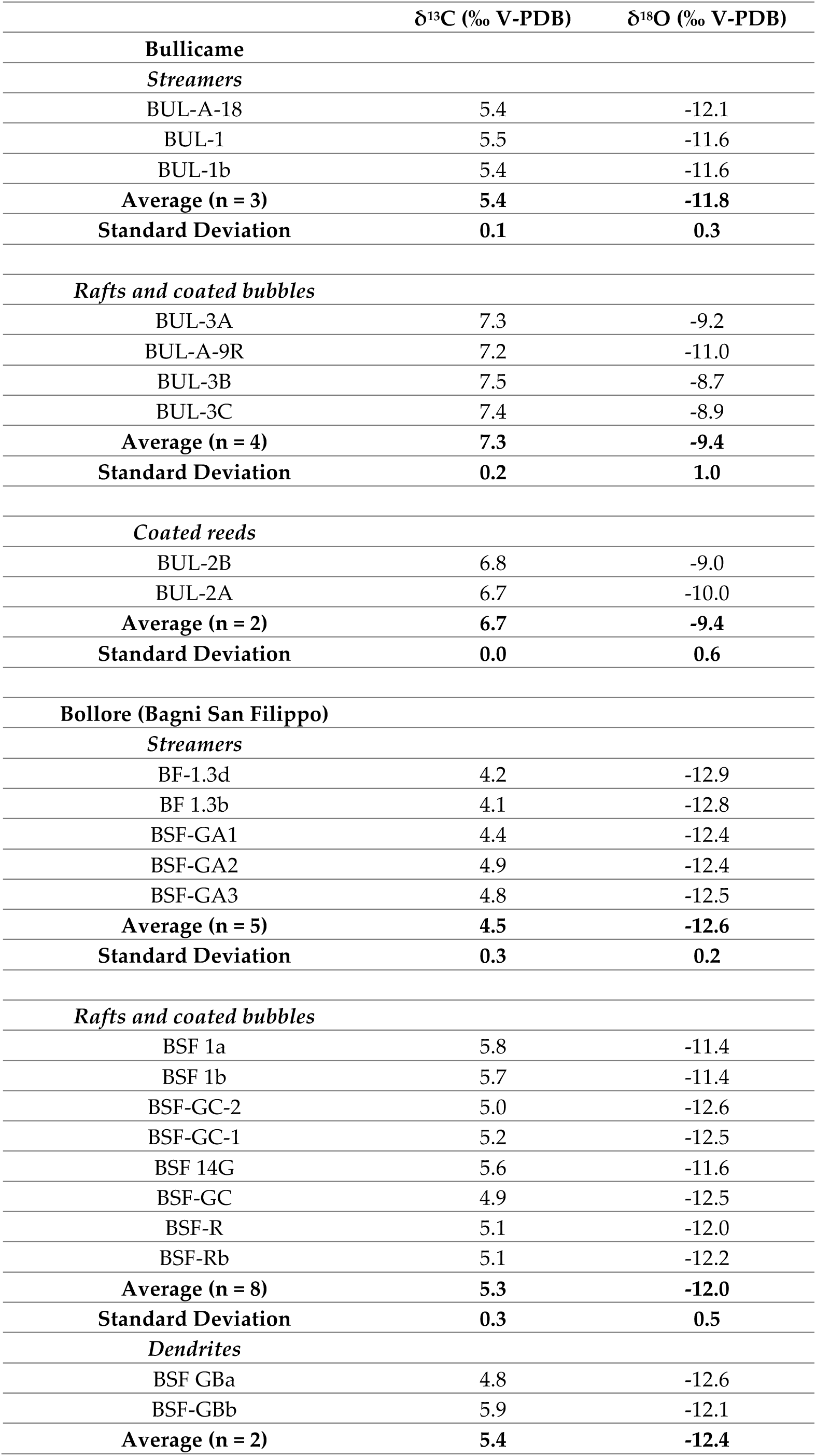
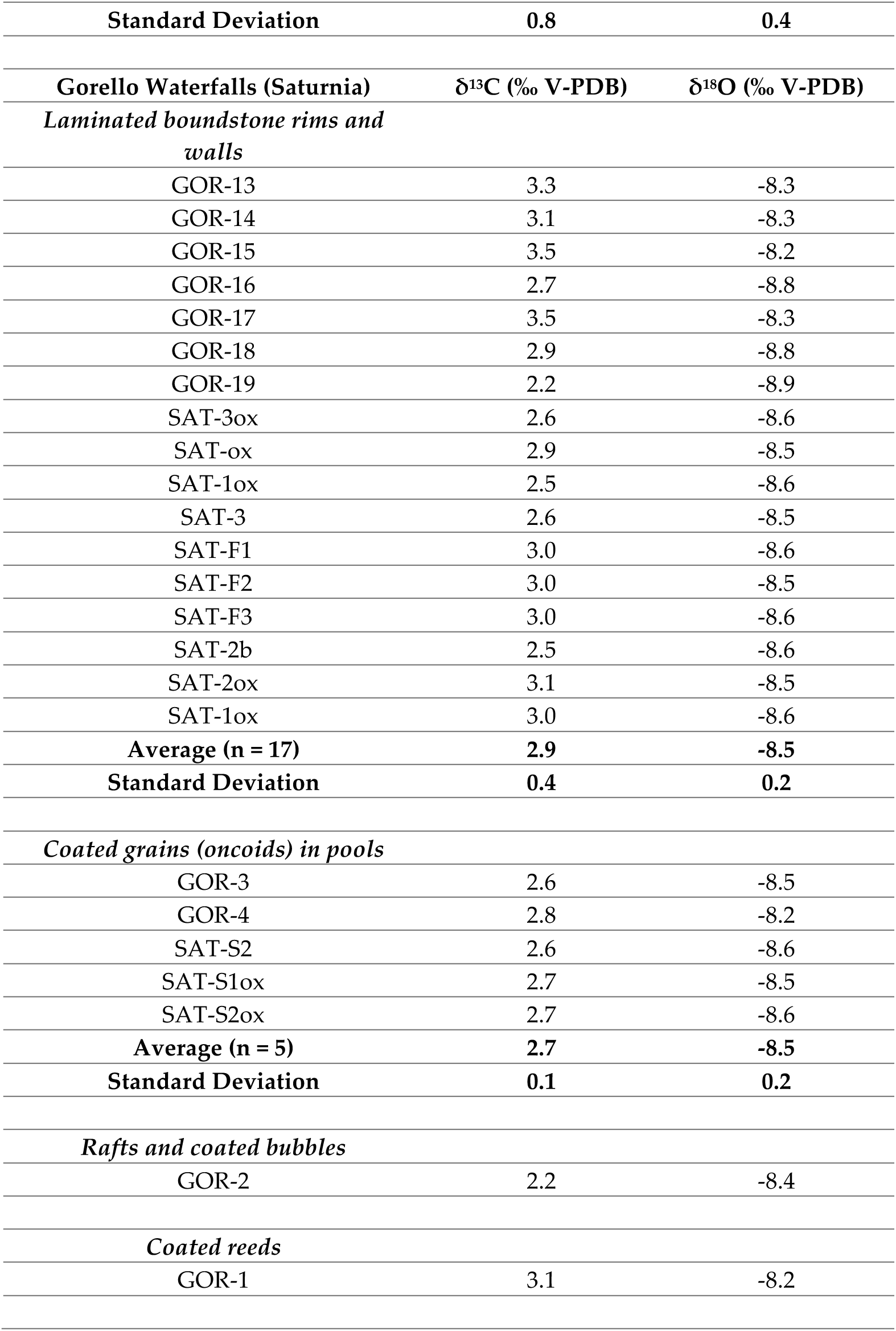
Results of stable carbon and oxygen isotope analyses from the three travertine study sites.

## Notes

### Competing Interest Statement

The authors have declared no competing interest.

## References

Addadi, L., Weiner, S., 1985. Interactions between acidic proteins and crystals: stereochemical requirements in biomineralization. Proceedings of the National Academy of Sciences 82, 12, 4110–4114.

Alain, K., Rolland, S., Crassous, P., Lesongeur, F., Zbinden, M., Le Gall, C., Godfroy, A., Page, A., Juniper, S.K., Cambon-Bonavita, M.A., Duchiron, F., 2003. *Desulfurobacterium crinifex* sp. nov., a novel thermophilic, pinkish-streamer forming, chemolithoautotrophic bacterium isolated from a Juan de Fuca Ridge hydrothermal vent and amendment of the genus *Desulfurobacterium*. Extremophiles 7, 361–370. DOI: 10.1007/s00792-003-0329-4

Allen, C.C., Albert, F.G., Chafetz, H.S., Combie, J., Graham, C.R., Kieft, T.L., Kivett, S.J., McKay, D.S., Steele, A., Taunton, A.E., Taylor, M.R., 2000. Microscopic physical biomarkers in carbonate hot springs: implications in the search for life on Mars. Icarus 147, 49–67. DOI: 10.1006/icar.2000.6435

Allwood, A.C., Walter, M.R., Kamber, B.S., Marshall, C.P., Burch, I.W., 2006. Stromatolite reef from the Early Archean era of Australia. Nature 441, 714–718. DOI: 10.1038/nature04764

Andreassen, J.P., Beck, R., Nergaard, M., 2012. Biomimetic type morphologies of calcium carbonate grown in absence of additives. Faraday Discussions 159, 247–261. DOI: 10.1039/c2fd20056b

Andres, M.S., Sumner, D.Y., Reid, R.P., Swart, P.K., 2006. Isotopic fingerprints of microbial respiration in aragonite from Bahamian stromatolites. Geology 34(11), 973–976. DOI: 10.1130/G22859A.1

Arp, G., Bissett, A., Brinkmann, N., Cousin, S., De Beer, D., Friedl, T., Mohr, K.I., Neu, T.R., Reimer, A., Shiraishi, F. and Stackebrandt, E., 2010. Tufa-forming biofilms of German karstwater streams: microorganisms, exopolymers, hydrochemistry and calcification. In: Pedley, H. M. & Rogerson, M. (eds) Tufas and Speleothems: Unravelling the Microbial and Physical Controls. Geological Society, London, Special Publications 336, 83–118. DOI: 10.1144/SP336.6

Arp, G., Thiel, V., Reimer, A., Michaelis, W., Reitner, J., 1999. Biofilm exopolymers control microbialites formation at thermal springs discharging into the alkaline Pyramid Lake, Nevada, USA. Sedimentary Geology 126, 159–176. DOI: 10.1016/S0037-0738(99)00038-X

Arp, G., Reimer, A., Reitner, J., 2001. Photosynthesis-induced biofilm calcification and calcium concentrations in Phanerozoic oceans. Science, 292(5522), 1701–1704. DOI: 10.1126/science.1057204

Arp, G., Reimer, A., Reitner, J., 2002. Calcification of cyanobacterial filaments: *Girvanella* and the origin of lower Paleozoic lime mud: comment. Geology 30, 579–580. DOI: 10.1130/0091-7613(2002)030<0579:COCFGA>2.0.CO;2

Arp, G., Reimer, A., Reitner, J., 2003. Microbialite formation in seawater of increased alkalinity, Satonda Crater Lake, Indonesia. Journal of Sedimentary Research 73, 105–127. DOI: 1527-1404/03/073-105/$03.00

Arp, G., Helms, G., Karlinska, K., Schumann, G., Reimer, A., Reitner, J., Trichet, J., 2012. Photosynthesis versus exopolymer degradation in the formation of microbialites on the atoll of Kiritimati, Republic of Kiribati, Central Pacific. Geomicrobiol. J. 29, 29–65. DOI: 10.1080/01490451.2010.521436

Baumgartner, L.K., Reid, R.P., Dupraz, C., Decho, A.W., Buckley, D.H., Spear, J.R., Przekop, K.M., Visscher, P.T., 2006. Sulfate reducing bacteria in microbial mats: changing paradigms, new discoveries. Sedimentary Geology 185(3-4), 131–145. DOI:10.1016/j.sedgeo.2005.12.008

Bazzichelli, G., Abdelhad, N., Florenzano, G., Tomaselli, L., 1978. Contributo alla conoscenza delle comunità fototrofiche delle Terme di Saturnia (Toscana). Ann. Botanica 37, 293–231.

Bissett, A., de Beer, D., Schoon, R., Shiraishi, F., Reimer, A., Arp, G., 2008. Microbial mediation of stromatolite formation in karst-water creeks. Limnology and Oceanography 53, 1159–1168. DOI: 10.4319/lo.2008.53.3.1159

Bigi G., Cosentino D., Parotto M., Sartori R., Scandone P., 1990. Structural Model of Italy 1: 500,000. La Ricerca Scientifica, Quaderni, C.N.R. 114 (3).

Bonny, S.M., Jones, B., 2008. Controls on the precipitation of barite (BaSO_4_) crystals in calcite travertine at Twitya Spring, a warm sulphur spring in Canada’s Northwest Territories. Sedimentary Geology 203, 36–53. DOI: 10.1016/j.sedgeo.2007.10.003

Bosak, T., Newman, D.K., 2005. Microbial kinetic controls on calcite morphology in supersaturated solutions. Journal Sedimentary Research 75, 190–199. DOI: 10.2110/jsr.2005.015

Braissant, O., Cailleau, G., Dupraz, C., Verrecchia, E.P., 2003. Bacterially induced mineralization of calcium carbonate in terrestrial environments: the role of exopolysaccharides and amino acids. Journal Sedimentary Research 73, 485–490. DOI: 10.1306/111302730485

Braissant, O., Decho, A. W., Dupraz, C., Glunk, C., Przekop, K.M., Visscher, P.T., 2007. Exopolymeric substances of sulfate-reducing bacteria: Interactions with calcium at alkaline pH and implication for formation of carbonate minerals. Geobiology 5, 401–411. DOI: 10.1111/j.1472-4669.2007.00117.x

Braissant, O., Decho, A.W., Przekop, K.M., Gallagher, K.L., Glunk, C., Dupraz, C., Visscher, P.T., 2009. Characteristics and turnover of exopolymeric substances in a hypersaline microbial mat. FEMS Microbiology Ecology 67(2), 293–307. DOI: :10.1111/j.1574-6941.2008.00614.x

Brasier, A.T., 2011. Searching for travertines, calcretes and speleothems in deep time: Processes, appearances, predictions and the impact of plants. Earth-Science Reviews 104, 213–239. DOI: doi:10.1016/j.earscirev.2010.10.007

Brogi, A., Capezzuoli, E., 2009. Travertine deposition and faulting: the fault-related travertine fissure ridge at Terme S. Giovanni, Rapolano Terme (Italy). Int. J. Earth Sci. 98, 931–947. DOI: 10.1007/s00531-007-0290-z

Brogi, A., Fabbrini, L., 2009. Extensional and strike-slip tectonics across the Monte Amiata–Monte Cetona transect (Northern Apennines, Italy) and seismotectonic implications. Tectonophysics 476, 195–209. DOI: 10.1016/j.tecto.2009.02.020

Brogi A., Liotta D., Ruggieri G., Capezzuoli E., Meccheri M., Dini A., 2016. An overview on the characteristics of geothermal carbonate reservoirs in southern Tuscany. Ital. J. Geosci. 135, 17–29. DOI: 10.3301/IJG.2014.41

Buick, R., Brauhart, C.W., Morant, P., Thornett, J.R., Maniw, J.G., Archibald, N.J., Doepel, M.G., Fletcher, I.R., Pickard, A.L., Smith, J.B., Barley, M.E., McNaughton, N.J., Groves, D.I., 2002. Geochronology and stratigraphic relationships of the Sulphur Springs Group and Strelley Granite: a temporally distinct igneous province in the Archaean Pilbara Craton, Australia. Precambrian Research 114, 87–120. DOI: 10.1016/S0301-9268(01)00221-2

Cady, S.L., Farmer, J.D., 1996. Fossilization processes in siliceous thermal springs: trends in preservation along thermal gradients, in: Evolution of hydrothermal ecosystems on Earth (and Mars?). Ciba Foundation Symposium 202 (Bock, G. R., Goode, G. A., Eds.), Wiley, Chichester, pp. 150–173.

Cady, S.L., Noffke, N., 2009. Geobiology: evidence for early life on Earth and the search for life on other planets. GSA Today 19/11, 4–10. doi:10.1130/GSATG62A.1

Cady, S.L.S., Skok, J.R., Gulick, V.G., Berger, J.A., Hinman, N.W., 2018. Siliceous hot spring deposits: Why they remain key astrobiologial targets, in From Habitability to Life on Mars, N. A. Cabrol and E. A. Grin, Eds. Edmond, OK: Elsevier Science, 179–210. DOI: 10.1016/B978-0-12-809935-3.00007-4

Campbell, K.A., Guido, D.M., Gautret, P., Foucher, F., Ramboz, C., Westall, F., 2015. Geyserite in hot-spring siliceous sinter: window on Earth’s hottest terrestrial (paleo) environment and its extreme life. Earth-Science Reviews 148, 44–64. DOI: 10.1016/j.earscirev.2015.05.009

Capezzuoli, E., Gandin, A., Pedley, M., 2014. Decoding tufa and travertine (fresh water carbonates) in the sedimentary record: the state of the art. Sedimentology 61, 1–21. DOI: 10.1111/sed.12075

Carminati, E., Doglioni, C., 2012. Alps vs. Apennines: the paradigm of a tectonically asymmetric Earth. Earth-Science Reviews 112, 67–96. DOI: 10.1016/j.earscirev.2012.02.004

Castanier, S., Le Métayer-Levrel, G., Perthuisot, J.P., 1999. Ca-carbonates precipitation and limestone genesis - the microbiogeologist point of view. Sedimentary Geology 126, 9–23. DOI: /10.1016/S0037-0738(99)00028-7

Chafetz, H.S., Buczynski, C., 1992. Bacterially induced lithification of microbial mats. Palaios 7, 277–293.

Chafetz, H.S., Folk, R.L., 1984. Travertines: depositional morphology and the bacterially constructed constituents. J. Sediment. Pet. 54, 289–316. DOI: 10.1306/212F8404-2B24-11D7-8648000102C1865D

Chafetz, H.S., Guidry, S.A. 1999. Bacterial shrubs, crystal shrubs, and ray-rystal shrubs: bacterial vs. abiotic precipitation. Sediment. Geol. 126, 57–74. DOI: 10.1016/S0037-0738(99)00032-9

Chekroun, K.B., Rodríguez-Navarro, C., González-Muñoz, M.T., Arias, J.M., Cultrone, G., Rodríguez-Gallego, M., 2004. Precipitation and growth morphology of calcium carbonate induced by *Myxococcus xanthus*: implications for recognition of bacterial carbonates. Journal Sedimentary Research 74, 868–876. DOI: 10.1306/050504740868

Cölfen, H., 2003. Precipitation of carbonates: recent progress in controlled production of complex shapes. Current Opinion Colloid Interface Sci. 8, 23–31. DOI: 10.1016/S1359-0294(03)00012-8

Cölfen, H., Antonietti, M., 2005. Mesocrystals: inorganic superstructures made by highly parallel crystallization and controlled alignment. Angewandte Chemie Inter. Ed. 44, 5576–5591. DOI: 10.1002/anie.200500496

Croci, A., Della Porta, G., Capezzuoli, E., 2016. Depositional architecture of a mixed travertine-terrigenous system in a fault-controlled continental extensional basin (Messinian, Southern Tuscany, Central Italy). Sedimentary Geology 332, 13–39. DOI: 10.1016/j.sedgeo.2015.11.007

Davis, K.J., Dove, P.M., De Yoreo, J.J., 2000. The role of Mg^2+^ as an impurity in calcite growth. Science 290, 1134–1137. DOI: 10.1126/science.290.5494.1134

Decho, A.W., 2010. Overview of biopolymer-induced mineralization: what goes on in biofilms? Ecological Engineering 36, 137–144. DOI: doi:10.1016/j.ecoleng.2009.01.003

Decho, A., Gutierrez, T., 2017. Microbial Extracellular Polymeric Substances (EPSs) in Ocean Systems. Frontiers in Microbiology 8, Article 922. doi: 10.3389/fmicb.2017.00922.

Défarge, C., Trichet, J., 1995. From biominerals to “organominerals”: the example of the modern lacustrine calcareous stromatolites from Polynesian atolls. Bull Inst Océanogr Monaco Spec Issue 14, 265–271.

Défarge, C., Trichet, J., Jaunet, A.-M., Robert, M., Tribble, J., Sansone, F.J., 1996. Texture of microbial sediments revealed by cryo-scanning electron microscopy. Journal of Sedimentary Research 66, 935–947. DOI: 10.1306/D4268446-2B26-11D7-8648000102C1865D

Défarge, C., Gautret, P., Reitner, J., Trichet, J., 2009. Defining organominerals: Comment on ‘Defining biominerals and organominerals: Direct and indirect indicators of life’ by Perry et al. (2007, Sedimentary Geology, 201, 157-179). Sedimentary Geology 213, 152–155. DOI: 10.1016/j.sedgeo.2008.04.002

Della Porta, G., 2015. Carbonate build-ups in lacustrine, hydrothermal and fluvial settings: comparing depositional geometry, fabric types and geochemical signature, in: Bosence, D.W.J., Gibbons, K.A., Le Heron, D.P., Morgan, W.A., Pritchard, T., Vining, B.A. (Eds.), Microbial Carbonates in Space and Time: Implications for Global Exploration and Production. Geol. Soc. London Spec. Publ. 418, pp. 17–68. DOI: 10.1144/SP418.4

Della Porta G., Capezzuoli, E., De Bernardo, A., 2017a. Facies character and depositional architecture of hydrothermal travertine slope aprons (Pleistocene, Acquasanta Terme, Central Italy), Marine Petroleum Geology 87, 171–187. DOI: 10.1016/j.earscirev.2003.09.001

Della Porta, G., Croci, A., Marini, M., Kele, S., 2017b. Depositional architecture, facies character and geochemical signature of the Tivoli travertines (Pleistocene, Acque Albule Basin, Central Italy). RIPS Riv. It. Paleont. Strat. 123, 487–540. DOI: 10.13130/2039-4942/9148

Des Marais, D.J., Walter, M.R., 2019. Terrestrial hot spring systems: introduction. Astrobiology 19(12), 1419–1432. DOI: 10.1089/ast.2018.1976

Di Benedetto, F., Montegrossi, G., Minissale, A., Pardi, L.A., Romanelli, M., Tassi, F., Delgado Huertas, A., Pampin, E.M., Vaselli, O., Borrini, D., 2011. Biotic and inorganic control on travertine deposition at Bullicame 3 spring (Viterbo, Italy): a multidisciplinary approach. Geochimica Cosmochimica Acta 75, 4441–4455. DOI: 10.1016/j.gca.2011.05.011

Djokic, T., Van Kranendonk, M.J., Campbell, K.A., Walter, M.R., Ward, C.R., 2017. Earliest signs of life on land preserved in ca. 3.5 Ga hot spring deposits. Nature communications 8, 1–9. DOI: 10.1038/ncomms15263

Doglioni, C., 1991. A proposal for the kinematic modelling of W-dipping subductions – possible applications to the Tyrrhenian-Apennines system. Terra Nova 3, 423–434. DOI: 10.1111/j.1365-3121.1991.tb00172.x

Duda, J.-P., Thiel, V., Bauersachs, T., Mißbach, H., Reinhardt, M., Schäfer, N., Van Kranendonk, M.J., Reitner, J., 2018. Ideas and perspectives: hydrothermally driven redistribution and sequestration of early Archaean biomass – the “hydrothermal pump hypothesis”. Biogeosciences 15, 1535–1548. DOI: 10.5194/bg-15-1535-2018

Dunckel, A.E., Cardenas, M.B., Sawyer, A.H., Bennett, P.C., 2009. High-resolution in-situ thermal imaging of microbial mats at El Tatio Geyser, Chile shows coupling between community color and temperature. Geophysical Research Letters 36, L23403. Doi: 10.1029/2009GL041366

Dupraz, C., Reid, R.P., Braissant, O., Decho, A.W., Norman, R.S., Visscher, P.T., 2009. Processes of carbonate precipitation in modern microbial mats. Earth-Science Reviews 96, 141–162. DOI: 10.1016/j.earscirev.2008.10.005

Dupraz, C., Visscher, P.T., 2005. Microbial lithification in marine stromatolites and hypersaline mats. Trends in Microbiology 13, 429–438. DOI: 10.1016/j.tim.2005.07.008

Dupraz, C., Visscher, P.T., Baumgartner, L.K., Reid, R.P., 2004. Microbe-mineral interactions: early carbonate precipitation in a hypersaline lake (Eleuthera Island, Bahamas). Sedimentology 51, 745–765. DOI: 10.1111/j.1365-3091.2004.00649.x

Eder, W., Huber, R., 2002. New isolates and physiological properties of the Aquificales and description of *Thermocrinis albus* sp. nov. Extremophiles 6, 309–318. DOI: 0.1007/s00792-001-0259-y

Edwards, D., Kenrick, P., Dolan, L., 2017. History and contemporary significance of the Rhynie cherts - our earliest preserved terrestrial ecosystem. Philosiphical Transaction Royal Society London B, Biol Sci 373, 20160489. DOI:10.1098/rstb.2016.0489.

Erthal, M.M., Capezzuoli, E., Mancini, A., Claes, H., Soete, J., Swennen, R., 2017. Shrub morpho-types as indicator for the water flow energy - Tivoli travertine case (Central Italy). Sedimentary Geology 347, 79–99. DOI: 10.1016/j.sedgeo.2016.11.008

Eymard, I., Alvarez, M.D.P., Bilmes, A., Vasconcelos, C., Ariztegui, D., 2020. Tracking organomineralization processes from living microbial mats to fossil microbialites. Minerals 10, 605. DOI:10.3390/min10070605

Faccenna, C., Soligo, M., Billi, A., De Filippis, L., Funiciello, R., Rossetti, C., Tuccimei, P., 2008. Late Pleistocene depositional cycles of the Lapis Tiburtinus travertine (Tivoli, central Italy): possible influence of climate and fault activity. Global and Planetary Change 63, 299–308. DOI: doi:10.1016/j.gloplacha.2008.06.006

Falini, G., Fermani, S., Gazzano, M., Ripamonti, A., 2000. Polymorphism and architectural crystal assembly of calcium carbonate in biologically inspired polymeric matrices. J. Chemic. Soc., Dalton Transactions 21, 3983–3987. DOI: 10.1039/B003334K

Farmer, J., 1998. Thermophiles, early biosphere evolution, and the origin of life on Earth: implications for the exobiological exploration of Mars. J. Geophys. Res. Planets 103(E12), 28457–28461. DOI: 10.1029/98JE01542

Farmer, J.D., 2000. Hydrothermal systems: doorways to early biosphere evolution. GSA Today 10, 1–9.

Fernàndez-Díaz, L., Putnis, A., Prieto, M. and Putnis, C.V., 1996. The role of magnesium in the crystallization of calcite and aragonite in a porous medium. Journal Sedimentary Research 66, 482–491. DOI: 10.1306/D4268388-2B26-11D7-8648000102C1865D

Flores, G.E., Liu, Y., Ferrera, I., Beveridge, T.J., Reysenbach, A.L., 2008. *Sulfurihydrogenibium kristjanssonii* sp. nov., a hydrogen-and sulfur-oxidizing thermophile isolated from a terrestrial Icelandic hot spring. Int. J. System. Evolution. Microbiol. 58, 1153–1158. DOI: 10.1099/ijs.0.65570-0

Folk, R.L., 1993. SEM imaging of bacteria and nannobacteria in carbonate sediments and rocks. Journal Sedimentary Research 63, 990–999. DOI: 10.1306/D4267C67-2B26-11D7-8648000102C1865D

Folk, R., 1994. Interaction between bacteria, nannobacteria, and mineral precipitation in hot springs of central Italy. Géograp. Phys. Quat. 48, 233–246. DOI: 10.7202/033005ar

Folk, R.L., Chafetz H.S., Tiezzi P.A., 1985. Bizarre forms of depositional and diagenetic calcite in hot-spring travertines, central Italy, in Schneidermann, N., Harris, P.M. (Eds.), Carbonate cements. SEPM Spec. Publ. 36, 349–369.

Fouke, B.W., 2011. Hot-spring Systems Geobiology: abiotic and biotic influences on travertine formation at Mammoth Hot Springs, Yellowstone National Park, USA. Sedimentology 58, 170–199. DOI: 10.1111/j.1365-3091.2010.01209.x

Fouke, B.W., Bonheyo, G.T., Sanzenbacher, B., Frias-Lopez J., 2003. Partitioning of bacterial communities between travertine depositional facies at Mammoth Hot Springs, Yellowstone National Park, USA. Can. J. Earth Sci. 40, 1531–1548. DOI: 10.1139/e03-067

Fouke, B.W., Farmer, J.D., Des Marais, D.J., Pratt, L., Sturchio, N.C., Burns, P.C., Discipulo, M.K., 2000. Depositional facies and aqueos-solid geochemistry of Travertine-depositing hot spring (Angel Terrace, Mammoth hot spring, Yellowstone National Park, U.S.A.). J. Sediment. Res. 70, 202–207. DOI: org/10.1306/2DC40929-0E47-11D7-8643000102C1865D

Franchi, F., Frisia, S., 2020. Crystallization pathways in the Great Artesian Basin (Australia) spring mound carbonates: Implications for life signatures on Earth and beyond. Sedimentology. DOI: 10.1111/sed.12711

Gandin, A., Capezzuoli, E., 2008. Travertine vs. calcareous tufa: distinctive petrologic features and stable isotope signatures. It. J. Quat. Sci. 21, 125–136.

Gandin, A., Capezzuoli, E., 2014. Travertine: distinctive depositional fabrics of carbonates from thermal spring systems. Sedimentology 61, 264–290. DOI: 10.1111/sed.12087

Giovannoni, S.J., Revsbech, N.P., Ward, D.M., Castenholz, R.W., 1987. Obligately phototrophic *Chloroflexus*: primary production in anaerobic hot spring microbial mats. Archiv. Microbiol. 147, 80–87. DOI: 10.1007/BF00492909

Glunk, C., Dupraz, C., Braissant, O., Gallagher, K.L., Verrecchia, E.P., Visscher, P.T., 2011. Microbially mediated carbonate precipitation in a hypersaline lake, Big Pond (Eleuthera, Bahamas). Sedimentology 58(3), 720–736. DOI: 10.1111/j.1365-3091.2010.01180.x

Golubic, S., Seong-Joo, L., Browne, K.M., 2000. Cyanobacteria: architects of sedimentary structures. In: R.E. Riding and S.M. Awramik (Editors), Microbial Sediments. Springer-Verlag, Berlin, Heidelberg, 57–67.

Gong, Y.U., Killian, C.E., Olson, I.C., Appathurai, N.P., Amasino, A.L., Martin, M.C., Holt, L.J., Wilt, F.H., Gilbert, P.U.P.A., 2012. Phase transitions in biogenic amorphous calcium carbonate. Proceedings of the National Academy of Sciences 109(16), 6088–6093. DOI: 10.1073/pnas.1118085109

Gong, J., Myers, K.D., Munoz-Saez, C., Homann, M., Rouillard, J., Wirth, R., Schreiber, A., van Zuilen, M.A., 2020. Formation and preservation of microbial palisade fabric in silica deposits from El Tatio, Chile. Astrobiology 20, 4, 500–524. Doi: 10.1089/ast.2019.2025

Gower, L.A., Tirrell, D.A.J., 1998. Calcium carbonate films and helices grown in solutions of poly (aspartate). Journal of Crystal Growth 191, 153–160. DOI: 10.1016/S0022-0248(98)00002-5

Guido, D.M., Campbell, K.A., 2011. Jurassic hot spring deposits of the Deseado Massif (Patagonia, Argentina): characteristics and controls on regional distribution. Journal of Volcanology and Geothermal Research, 203(1-2), 35–47. DOI: 10.1016/j.jvolgeores.2011.04.001

Guo, L., Andrews, J., Riding, R., Dennis, P., Dresser, Q., 1996. Possible microbial effects on stable carbon isotopes in hot-spring travertines. J. Sediment. Res. 66, 468–473. DOI: 10.1306/D4268379-2B26-11D7-8648000102C1865D

Guo, L., Riding, R., 1992. Aragonite laminae in hot water travertine crusts, Rapolano Terme, Italy. Sedimentology 39, 1067–1079. DOI: 10.1111/j.1365-3091.1992.tb01997.x

Guo, L., Riding, R., 1994. Origin and diagenesis of Quaternary travertine shrub fabrics, Rapolano Terme, central Italy. Sedimentology 41, 499–520. DOI: 10.1111/j.1365-3091.1994.tb02008.x

Guo, L., Riding, R., 1998. Hot-spring travertine facies and sequences, Late Pleistocene, Rapolano Terme, Italy. Sedimentology 45, 163–180. DOI: 10.1046/j.1365-3091.1998.00141.x

Guo, L., Riding, R., 1999. Rapid facies changes in Holocene fissure ridge hot spring travertines, Rapolano Terme, Italy. Sedimentology 46, 1145–1158. DOI: 10.1046/j.1365-3091.1999.00269.x

Hoffmann, F., Janussen, D., Dröse, W., Arp, G., Reitner, J., 2003. Histological investigation of organisms with hard skeletons: a case study of siliceous sponges. Biotechnic & Histochemistry 78, 191–199. DOI: 10.1080/10520290310001613042

Ionescu, D., Spitzer, S., Reimer, A., Schneider, D., Daniel, R., Reitner, J., de Beer, D., Arp, G., 2014. Calcium dynamics in microbialite-forming exopolymer-rich mats on the atoll of Kiritimati, Republic of Kiribati, Central Pacific. Geobiology 13(2), 170–180. DOI: 10.1111/gbi.12120.

Jones, B., 2009. Phosphatic precipitates associated with actinomycetes in speleothems from Grand Cayman, British West Indies. Sedimentary Geology 219, 302–317. DOI: 10.1016/j.sedgeo.2009.05.020

Jones, B., 2017a. Review of calcium carbonate polymorph precipitation in spring systems. Sedimentary Geology 353, 64–75. DOI: 10.1016/j.sedgeo.2017.03.006

Jones, B., 2017b. Review of aragonite and calcite crystal morphogenesis in thermal spring systems. Sediment. Geol. 354, 9–23. DOI: 10.1016/j.sedgeo.2017.03.012

Jones, B., Peng, X., 2012. Amorphous calcium carbonate associated with biofilms in hot spring deposits. Sedimentary Geology 269, 58–68. DOI: 10.1016/j.sedgeo.2012.05.019

Jones, B., Peng, X., 2014a. Signatures of biologically influenced CaCO_3_ and Mg–Fe silicate precipitation in hot springs: case study from the Ruidian geothermal area, western Yunnan Province, China. Sedimentology 61, 56–89. DOI: 10.1111/sed.12043

Jones, B., Peng, X., 2014b. Hot spring deposits on a cliff face: a case study from Jifei, Yunnan Province, China. Sedimentary Geology 302, 1–28. DOI: 10.1016/j.sedgeo.2013.12.009

Jones, B., Peng, X., 2015. Laminae development in opal-A precipitates associated with seasonal growth of the form-genus *Calothrix* (Cyanobacteria), Rehai geothermal area, Tengchong, Yunnan Province, China. Sedimentary Geology 319, 52–68. DOI: 10.1016/j.sedgeo.2015.01.004

Jones, B., Peng, X., 2016. Mineralogical, crystallographic, and isotopic constraints on the precipitation of aragonite and calcite at Shiqiang and other hot springs in Yunnan Province, China. Sedimentary Geology 345, 103–125. DOI: 10.1016/j.sedgeo.2016.09.007

Jones, B., Renaut, R.W., 1995. Noncrystallographic calcite dendrites from hot-spring deposits at Lake Bogoria, Kenya. Journal Sedimentary Research 65, 154–169. DOI: 10.1306/D4268059-2B26-11D7-8648000102C1865D

Jones, B., Renaut, R.W., 2010. Calcareous spring deposits in continental settings. In: Alonso-Zarza, M. & Tanner, L. H. (eds) Carbonates in Continental Settings: Facies, Environments and Processes. Developments in Sedimentology 61, Elsevier, Amsterdam, 177–224.

Jones, B, Renaut, R.W., 2011. Hot springs and Geysers. In: Reitner, J. & Thiel, V., Encyclopedia of Geobiology, Springer, Berlin, Heidelberg, 447–451.

Kamennaya, N.A., Ajo-Franklin, C.M., Northen, T., Jansson, C., 2012. Cyanobacteria as biocatalysts for carbonate mineralization. Minerals 2(4), 338–364. DOI: 10.3390/min2040338

Kato, T., Sugawara, A., Hosoda, N., 2002. Calcium carbonate–organic hybrid materials. Adv. Mater. 14, 869–877. DOI: 10.1002/1521-4095(20020618)14:12<869::AID-ADMA869>3.0.CO;2-E

Kele, S., Breitenbach, S.F., Capezzuoli, E., Meckler, A.N., Ziegler, M., Millan, I.M., Kluge, T., Deák, J., Hanselmann, K., John, C.M., Yan, H., 2015. Temperature dependence of oxygen- and clumped isotope fractionation in carbonates: a study of travertines and tufas in the 6-95°C temperature range. Geochimica Cosmochimica Acta 168, 172–192. DOI: 10.1016/j.gca.2015.06.032

Keller, H., Plank, J., 2013. Mineralisation of CaCO_3_ in the presence of polycarboxylate comb polymers. Cement Concrete Research 54, 1–11. DOI: 10.1016/j.cemconres.2013.06.017

Kirkham, A., Tucker, M.E., 2018. Thrombolites, spherulites and fibrous crusts (Holkerian, Purbeckian, Aptian): Context, fabrics and origins. Sedimentary Geology 374, 69–84. DOI: 10.1016/j.sedgeo.2018.07.002

Konhauser, K., 2007. Introduction to Geomicrobiology. Blackwell Publishing, Singapore, 425 pp.

Konhauser, K.O., Phoenix, V.R., Bottrell, S.H., Adams, D.G., Head, I.M., 2001. Microbial-silica interactions in Icelandic hot spring sinter: possible analogues for some Precambrian siliceous stromatolites. Sedimentology 48, 415–433. DOI: 10.1046/j.1365-3091.2001.00372.x

Konopacka-Łyskawa, D., Kościelska, B., Karczewski, J., 2017. Controlling the size and morphology of precipitated calcite particles by the selection of solvent composition. Journal of Crystal Growth 478, 102–110. DOI: 10.1016/j.jcrysgro.2017.08.033

Kosanović, C., Fermani, S., Falini, G., Kralj, D., 2017. Crystallization of Calcium Carbonate in Alginate and Xanthan Hydrogels. Crystals 7, 355–370. DOI: 10.3390/cryst7120355

Knorre, H.V., Krumbein, W.E., 2000. Bacterial Calcification. In: R.E. Riding and S.M. Awramik (Editors), Microbial Sediments. Springer-Verlag, Berlin Heidelberg, 23–31.

Kremer, B., Kaźmierczak, J., Kempe, S., 2019. Authigenic replacement of cyanobacterially precipitated calcium carbonate by aluminium-silicates in giant microbialites of Lake Van (Turkey). Sedimentology 66, 285–304. DOI: 10.1111/sed.12529

Kubo, K., Knittel, K., Amann, R., Fukui, M., Matsuura, K., 2011. Sulfur-metabolizing bacterial populations in microbial mats of the Nakabusa hot spring, Japan. System. Appl. Microbiol. 34, 293–302. DOI: 10.1016/j.syapm.2010.12.002

Lakshtanov, L.Z., Stipp, S.L.S., 2010. Interaction between dissolved silica and calcium carbonate: Spontaneous precipitation of calcium carbonate in the presence of dissolved silica. Geochimica Cosmochimica Acta 74, 2655–2664. DOI: 10.1016/j.gca.2010.02.009

Lalonde, S.V., Konhauser, K.O., Reysenbach, A.L., Ferris, F.G., 2005. The experimental silicification of Aquificales and their role in hot spring sinter formation. Geobiology 3, 41–52. DOI: 10.1111/j.1472-4669.2005.00042.x

Lowenstam, H.A., Weiner, S., 1989. On Biomineralization. Oxford University Press, New York Oxford, 324 pp.

Ma, Y., Feng, Q., 2015. A crucial process: organic matrix and magnesium ion control of amorphous calcium carbonate crystallization on β-chitin film. CrystEngComm 17(1), 32–39. DOI: 10.1039/C4CE01616E.

Madigan, M.T., 2003. Anoxygenic phototrophic bacteria from extreme environments. Photosynthesis Res., 76, 157–171. DOI: 10.1023/A:1024998212684

Malinverno, A., Ryan, W.B.F., 1986. Extension in the Tyrrhenian Sea and shortening in the Apennines as result of arc migration driven by sinking of the lithosphere. Tectonics 5, 227–254. DOI: 10.1029/TC005i002p00227

Meldrum, F.C., Cölfen, H., 2008. Controlling mineral morphologies and structures in biological and synthetic systems. Chemical Reviews 108, 4332–4432. DOI: 10.1021/cr8002856

Meldrum, F.C., Hyde, S.T., 2001. Morphological influence of magnesium and organic additives on the precipitation of calcite. Journal of Crystal Growth 231, 544–558. DOI: 10.1016/S0022-0248(01)01519-6

Melezhik, V.A., Fallick, A.E., 2001. Palaeoproterozoic travertines of volcanic affiliation from a ^13^C-rich rift lake environment. Chemical Geology 173, 293–312. DOI: 10.1016/S0009-2541(00)00281-3

Mercedes-Martín, R., Rogerson, M.R., Brasier, A.T., Vonhof, H.B., Prior, T.J., Fellows, S.M., Reijmer, J.J.G., Billing, I., Pedley, H.M., 2016. Growing spherulitic calcite grains in saline, hyperalkaline lakes: experimental evaluation of the effects of Mg-clays and organic acids. Sedimentary Geology 335, 93–102. DOI: 10.1016/j.sedgeo.2016.02.008

Merz, M.U.E., 1992. The biology of carbonate precipitation by cyanobacteria. Facies 26, 81–102. DOI: 10.1007/BF02539795

Merz-Preiß, M., 2000. Calcification in cyanobacteria. In: R.E. Riding and S.M. Awramik (Editors), Microbial Sediments. Springer-Verlag, Berlin, Heidelberg, 50–56.

Merz-Preiß, M., Riding, R., 1999. Cyanobacterial tufa calcification in two freshwater streams: ambient environment, chemical thresholds and biological processes. Sedimentary Geology 126, 103–124. DOI: 10.1016/S0037-0738(99)00035-4

Minissale, A., 2004. Origin, transport and discharge of CO_2_ in central Italy. Earth-Science Reviews 66, 89–141. DOI:10.1016/j.earscirev.2003.09.001

Minissale, A., Kerrick, D.M., Magro, G., Murrell, M.T., Paladini, M., Rihs, S., Sturchio, N.C., Tassi, F., Vaselli, O., 2002a. Geochemistry of Quaternary travertines in the region north of Rome (Italy): Structural, hydrologic and paleoclimatic implications. Earth Planet. Sci. Lett. 203, 709–728. DOI: 10.1016/S0012-821X(02)00875-0

Minissale, A., Vaselli, O., Tassi, F., Magro, G., Grechi, G.P., 2002b. Fluid mixing in carbonate aquifers near Rapolano (central Italy): chemical and isotopic constraints. Appl. Geochem. 17, 1329–1342. DOI: 10.1016/S0883-2927(02)00023-9

Mori, K., Suzuki, K.I., 2008. *Thiofaba tepidiphila* gen. nov., sp. nov., a novel obligately chemolithoautotrophic, sulfur-oxidizing bacterium of the Gammaproteobacteria isolated from a hot spring. Int. J. System. Evolution. Microbiol. 58, 1885–1891. DOI: 10.1099/ijs.0.65754-0

Nakagawa, S., Shtaih, Z., Banta, A., Beveridge, T.J., Sako, Y., Reysenbach, A.L., 2005. *Sulfurihydrogenibium yellowstonense* sp. nov., an extremely thermophilic, facultatively heterotrophic, sulfur-oxidizing bacterium from Yellowstone National Park, and emended descriptions of the genus *Sulfurihydrogenibium, Sulfurihydrogenibium subterraneum* and *Sulfurihydrogenibium azorense*. Int. J. System. Evolution. Microbiol. 55, 2263–2268. DOI: 10.1099/ijs.0.63708-0

Neuweiler, F., Gautret, P., Thiel, V., Lange, R., Michaelis, W., Reitner, J., 1999. Petrology of Lower Cretaceous carbonate mud mounds (Albian, N. Spain): insights into organomineralic deposits of the geological record. Sedimentology 46, 837–859. DOI: /10.1046/j.1365-3091.1999.00255.x

Norris, T.B., Castenholz, R.W., 2006. Endolithic photosynthetic communities within ancient and recent travertine deposits in Yellowstone National Park. FEMS Microbiol. Ecol. 57, 470–483. DOI: 10.1111/j.1574-6941.2006.00134.x

Norris, T.B., McDermott, T.R., Castenholz, R.W., 2002. The long-term effects of UV exclusion on the microbial composition and photosynthetic competence of bacteria in hot-spring microbial mats. FEMS Microbiol. Ecol. 39, 193–209. DOI: 10.1111/j.1574-6941.2002.tb00922.x

Oaki, Y., Imai, H., 2003. Experimental demonstration for the morphological evolution of crystals grown in gel media. Crystal Growth Design 3, 711–716. DOI: 10.1021/cg034053e

Obst, M., Dynes, J.J., Lawrence, J.R., Swerhone, G.D.W., Benzerara, K., Karunakaran, C., Kaznatcheev, K., Tyliszczak, T., Hitchcock, A.P., 2009. Precipitation of amorphous CaCO_3_ (aragonite-like) by cyanobacteria: a STXM study of the influence of EPS on the nucleation process. Geochimica Cosmichimica Acta 73, 4180–4198. DOI: 10.1016/j.gca.2009.04.013

Okumura, T., Takashima, C., Kano, A., 2013. Textures and processes of laminated travertines formed by unicellular cyanobacteria in Myoken hot spring, southwestern Japan. Island Arc 22, 410–426. DOI: 10.1111/iar.12034

Pedley, H.M., 1990. Classification and environmental models of cool freshwater tufas. Sediment. Geol. 68, 143–154. DOI: 10.1016/0037-0738(90)90124-C

Pedley, M., 2014. The morphology and function of thrombolytic calcite precipitating biofilms: a universal model derived from freshwater mesocosm experiments. Sedimentology 61, 22–40. DOI: 10.1111/sed.12042

Pedley, H.M., Rogerson, M. Middleton, R., 2009. Freshwater calcite precipitates from in vitro mesocosm flume experiments: a case for biomediation of tufas. Sedimentology 56, 511–527. DOI: 10.1111/j.1365-3091.2008.00983.x

Peng, X., Jones, B., 2013. Patterns of biomediated CaCO_3_ crystal bushes in hot spring deposits. Sedimentary Geology 294, 105–117. DOI: 10.1016/j.sedgeo.2013.05.009

Pentecost, A., 1994. Formation of laminate travertines at Bagno Vignone, Italy. Geomicrobiol. J. 12, 239–251. DOI: 10.1080/01490459409377992

Pentecost, A., 1995a. Geochemistry of carbon dioxide in six travertine-depositing waters of Italy. J. Hydrol. 167, 263–278. DOI: 10.1016/0022-1694(94)02596-4

Pentecost, A., 1995b. The Quaternary travertine deposits of Europe and Asia Minor. Quat. Sci. Rev. 14, 1005–1028. DOI: 10.1016/0277-3791(95)00101-8

Pentecost, A., 2003. Cyanobacteria associated with hot spring travertines. Can. J. Earth Sci. 40, 1447–1457. DOI: 10.1139/e03-075

Pentecost, A., 2005. Travertine. Springer, Berlin Heidelberg, 445 pp.

Pentecost, A., Bayari, S., Yesertener, C., 1997. Phototrophic microorganisms of the Pamukkale travertine Turkey: their distribution and influence on travertine deposition. Geomicrobiol. J. 14, 269–283. DOI: 10.1080/01490459709378052

Pentecost, A., Coletta, P., 2007. The role of photosynthesis and CO_2_ evasion in travertine formation: a quantitative investigation at an important travertine-depositing hot spring, Le Zitelle, Lazio, Italy. J. Geol. Soc. 164, 843–853. DOI: 10.1144/0016-76492006-037

Pentecost, A., Riding, R., 1986. Calcification of cyanobacteria. In: B.S.C. Leadbeater and R. Riding (Editors), Biomineralization in Lower Plants and Animals. Systematic Association, Special Volume 30, 73–90.

Pentecost, A., Tortora, P., 1989. Bagni di Tivoli, Lazio: a modern travertine-depositing site and its associated microorganisms. Boll. Soc. Geol. It. 108, 315–324.

Piscopo, V., Barbieri, M., Monetti, V., Pagano, G., Pistoni, S., Ruggi, E., Stanzione, D., 2006. Hydrogeology of thermal waters in Viterbo area, central Italy. Hydrogeol. J. 14, 1508–1521. DOI: 10.1007/s10040-006-0090-8

Plée, K., Pacton, M., Ariztegui, D., 2010. Discriminating the role of photosynthetic and heterotrophic microbes triggering low-Mg calcite precipitation in freshwater biofilms (Lake Geneva, Switzerland). Geomicrobiology Journal, 27, 391–399. DOI: 10.1080/01490450903451526

Rainey, D. K., Jones, B., 2009. Abiotic v. biotic controls on the development of the Fairmont Hot Springs carbonate deposit, British Columbia, Canada. Sedimentology 56, 1832–1857. DOI: 10.1111/j.1365-3091.2009.01059.x

Reinhardt, M., Goetz, W., Duda, J.P., Heim, C., Reitner, J., Thiel, V., 2019. Organic signatures in Pleistocene cherts from Lake Magadi (Kenya) - implications for early Earth hydrothermal deposits. Biogeosciences 16(12), 2443–2465. DOI: 10.5194/bg-16-2443-2019

Reitner, J., 1993. Modern cryptic microbialite/metazoan facies from Lizard Island (Great Barrier Reef, Australia): Formation and Concept. Facies 29, 3–40. DOI: 10.1007/BF02536915

Reitner, J., 2004. Organomineralization: a clue to the understanding of meteorite related “bacteria-shaped” carbonate particles. In: Seckbach, J. (Ed.), Origins. Genesis, Evolution and Diversity of Life. Kluwer Academic Publishers, Dordrecht, The Netherlands, 195–212.

Reitner, J., Gautret, P., Marin, F., Neuweiler, F., 1995a. Automicrites in a modern marine microbialite. Formation model via organic matrices (Lizard Island, Great Barrier Reef, Australia). Bull Inst Océanogr Monaco Spec Issue 14, 237–263.

Reitner, J., Neuweiler, F. and Gautret, P., 1995b. Modern and fossil automicrites: implications for mud mound genesis. In: J. Reitner and F. Neuweiler (Coords), A polygenetic spectrum of fine-grained carbonate buildups. Facies 32, 4–17.

Reitner, J., Thiel, V., Zankl, H., Michaelis, W., Wörheide, G., Gautret, P., 2000. Organic and biogeochemical patterns in cryptic microbialites. In: R.E. Riding and S.M. Awramik (Editors), Microbial Sediments. Springer-Verlag, 149–160.

Reitner, J., Paul, J., Arp, G., Hause-Reitner, D., 1996. Lake Thetis domal microbialites – a complex framework of calcified biofilms and organomicrites (Cervantes, Western Australia). In: Reitner, J., Neuweiler, F. & Gunkel, F. (eds) Reef Evolution. Research Reports. Göttinger Arbeiten zur Geologie und Paläontologie, Sb2, 85–89.

Reitner, J., Arp, G., Thiel, V., Gautret, P., Galling, U., Michaelis, W., 1997. Organic matter in Great Salt Lake ooids (Utah, USA) – first approach to a formation of organic matrices. Facies 36, 210–219.

Reitner, J., Hoffmann, F., Dröse, W., 2004. Workshop Geohistologie.- In: Reitner, J., Reich, M., Schmidt, G. (eds.), Geobiologie 2, 74. Jahrestagung der Paläontologischen Gesellschaft in Göttingen 02. Bis 08. Oktober 2004, Exkursionen und Workshops; 211–226. Universitätsdrucke, Göttingen

Reitner, J., Wörheide, G., Lange, R., Schumann-Kindel, G., 2001. Coralline Demosponges - A geobiological portrait. In, Mori, K., Ezaki, Y. & Sorauf, J., eds., Proceedings of the 8th International Symposium on Fossil Cnidaria and Porifera, September 1999, Sendai. Bulletin of the Tohoku University Museum, 1: 219–235.

Reysenbach, A.L., Cady, S.L., 2001. Microbiology of ancient and modern hydrothermal systems. Trends Microbiol. 9, 79–86. DOI: 10.1016/S0966-842X(00)01921-1

Reysenbach A.L., Ehringer M., Hershberger, K., 2000. Microbial diversity at 83°C in Calcite Springs, Yellowstone National Park: another environment where the *Aquificales* and “Korarchaeota” coexist. Extremophiles 4, 61–67. DOI: 10.1007/s007920050008

Reysenbach, A.L., Seitzinger, S., Kirshtein, J., McLaughlin, E., 1999. Molecular constraints on a high-temperature evolution of early life. Biologic. Bull. 196, 367–372. DOI: 10.2307/1542972

Reysenbach, A.L., Shock, E., 2002. Merging genomes with geochemistry in hydrothermal ecosystems. Science 296, 1077–1082. DOI: 10.1126/science.1072483

Rice, C.M., Trewin, N.H., Anderson, L.I., 2002. Geological setting of the Early Devonian Rhynie Cherts, Aberdeenshire, Scotland: an early terrestrial hot spring system. J Geol Soc London 159, 203–214. DOI: 10.1144/0016-764900-181

Riding, R., 2000. Microbial carbonates: the geological record of calcified bacterial-algal mats and biofilms. Sedimentology 47, 179–214. DOI: 0.1046/j.1365-3091.2000.00003.x

Riding, R., 2008. Abiogenic, microbial and hybrid authigenic carbonate crusts: Components of Precambrian stromatolites. Geologia Croatica 61, 73–103.

Robbins, L.L., Blackwelder, P.L., 1992. Biochemical and ultrastructural evidence for the origin of whitings: A biologically induced calcium carbonate precipitation mechanism. Geology 20, 464–468. DOI: 10.1130/0091-7613(1992)020<0464:BAUEFT>2.3.CO;2

Rodriguez-Navarro, C., Kudłacz, K., Cizer, Ö., Ruiz-Agudo, E., 2015. Formation of amorphous calcium carbonate and its transformation into mesostructured calcite. CrystEngComm 17, 58–72. DOI: 10.1039/c4ce01562b

Roeselers, G., Norris, T.B., Castenholz, R.W., Rysgaard, S., Glud, R.N., Kühl, M., Muyzer, G., 2007. Diversity of phototrophic bacteria in microbial mats from Arctic hot springs (Greenland). Environ. Microbiol. 9, 26–38. DOI: 10.1111/j.1462-2920.2006.01103.x

Romeis, B., 1989. Mikroskopische Technik (ed. by P. Böck). 17. Aufl., Urban & Schwarzenberg, München.

Ronchi, P., Cruciani, F., 2015. Continental carbonates as a hydrocarbon reservoir, an analog case study from the travertine of Saturnia, Italy. AAPG Bulletin 99, 711–734. DOI: 0.1306/10021414026

Rothschild, L.J., Mancinelli, R.L., 2001. Life in extreme environments. Nature 409, 1092–1101. DOI: 10.1038/35059215

Roy, S., Debnath, M., Ray, S., 2014. Cyanobacterial flora of the geothermal spring at Panifala, West Bengal, India. Phykos 44, 1–8.

Ruff, S.W., Farmer, J.D., 2016. Silica deposits on Mars with features resembling hot spring biosignatures at El Tatio in Chile. Nature communications 7(1), 1–10. DOI: 10.1038/ncomms13554

Salminen, P.E., Brasier, A.T., Karhu, J.A., Melezhik, V.A., 2014. Travertine precipitation in the Paleoproterozoic Kuetsjärvi Sedimentary Formation, Pechenga Greenstone Belt, NE Fennoscandian Shield. Precambrian Research 255, 181–201. DOI: 0.1016/j.precamres.2014.09.023

Sanchez-Garcia, L., Fernandez-Martinez, M.A., García-Villadangos, M., Blanco, Y., Cady, S.L., Hinman, N., Bowden, M.E., Pointing, S.B., Lee, K.C., Warren-Rhodes, K., Lacap-Bugler, D., 2019. Microbial biomarker transition in high-altitude sinter mounds from El Tatio (Chile) through different stages of hydrothermal activity. Frontiers in Microbiology 9, 3350. DOI: 10.3389/fmicb.2018.03350

Sánchez-Navas, A., Martín-Algarra, A., Rivadeneyra, M.A., Melchor, S., Martín-Ramos, J.D., 2009. Crystal-growth behavior in Ca-Mg carbonate bacterial spherulites. Crystal Growth Design 9, 2690–2699. DOI: 10.1021/cg801320p

Sand, K.K., Rodriguez-Blanco, J.D., Makovicky, E., Benning, L.G., Stipp, S.L.S., 2011. Crystallization of CaCO_3_ in water-alcohol mixtures: spherulitic growth, polymorph stabilization, and morphology change. Crystal Growth Design 12, 842–853. DOI: 10.1021/cg2012342

Scott, J.E., Dorling, J., 1965. Differential staining of acid glycosaminoglycans (mucopolysaccharides) by Alcian blue in salt solutions. Histochemie 5, 221–33.

Shiraishi, F., Eno, Y., Nakamura, Y., Hanzawa, Y., Asada, J., Bahniuk, A.M., 2019. Relative influence of biotic and abiotic processes on travertine fabrics, Satono-yu hot spring, Japan. Sedimentology 66, 459–479. DOI: 10.1111/sed.12482

Shtukenberg, A.G., Punin, Y.O., Gunn, E., Kahr, B., 2011. Spherulites. Chemical Reviews 112, 1805–1838. DOI: 10.1021/cr200297f

Smythe, W.F., McAllister, S.M., Hager, K.W., Hager, K.R., Tebo, B.M., Moyer, C.L., 2016. Silica biomineralization of *Calothrix*-dominated biofacies from Queen’s Laundry Hot-Spring, Yellowstone National Park, USA. Frontiers Environ. Sci. 4, article 40, 1–11. DOI: 10.3389/fenvs.2016.00040

Sompong, U., Hawkins, P.R., Besley, C., Peerapornpisal, Y., 2005. The distribution of cyanobacteria across physical and chemical gradients in hot springs in northern Thailand. FEMS Microbiol. Ecol. 52, 365–376. DOI: 10.1016/j.femsec.2004.12.007

Sugihara, C., Yanagawa, K., Okumura, T., Takashima, C., Harijoko, A., Kano, A., 2016. Transition of microbiological and sedimentological features associated with the geochemical gradient in a travertine mound in northern Sumatra, Indonesia. Sedimentary Geology 343, 85–98. DOI: 10.1016/j.sedgeo.2016.07.012

Tobler, D.J., Blanco, J.R., Dideriksen, K., Sand, K.K., Bovet, N., Benning, L.G., Stipp, S.L.S., 2014. The effect of aspartic acid and glycine on amorphous calcium carbonate (ACC) structure, stability and crystallization. Procedia Earth and Planetary Science 10, 143–148. DOI: 10.1016/j.proeps.2014.08.047

Tong, H., Ma, W., Wang, L., Wan, P., Hu, J., Cao, L., 2004. Control over the crystal phase, shape, size and aggregation of calcium carbonate via a L-aspartic acid inducing process. Biomaterials 25, 3923–3929. DOI: 10.1016/j.biomaterials.2003.10.038

Tosca, N.J., Wright, V.P., 2018. Diagenetic pathways linked to labile Mg-clays in lacustrine carbonate reservoirs: a model for the origin of secondary porosity in the Cretaceous pre-salt Barra Velha Formation, offshore Brazil, in Reservoir Quality of Clastic and Carbonate Rocks: Analysis, Modelling and Prediction. Armitage, P.J., Butcher, A.R., Churchill, J.M., Csoma, A.E., Hollis, C., Lander, R.H., Omma, J.E., Worden, R.H. (Eds.). Geol. Soc. London, Spec. Publ. 435, 33–46. DOI: 10.1144/SP435.1

Tracy, S.L., Williams, D.A., Jennings, H.M., 1998. The growth of calcite spherulites from solution: II. Kinetics of formation. J. Crystal Growth 193, 382–388. DOI: 10.1016/S0022-0248(98)00521-1

Trichet, J., Défarge, C., 1995. Non-biologically supported organomineralisation. Bull Inst Océanogr Monaco Spec Issue 14, 203–236.

Valeriani, F., Crognale, S., Protano, C., Gianfranceschi, G., Orsini, M., Vitali, M., Romano Spica, V., 2018. Metagenomic analysis of bacterial community in a travertine depositing hot spring. New Microbiol. 41, 126–135.

Van Kranendonk, M.J., 2006. Volcanic degassing, hydrothermal circulation and the flourishing of early life on Earth: A review of the evidence from c. 3490-3240 Ma rocks of the Pilbara Supergroup, Pilbara Craton, Western Australia. Earth-Science Reviews, 74(3-4), 197–240. DOI: 10.1016/j.earscirev.2005.09.005

Veysey, J., Fouke, B.W., Kandianis, M.T., Schickel, T.J., Johnson, R.W., Goldenfeld, N., 2008. Reconstruction of water temperature, pH, and flux of ancient hot springs from travertine depositional facies. Journal Sedimentary Research 78, 69–76. DOI: 10.2110/jsr.2008.013

Viehmann, S., Reitner, J., Tepe, N., Hohl, S.V., Van Kranendonk, M., Hofmann, T., Koeberl, C., Meister, P., 2020. Carbonates and cherts as archives of seawater chemistry and habitability on a carbonate platform 3.35 Ga ago: Insights from Sm/Nd dating and trace element analysis from the Strelley Pool Formation, Western Australia. Precambrian Research 344, 105742. DOI: 10.1016/j.precamres.2020.105742

Visscher, P.T., Reid, R.P., Bebout, B.M., 2000. Microscale observations of sulfate reduction: correlation of microbial activity with lithified micritic laminae in modern marine stromatolites. Geology 28(10), 919–922. DOI: 10.1130/0091-7613(2000)28<919:MOOSRC>2.0.CO;2

Ward, D.M., Ferris, M.J., Nold, S.C., Bateson, M.M., 1998. A natural view of microbial biodiversity within hot spring cyanobacterial mat communities. Microbiol. Mol. Biol. Rev. 62, 1353–1370. DOI: 10.1128/MMBR.62.4.1353-1370.1998

Westall, F., Steele, A., Toporski, J., Walsh, M., Allen, C., Guidry, S., McKay, D., Gibson, E., Chafetz, H., 2000. Polymeric substances and biofilms as biomarkers in terrestrial materials: implications for extraterrestrial samples. Journal of Geophysical Research: Planets 105(E10), 24511–24527. DOI: 10.1029/2000JE001250

White, N.C., Wood, D.G., Lee, M.C., 1989. Epithermal sinters of Paleozoic age in north Queensland, Australia. Geology 17, 718–722. DOI: 10.1130/0091-7613(1989)017<0718:ESOPAI>2.3.CO;2

Wingender, J., Neu, T.R., Flemming, H.C., Eds., 1999, Microbial Extracellular Polymeric Substances, Berlin, Springer-Verlag, 251 p.

Wright, V.P., Barnett, A., 2015. An abiotic model for the development of textures in some South Atlantic early Cretaceous lacustrine carbonates, in Bosence, D.W.J., Gibbons, K.A., Le Heron, D.P., Morgan, W.A., Pritchard, T., Vining, B.A. (Eds.), Microbial Carbonates in Space and Time: Implications for Global Exploration and Production. Geol. Soc. London, Spec. Publ. 418, 209–219. DOI: 10.1144/SP418.3

Zhang, C.L., Fouke, B.W., Bonheyo, G.T., Peacock, A.D., White, D.C., Huang, Y., Romanek, C.S., 2004. Lipid biomarkers and carbon-isotopes of modern travertine deposits (Yellowstone National Park, USA): implications for biogeochemical dynamics in hot-spring systems. Geochimica Cosmochimica Acta 68, 3157–3169. DOI: 10.1016/j.gca.2004.03.005

